# Molecular evolution guided functional analyses reveals Nucleobindin-1 as a canonical E-box binding protein promoting Epithelial-to-Mesenchymal transition (EMT)

**DOI:** 10.1101/566836

**Authors:** Sansrity Sinha, Siddhart Pattnaik, Gopala Krishna Aradhyam

## Abstract

Calcium binding proteins (CBPs) function in response to changes in intracellular calcium (Ca^2+^) levels by modulating intracellular signaling pathways. Calcium sensors, including Nucleobindins (Nucb1/2) undergo Ca^2+^-binding induced conformational changes and bind to target proteins. Nucleobindins possess additional uncharacterized domains including partly characterized EF-hands. We study the molecular evolution of Nucleobindins in eukaryotes emphasizing on the N-terminal DNA binding domain (DBD) that emerged as a result of domain insertion event in Nucb1/2 domain-scaffold in an ancestor to the opisthokonts. Our results from *in silico* analyses and functional assays revealed that DBD of Nucb1 binds to canonical E-box sequences and triggers cell epithelial-mesenchymal transition (EMT). Thus, post gene duplication, Nucb1 has emerged as unconventional Ca^2+^-binding transcriptional regulators that can induce EMT.

**Highlights:**

1. Nucleobindins emerged from prokaryotic EF-hand containing protein.
2. The DNA-binding domain was gained in these in opisthokonts.
3. Gene duplication in ancestor of euteleostomes, lead to emergence of Nucb1/2.
4. Nucb1 emerged as a canonical E-box binding protein promoting EMT

## 1. Introduction

Calcium binding proteins (CBPs) display calcium (Ca^2+^) dependent changes in their expression profiles. A class of CBPs, calcium sensors, binds to target proteins in a Ca^2+^-dependent manner and generate the corresponding physiological response by modulating appropriate cellular signaling pathways [1–3]. Since Ca^2+^ is a well-known secondary messenger, aberrations in Ca^2+^ homeostasis is often a suggestive of metabolic and/or physiological imbalance (including cell death) [3–5]. Most CBPs are multiple EF-hand domain containing proteins with EF-hands contributing to their functional versatility and/or Ca^2+^ binding abilities. For instance, among the four EF-hands of calmodulin, EF-hand IV is clearly distinguishable from other EF-hands by isoelectric point and EF-hand III by volume [6–8]. Interestingly, contrary to other CBPs, Nucleobindins (Nucbs) (nucleobindin1/2, Nucb1/2) possess multiple domains in addition to EF-hand domains. In general, both Nucb1 and Nucb2 (in-paralogs) contain DNA-binding domain (DBD) followed by two EF-hands, leucine zipper (LZ) domains and unstructured poly glutamine stretch (pQ). These well characterized Ca^2+^ sensor proteins modulate various signal transduction processes in a cell type specific manner [9, 10]. Historically, Nucb1 (also known as Calnuc) was identified as a 55KDa protein that was triggered in response to antigenic nucleosomal DNA in systemic lupus-prone mice. This protein was shown to bind to λ-phage DNA by means of crucial DBD and pQ [11]. Later, it was identified as a 63KDa protein that binds with Gαi protein modulating G-protein signaling pathway [9, 12]. Recently, overexpression of Nucb1 was correlated with increased lymph node metastasis and lower overall survival rates in patients with gastric adenocarcinoma [13]. Nucb2 has also been identified as a potential biomarker for breast and renal clear cell carcinomas [14–18]. These lines of evidences indicate the involvement Nucb1/2 in the cancer progression. These studies have hypothesized that Ca^2+^-associated deregulation of signal transduction pathways could be the putative underlying cause for cancer progression in patients exhibiting higher expression profiles of Nucb1/2. However, the mechanistic details underlying its role in cancer progression remains unexplored. Also, the presence of an uncharacterized DBD and physiological demand pertaining existence of two similar Nucleobindins remains unexplored.

A previous study with a limited dataset revealed that Nucleobindins share evolutionary origins with calmodulin and troponin-C family proteins, all originating as a result of gene duplication of a single EF-hand containing protein in prokaryotes [19]. Consistent with this study, we demonstrate that Nucb emerged from a single EF-hand containing CBP in the last eukaryotic common ancestor (LECA) using the presence and absence of genes, the conservation patterns of protein domains and gene structure across Nucb orthologs across the eukaryotes. Also, the divergence of Nucb orthologs in eukaryotes was limited to addition/deletion of domains in the terminals of the nearly conserved EF-hands. Next, specific sequence level changes in the DBD concomitant to the deletion of C-terminal unstructured regions in Nucb1 lead to emergence of functionally divergent, Nucb1 and Nucb2 in-paralogs in euteleostomes. Together, the results from evolutionary analyses, *in silico* analyses and functional characterization strongly support the notion that Nucb1 evolved as a canonical E-Box binding protein following the gene duplication event in the euteleostomes that can trigger EMT in cell line models. Collectively, we report the molecular evolution of Nucleobindins in eukaryotes and suggest a plausible physiological evidence for the neo-functionalization of Nucb1 post gene duplication.

## 2. Materials & Methods

### 2.1 Data retrieval, multiple sequence alignment and phylogenetic analyses

Data mining for full-length orthologous protein sequences was done using Human Nucb1 and Human Nucb2 sequences as queries using NCBI BLAST tool (BLASTp, BLASTn, and tBLASTn strategy) (http://blast.ncbi.nlm.nih.gov/Blast.cgi). The sequences were curated for the domain architecture using NCBI CDD (http://www.ncbi.nlm.nih.gov/Structure/cdd/wrpsb.cgi) and COILS (http://embnet.vital-it.ch/software/COILS_form.html). TargetP was used to determine the subcellular localization of the protein (http://www.cbs.dtu.dk/services/TargetP/) [20, 21]. Only full-length sequences were taken up for further studies and are enlisted in Table S1. The alignment was done using MAFFT v7.245 (G-INS-1 strategy using BLOSUM62 as scoring matrix, gap opening penalty of 1.53, alignment threshold: score = 39, E-value = 8e-11). Bioedit7.25software was used to edit the alignment removing the DBD across all sequences [22]. Phylogenetic analysis (using ML and Bayesian methods) was done after choosing the best fit model (LG + G) based on AIC criteria using MEGA6.0 [23, 24]. Trees were built using RaxML and Mr. Bayes 3.2 and were visualized using iTOL (http://itol.embl.de/) [25, 26]. In Bayesian analysis optimal tree topology and posterior probability (PP) values for the nodes, with 100,000,000 Markov Chain Monte Carlo generations using stop value of 0.01 and the burn-in value were determined graphically by removing trees before the plateau. DBDs and EF-hand regions corresponding to sequences of euteleostome Nucb1 and Nucb2 were examined for functional divergence (type I and type II) using DIVERGE 2.0 software [27].

### 2.2 Gene structure analysis

The exon-intron organization of all genes was determined using the Gene Structure Display Server (GSDS) software (http://gsds.cbi.pku.edu.cn/)[28] and residues corresponding to the splice sites are shown in in File.S1.

### 2.3 Principal Component Analysis

Principal component analyses (PCA) were performed [29] to correlate biophysical properties of various domains across the Nucb orthologs using XL-STAT MS-Excel (XLSTAT 2017: Data Analysis and Statistical Solution for Microsoft Excel. Addinsoft, Paris, France (2017)).

### 2.4 Cell culture, maintenance and transfections

HEK293T and MCF7 cells were obtained from NCCS, India; HeLa and DLD1 cells were received as a kind gift from Dr Nitish. R. Mahapatra and Dr Rama Shankar Verma (Department of Biotechnology, Indian Institute of Technology, Madras, India). Cells were cultured in DMEM media (10% FBS in 5% CO_2_ incubator). Transfection was done using Lipofectamine 3000 using standard manufacturer’s instructions.

### 2.5 DNA manipulations and cloning

Nucb1 was cloned between HindIII and SacII in pcDNA6bv5His to generate Nucb1-pcDNA. Cripto promoter cloned in pGL4.20 and pGL4.20 human Cripto promoter were gifted by Dr. David Salamon. pRLTK for luciferase assay was a gift from Dr. Nitish. R. Mahapatra.

### 2.6 Electrophoretic Mobility Shift Assay

Canonical E-box containing oligos were annealed as per standard instructions. For *in vitro* binding assays, purified Nucb1 protein (2μM) was incubated with oligos (canonical E-box containing oligos, E1, and non-E-box oligos: E2 and E3, details in Table.S4) in the presence of Binding buffer (1.2M Tris, pH 8.0, 10mM EDTA, 1% Triton X 100 and 10%v/v glycerol) at room temperature for 30mins following which the products were loaded on 7% DNA PAGE gel. The binding reaction was also performed in the presence of 10μM CaCl_2_ to check the effect of Ca^2+^ on DNA binding ability of Nucb1. The electrophoresis was performed at 8V per cm voltage and the gel was stained using ethidium bromide and Coomassie brilliant blue to visualize DNA-protein complex/DNA and protein, respectively.

The details of methods used for transcript and protein quantification from cell lysates and cell based assays (promoter luciferase assay, wound healing and proliferation assay) are described in supplementary text S1.

## 3. Results

### 3.1 Origin of Nucb family of proteins

Orthologs for Nucb were identified in bacteria suggesting prokaryotic origin of Nucbs. In the eukaryotes a single Nucb ortholog was identified in both unikonts (Amoebozoa, Fungi, Choanoflagellata, Protostomia and Invertebrata) and bikonts (Chromalveolata and Excavata). This suggests that the ancestral Nucb probably emerged in the ancestor to unikonts and bikont lineages (possibly in the LECA). Interestingly, in the euteleostomes, two copies of Nucb (in-paralogs, Nucb1 and Nucb2) were identified (Fig.1A). Paralogs of Nucb (Nucb1/2) were identified in Actinopterygii (Bony fishes) and tetrapods but a single ortholog was identified in coelacanth (Sarcopterygii). This suggests i) a recent gene duplication in an ancestor to euteleostomes before the divergence of actinopterygian and sacropterygian lineages leading to the origin of the in-paralogs (Nucb1 and Nucb2) and ii) Nucb2 was subsequently lost in the coelacanth lineage in Sarcopterygii. Sequence based homology searches in Plantae lineage [namely the Angiospermae (*Arabidopsis thaliana*), Chlorophyta (*Chlorella variabilis* and *Monoraphidium neglectum*) and red alga (*Chondrus crispus*)] picked up single EF-hand containing putative Nucb orthologs. However, these were highly divergent (E-value>0.05) from other Nucb orthologs in the dataset and hence were ignored for further analyses. However, Nucb orthologs were not identified from Glaucophyta lineage suggesting gene loss in this lineage.

**Fig.1:**
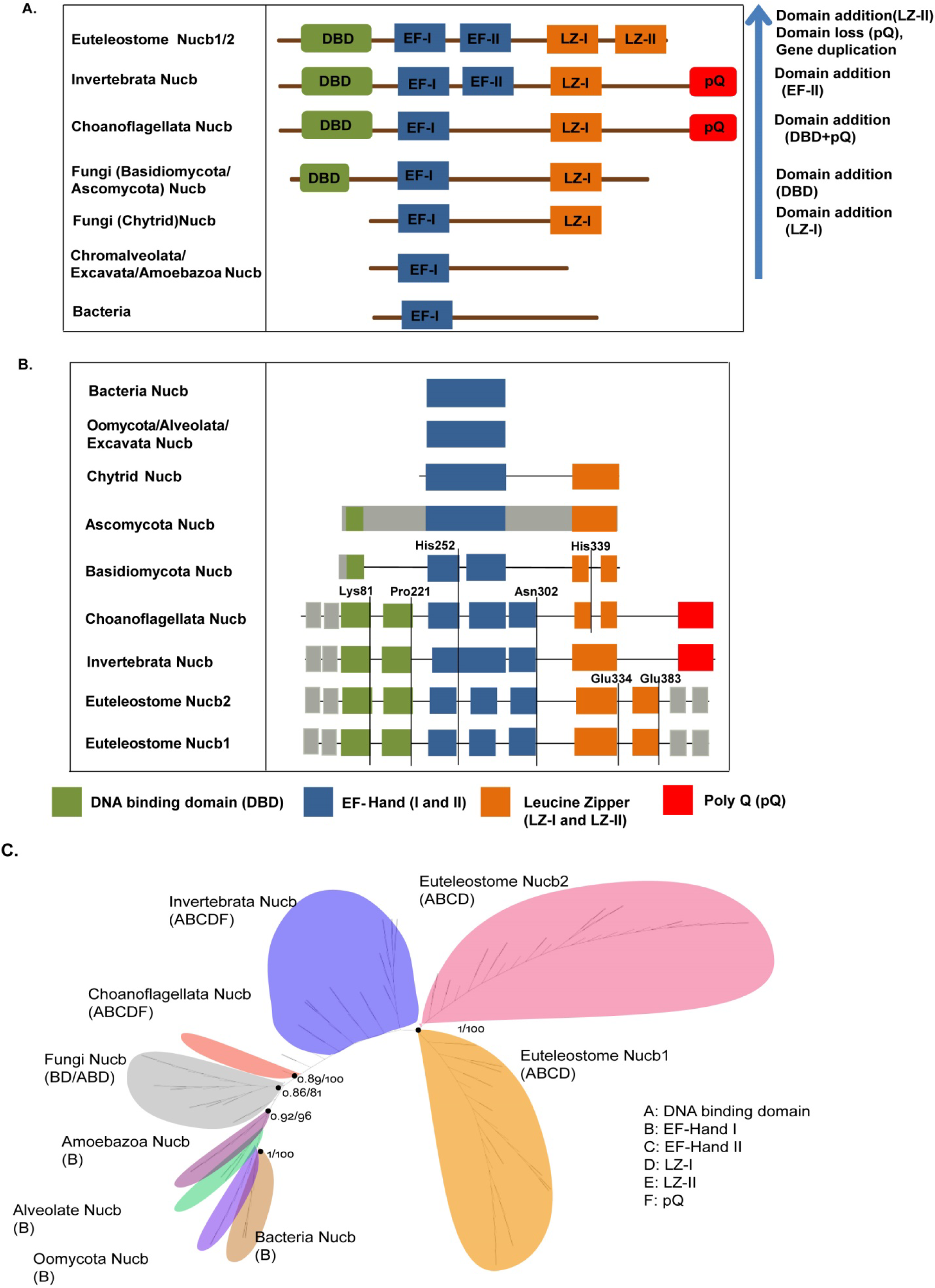
Evolution and diversification of Nucb proteins in eukaryotes. (A) Protein domain architecture diagrams for Nucb orthologs: The domains shown here, roughly representing the full-length protein sequence alignment. The lineages corresponding to the Nucb orthologs are shown in the column to the left. The direction of evolution is shown by the upward vertical arrow in the right. The crucial events (domain addition/deletion) in the evolution of Nucb orthologs in eukaryotes are marked on the right side of the table. The highly-divergent interdomain regions (between DBD, EF-hands, LZ and pQ) are represented by brown lines (not to scale) and do not provide any information concerning the sequence conservation in the alignment. (B) Gene structures of genes encoding for Nucb proteins in different phylogenetic groups: The lineages corresponding to the genes encoding the corresponding Nucb orthologs are shown in the column to the left. Intronic regions are represented as black lines. The exon-structures (limited to well-defined domains) are represented here roughly as per the full-length alignment. The gene structure features corresponding to interdomain regions are ignored. The color code followed for representing exons for various domains is shown at the bottom. The intradomain-shared splice site residues and their position are mentioned below the vertical blue bars. (C) Unrooted tree for 160 Nucb orthologous sequences using LG model with gamma distributed site variation up to 200bootstrap replicates. The numbers next to the black circles (marking the nodes) before and after “/” symbol correspond to the Bayesian and ML branch support values are mentioned. The tree was collapsed with a bootstrap cut-off of 70%. The orthologs of different lineages are marked in shaded colors along with the corresponding domain architecture. The alphabetical code representing the domains in the tree is shown in the key at the bottom of the tree.

### 3.2 Orthology of Nucb proteins

Eukaryotic multidomain proteins often undergo domain rearrangements, domain loss/gain events during the course of evolution leading to emergence of functionally divergent isoforms. In addition to protein domain architecture, conservation of gene structure in terms of exon number, splice site positions and intron loss-gain events provide crucial signatures for defining evolutionary relationships [30–33]. The conservation and divergence patterns in the domain architecture and gene structure were therefore analyzed for identified Nucb orthologs to decipher evolutionary relationships.

#### 3.2.1 Conservation of domain architecture

Nucb orthologs in the unikonts lineages (Amoebozoa and oomycota lineage of Fungi) and in the bikonts lineages (namely chromalveolates and excavates) were composed of a single EF-hand, similar to bacterial Nucb orthologs suggesting ancestral Nucb in the LECA probably possesed a single EF-hand. This is consistent with an earlier phylogenetic analysis stating an ancestral CBP in prokaryotes diversified to give rise to three distinct calcium binding proteins (namely troponin-C, calmodulin and Nucb) [19]. While all the bikont Nucb orthologs comprised only of a single EF-hand, the multi-domain architecture was observed specifically in the opisthokont lineage of the unikonts. The first multidomain Nucb, consisting of the N-terminal putative DBD (in Ascomycota lineage) followed by EF-hand and LZ, occurred in specific fungal lineage (Ascomycota and Basidiomycota) (Fig.1A). However, the Nucb orthologs in Ascomycota possess an uncharacterized truncated (or shorter) DBD. Nucb orthologs in the Urmetazoa lineage of unikonts resemble the fungal Nucb orthologs in terms of domain architecture albeit a longer DNA binding domain and an additional C-terminal pQ domain (Fig.1A and Fig.2A). The presence of pQ indicates a plausible domain addition event in choanoflagellates Nucb orthologs. Moreover, another EF-hand domain (non-canonical EF-hand, EF-hand II) was added in the Nucb orthologs (possibly as a result of domain duplication. Furthermore, in the eutelostomean additional LZ domain (LZ-II)) was added to the Nucb orthologs post gene duplication. The presence of EF-hand II in addition to EF-hand I in Urmetazoan Nucb orthologs is consistent with the classical EF-hand duplication model proposed for evolution of CBPs [8, 34].

**Fig.2:**
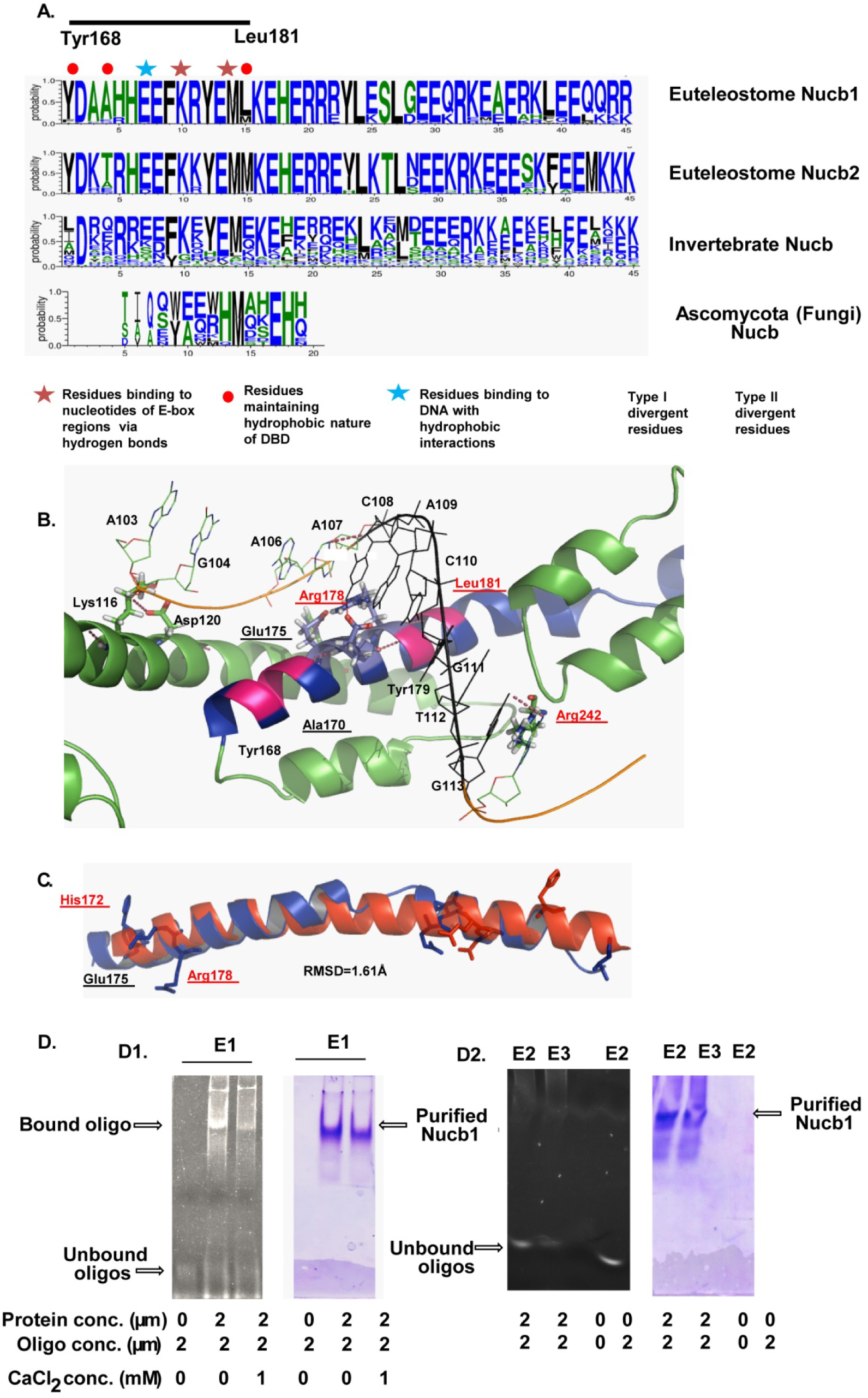
Characterization of DNA binding domain (DBD) of Nucb orthologs. (A) Weblogos displaying sequence consensus in Euteleostome Nucb1/2, Invertebrate and Fungal Nucb. The Y-axis corresponds to the probability of occurrence of amino acid at that position which the X-axis represents the position of those in the DBD of Nucb orthologs. The shared region of the DBD of Opisthokont Nucb and Myc is marked with dark horizontal black line (Tyr168-Leu18) and represents the minimal DBD. The key for all symbols mentioned on the top of Nucb Weblogos is mentioned below the figure panel A. Owing to shared features of Euteleostome Nucb1 with Myc, the DBD of euteleostome Nucb1 was predicted as a putative E-box (consensus 5′-CANNTG-3′) binding protein. (B) Cartoon representing the structure of Nucb1-DNA complex is shown as predicted by COACH server. The hydrogen bond interactions between amino acid residue of Nucb and nucleotide of DNA are shown by red dotted lines. The amino acid residues binding to DNA are shown in sticks. The residues showing functional divergence are underlined (Black font and red font used for type I and type II functional divergence, respectively). The DBD is shown in blue color while the regions flanking to it are shown with green in the cartoon representation of Nucb1-DNA complex. (C) Structural superposition of DBDs of Nucb1 (shown in blue) and Myc (PDB:5I50, shown in Red) (D) EMSA results demonstrating E-box binding specificity of Nucb1. The ethidium bromide (left) and commassie brilliant blue (right) stained gel images are shown next to each other for visualization of DNA + DNA-protein complex and protein, respectively. Native PAGE images showing binding of purified Nucb1 (D1) with canonical E-box [E1] oligos (2μM) (bands corresponding to bound oligo/shift) with and without 1mM CaCl_2_; and (D2) Non-E-box oligos [E2 and E3, 2μM each], respectively.

The pQ region was absent in euteleostome Nucb1/2 orthologs, suggesting domain loss subsequent to the gene duplication event in an ancestor to euteleostomes (Fig.1A). Interestingly, an extended non-homologous pQ region was identified only in Nucb1 orthologs corresponding to primates suggesting lineage specific feature of primates. The functional implication of this extended region remains to be understood. This is also evident by the presence of co-evolving groups in across Nucb1/2 sequences. The presence of residues across DBD, EF-hands and LZ domains in multiple co-evolving groups suggest that these domains have indeed co-evolved since their first emergence in Nucb proteins in eukaryotes (File.S3).

#### 3.2.2 Conservation of intron/exon number and intradomain splice site positions

The genes encoding Nucb orthologs in specific lineages in both unikonts (namely the Ascomycota class of fungi) and bikonts (chromoalveolates and excavates) were found to be single exon genes (SEGs). Introns were first found in specific lineages of Fungi (Chytrids and basidiomycota). Intron number was found to increase from Fungi to Urmetazoa suggesting that the molecular evolution of these proteins follows the ‘intron-late hypothesis’ [35, 36].

The splice sites corresponding to exon boundaries and interdomain positions were significantly conserved across genes encoding Nucb homologs. The splice site position corresponding to DBD displayed two patterns. In the first case, the interdomain orthologous splice site marking within the DBD (Lys81 of human Nucb1) was identified across Nucb orthologs from Fungi and Choanoflagellata while in the second, the splice site marking the end of DBD (Pro221 of Human Nucb1) was identified across the Nucb orthologs from Choanoflagellata (Ile) and Metazoa. The terminal splice site corresponding to EF-hand I was conserved (occurring ±5 positions from the reference residue) across Nucb orthologs from Fungi, Choanoflagellata (Thr) and Metazoa (Glu271 of human Nucb1) while that corresponding to EF-hand II was conserved across metazoan Nucb orthologs (Asn302 of Human Nucb1). Interestingly, the interdomain splice site corresponding to EF-hand I (His252, human Nucb1) was shared across fungi, choanoflagellates and euteleostome Nucb orthologs. Similar grouping was observed for splice site positions corresponding to LZ, wherein first group consisted of choanoflagellate and fungal Nucb orthologs sharing LZ-I terminal splice site (His339 of human Nucb1). The other comprised to euteleostome Nucb orthologs wherein (Glu359 of human Nucb1) that contained shared splice site corresponding to the terminal of LZ-II. Conservation patterns of the intradomain gene signatures in terms of shared orthologous splice site position, corresponding residues and exon numbers corresponding to each domain across the euteleostomes suggests the evolution of the in-paralogs Nucb1 and Nucb2 under strong negative purifying selection pressure (Fig.1B, File S1). Given the variation in the length of pQ across orthologs of Nucb from Metazoa, the corresponding orthologous splice site position was not identified. Together, the above results enable us to hypothesize orthologous relationship of Nucb proteins identified here.

### 3.3 Phylogenetic analyses of Nucb orthologs

In the light of above hypothesis, phylogenetic analysis was performed using full-length protein sequence alignment. The observed tree for Nucb orthologs was paraphyletic displaying three well supported clusters *viz;* (i) Nucb orthologs identified from bacteria and basal eukaryotes (the chromalveolates, oomycetes and the excavates), (ii) fungal Nucb orthologs and (iii) Urmetazoa Nucb orthologs (Fig.1C, File S2). The metazoan Nucb orthologs formed a monophyletic clade with a single bifurcation observed at a well-supported node separating the euteleostome Nucb1 and Nucb2 orthologs suggesting a recent gene duplication event in the ancestor to euteleostomes. The results of phylogeny support the hypothesis of orthologous relationship and common ancestry for Nucb orthologs (Fig1.C). The phylogenetic tree generated from the MSA lacking the DBD displayed star topology suggesting that the DBD carries sequence signatures largely contributing to evolution and diversification of Nucb orthologs. Since Nucb1/2 emerged from a recent gene duplication event, the in-paralog specific conserved and divergent patterns were analyzed further.

Principal component analyses (PCA) is a covariance analyses that is indicative of similarity and differences of a particular parameter across the samples. For a particular parameter being analyzed, the clustering of points corresponding to the samples in the dataset suggests similarity. On similar lines used previously, PCA was used to identify conservation and divergent patterns in the biophysical parameters of DBD and the loop regions between entry and exit helices of the two EF-hands across all Nucb ortholog examined here [6, 29]. The results from PCA for the loops of both EF-hands of the two in-paralogs suggest that these display similar flexibility profiles but differ in terms of hydrophobicity, isoelectric point and volume. The salient features of EF-hands and the loops across Nucb orthologs have been described in supplementary text S2.

### 3.4 In silico analyses suggest that Nucb1 may be a canonical E-Box binding protein

Results of PCA of the DBDs across Nucb orthologs revealed distinct biophysical attributes of fungal and metazoan DBDs (Fig.S3). The sequences corresponding to fungal and metazoan DBDs formed distinct clusters suggesting that these vary significantly in terms of all four biophysical parameters considered here. Distinct clusters were obtained in PCA for hydrophobicity for the euteleostome Nucb sequences (Nucb1 and Nucb2) suggesting the DBDs of Nucb1 and Nucb2 differ in hydrophobicity indices and confer functional divergence across the in-paralogs.

Interestingly the region of euteleostome DBD, corresponding to Tyr168-Leu181 of human Nucb1 was shared across the DBDs of all Opisthokont Nucb paralogs, suggesting that this region could be the putative core DBD present in these orthologs (Fig.2A). To gain functional insights to the observed differences, structural models of human Nucb1/2 were made (details in material and methods). The analyses of these models revealed that DBDs of Nucb1/2 are significantly similar (RMSD = 1.22Å). However, Nucb1 (COILS probability ∼ 0.9) possibly contains more structured and helical DBD as compared to that of Nucb2 (COILS probability ∼ 0.5).

The analyses by COACH server indicated that Nucb1 binds to E-box sequences of DNA suggesting it to be an E-box binding protein. Nucb1 was predicted to bind to 5′cytosine and 3′guanine of E-box sequences (consensus 5′-CANNTG-3′) of DNA via hydrogen bonds and hydrophobic interactions. The residues Tyr168, Ala170, Tyr179 and Leu181 were predicted to form the hydrophobic interface of DBD while Arg178 and Glu175 formed the charged surface on the DBD for interaction with DNA (Fig.2B). The structural comparison of DBD of Nucb1 and Myc (PDB:5I50) suggests significant similarity in the helical structure (RMSD=1.61Å) (Fig.2C). Both the structures also share residues that are crucial for Myc for binding to E-boxes sequences of DNA [37]. In addition to residues of DBD, sequences flanking the DBD (Lys116, Asp120 and Arg242) were also predicted to be involved in DNA binding. While Lys116 and Asp120 was found to be conserved across all euteleostome Nucb orthologs, Ala170, Glu175, Arg178 and Arg242 were found to be functionally divergent across Nucb1 and Nucb2 sequences (Fig.2B and Table 1). Given the fact that hydrophobic residues in the DBD of Nucb1 were substituted by charged residues in the DBD of Nucb2, it is plausible to assume that the hydrophobicity indices of the two DBDs may indeed be different as predicted by PCA (Fig.S3). Interestingly, the divergent residues were found to be fast evolving (MER∼2) than average (MER=1) in contrast to the conserved residues (MER∼0.2) (Fig.S1) suggesting that these sites may be evolving with differential evolutionary selection constraints.

**Table 1:** Type-II divergent amino acids identified from DBD of euteleostome Nucb1 and Nucb2 orthologs by Gu99 method.

| Type I divergence (Nucb1 vs. Nucb2) |  | Type II divergence (Nucb1 vs. Nucb2) |  |
| --- | --- | --- | --- |
| Z-score<br>(P value) | -2.867912 (P < 0.05) | $\alpha_{ML}$ | 0.401345 |
| $\theta_{I ML} \pm SE$ | 0.183200 ± 0.075261 | $\theta_{II} \pm SE$ | 0.019114 ± 0.059342 |
| LRT $\theta_I$ (P value) | 5.925283 (P < 0.05) | $G_R/G_C$ | 0.595238/0.016129 |
| Residues(PP) | R204 (0.474803) | N/C/R | 206/17/25 |
|  | Arg178 (0.418402) | F00,N/C/R | 0.524194/0.016129/0.018284 |
|  | Leu181 (0.474803) | Residues (PP) | A171K (4.973376) |

| Type I divergence (Nucb1 vs. Nucb2) |  | Type II divergence (Nucb1 vs. Nucb2) |  |
| --- | --- | --- | --- |
| Z-score<br>(P value) | -2.867912 (P < 0.05) | $\alpha_{ML}$ | 0.401345 |
| $\theta_{I}ML \pm SE$ | 0.183200± 0.075261 | $\theta_{II} \pm SE$ | 0.019114± 0.059342 |
| LRT $\theta_I$ (P value) | 5.925283 (P < 0.05) | $G_R/G_C$ | 0.595238/0.016129 |
| Residues(PP) | R204 (0.474803) | N/C/R | 206/17/25 |
|  | Arg178 (0.418402) | F00,N/C/R | 0.524194/0.016129/0.018284 |
|  | Leu181 (0.474803) | Residues (PP) | A171K (4.973376) |

Such sequence/motif specificity for DNA binding was not predicted for Nucb2. This suggests that sequence specificity of DNA may not be required for Nucb2 to bind to DNA and Nucb2 may bind to DNA with conserved Lys116 and Asp120 residues. Taken together, these analyses revealed differences in DBDs of Nucb1 and Nucb2 and it enables us to hypothesize that DNA binding specificity probably confers functional divergence to Nucb1/2.

### 3.5 Nucb1 is a canonical E-box binding protein

To validate sequence-specific DNA-binding ability of Nucb1, *in vitro* binding and mobility shift assays were performed wherein specific E-box oligos (consensus sequence 5′-CANNTG-3′) were alowed to bind to pure protein. The DNA-protein complex was subjected to native PAGE and the gel was stained with ethidium bromide and commassie brilliant blue to visualize DNA + DNA-protein complex and protein, respectively. The presence of band corresponding to DNA-protein complex and protein only in ethidium bromide and commassie brilliant blue stained gels, respectively, indicates binding of protein to DNA (oligos). The minimal concentration of DNA (oligo) and protein required to to visualize DNA-protein complex on ethidium bromide stained gel corresponds to the approximate K_d_ value of oligo for binding to protein. Nucb1 could bind to E-box oligo E1 (CACGTG) (K_d_ ∼2μM) and did not bind to E2 (CACGAG) and E3 (AACGTG) (Fig.2D) implying that Nucb1 is a canonical E-box binding protein that preferentially binds to specific E-box elements. These results suggest that substitution of 5′ cytosine by adenine (E3) and 3′ thymine by adenine (E2) probably abolishes the interaction of these bases of DNA to residues Arg178 and Arg242, respectively. In order to assess if the DNA binding ability of Nucb1 is dependent on its Ca^2+^ binding ability, *in silico* binding experiments were also performed in the presence of 1mM CaCl_2_, a source of Ca^2+^. The results revealed that the DNA binding ability of Nucb1 was affected by Ca^2+^ concentration, characterization of which needs further careful examination (Fig.2D1).

### 3.6 Underlying role of Nucb1 in cancer

E-Box binding proteins (for example: Myc, Snail etc.) are known to promote tumorigenesis and cell transforming abilities. Therefore, an analysis of expression profiles of Nucb1/2 from the TCGA datasets was performed (http://firebrowse.org/). Nucb1 was found to be upregulated in hepatic, head and neck and kidney cell carcinomas whereas it was found to be downregulated in cervical, ovarian, and colorectal carcinomas as compared to normal tissues respectively (Fig.S4A). There was no change in Nucb1 levels in Breast cancer, stomach cancer Uterine Corpus Endometrial Carcinomas. This indicates cancer type-specific deregulation of Nucb1. Hence, for further functional characterization of Nucb1, three cancer cell lines namely DLD1 (Colo-rectal cancer), HeLa (Cervical cancer), MCF7 (breast cancer) and a non-cancerous cell line HEK293T were considered. Since, Nucb1 was found to bind DNA *in vitro* and showed the presence of nuclear localization signal (NLS) in the N-terminal [predicted (NLS) from NES server ExPasy, Leu18-Ala23, Human Nucb1], we analyzed the cellular localization of Nucb1. Interestingly, Nucb1 localizes in the nucleus of the cells in addition to cytoplasmic and extracellular fractions (Fig.S4B), suggesting possible DNA binding function of Nucb1 *in vivo*. Together these results enables us to hypothesize that Nucb1 may bind to the E-box motifs of certain genes thereby modulating the expression of these genes and corresponding physiological response. Therefore, Nucb1 might have additional functions in the cells leading to cancer like phenotype apart from acting as a calcium sensor.

### 3.7 Nucb1 overexpression induces cell transformation properties and epithelial-to-mesenchymal transition (EMT)

To gain functional insights of Nucb1, cell transformation assays were performed in DLD1 and HeLa cell lines. The DLD1 and HeLa cells overexpressing Nucb1 displayed enhanced cellular proliferation (Fig.3A,3B and Fig.S6) and enhanced cellular migration (Fig.3C, 3D) rate indicating induction of EMT. This was demonstrated by characteristic changes in expression of known EMT markers (upregulation of Vimentin and Snail; and downregulation of E-cadherin) (Fig.4B-E). Only DLD1 cells overexpressing Nucb1 displayed spindle-shaped morphology characteristic of cells undergoing EMT (Fig.4A) indicating cell type specific feature of Nucb1. Thus, Nucb1 overexpression was correlated with EMT induction in DLD1 and HeLa cells suggesting that Nucb1 indeed plays a role in oncogenesis in these cell lines. Such effects were also observed for MCF7 cells (Fig.S6). In order to gain mechanistic insights regarding Nucb1 overexpression induced EMT, we analyzed the expression profiles of signal transducer molecules (AKT, MAPK). Interestingly, induction of phospho-AKT (pAKT) and its downstream target cyclin-D1 was observed in Nucb1 overexpressing HeLa cells (Fig.S7). Nucb1 overexpressing DLD1 cells displayed upregulation of VEGF at transcript and protein levels (Fig.4B,C), physiological relevance of which necessitates further characterization. Together, these results indicate that Nucb1 overexpression may trigger EMT. Incidentally, the results obtained here in our study for EMT induction by Nucb1 are consistent with those obtained for Nucb2 in colon cancer suggesting that similar to Nucb2, Nucb1 may also play a role in cancer progression [17].

**Fig.3:**
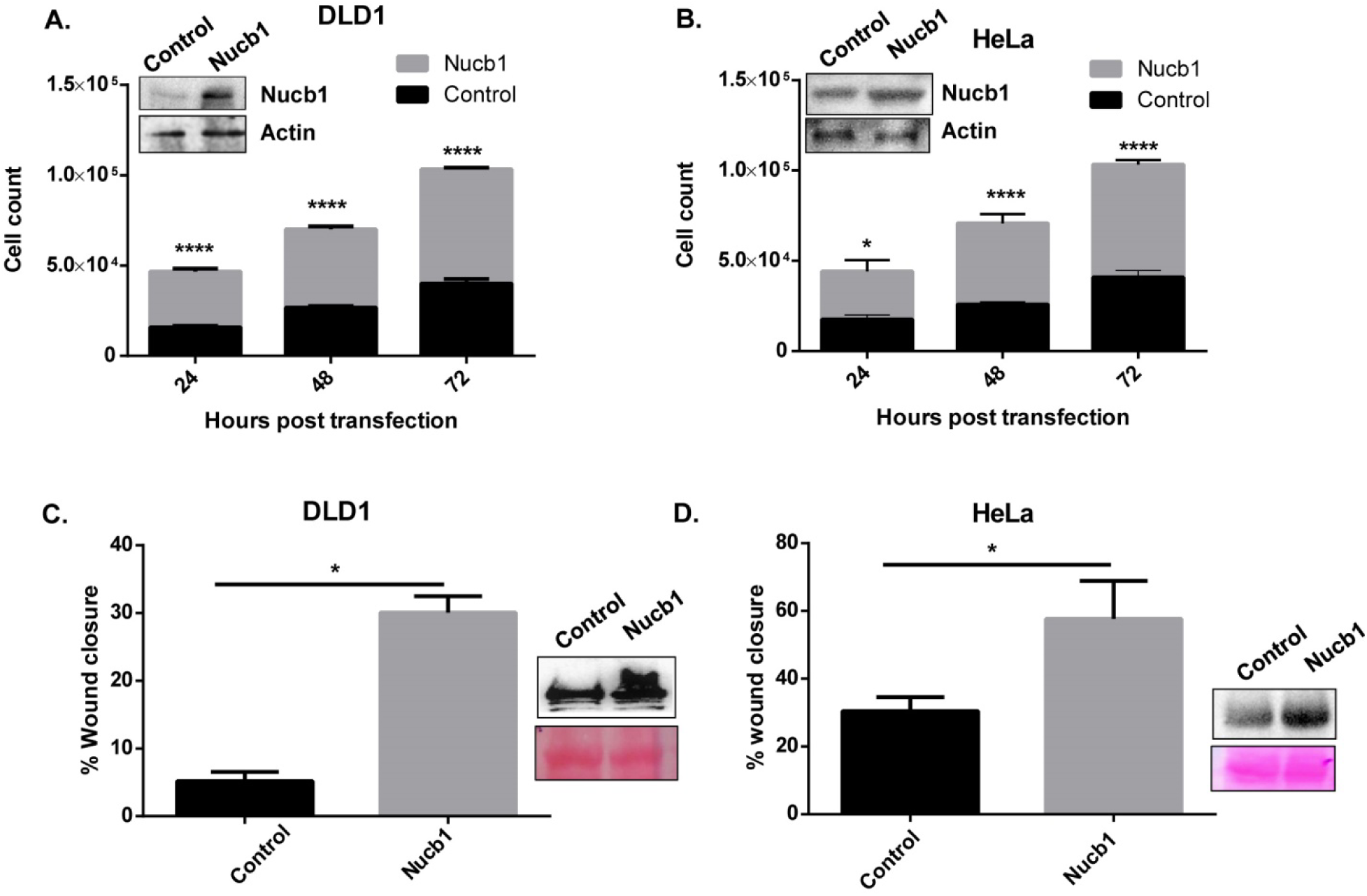
Nucb1 overexpression triggers cell proliferation and migration in DLD1 and HeLa cells. (A), (B) Bar graphs representing cell count measured in cell proliferation assay and western blot images of control and Nucb1 overexpressing DLD1 and HeLa cells, respectively (***P value<0.0001; ****P value P<0.00001; n=3, unpaired with Sidak’s multiple comparisions test). (C), (D) Bar graphs showing % wound closure in wound healing assay in Nucb1 overexpressing DLD1 and HeLa cells, respectively, at 0h and 24h with corresponding western blots showing Nucb1 overexpression. (*P value<0.05; n=3, unpaired with Sidak’s multiple comparisons test (alpha of 0.05). The ponceau images corresponding to the blot is also shown below the western blot for Nucb1.

**Fig.4:**
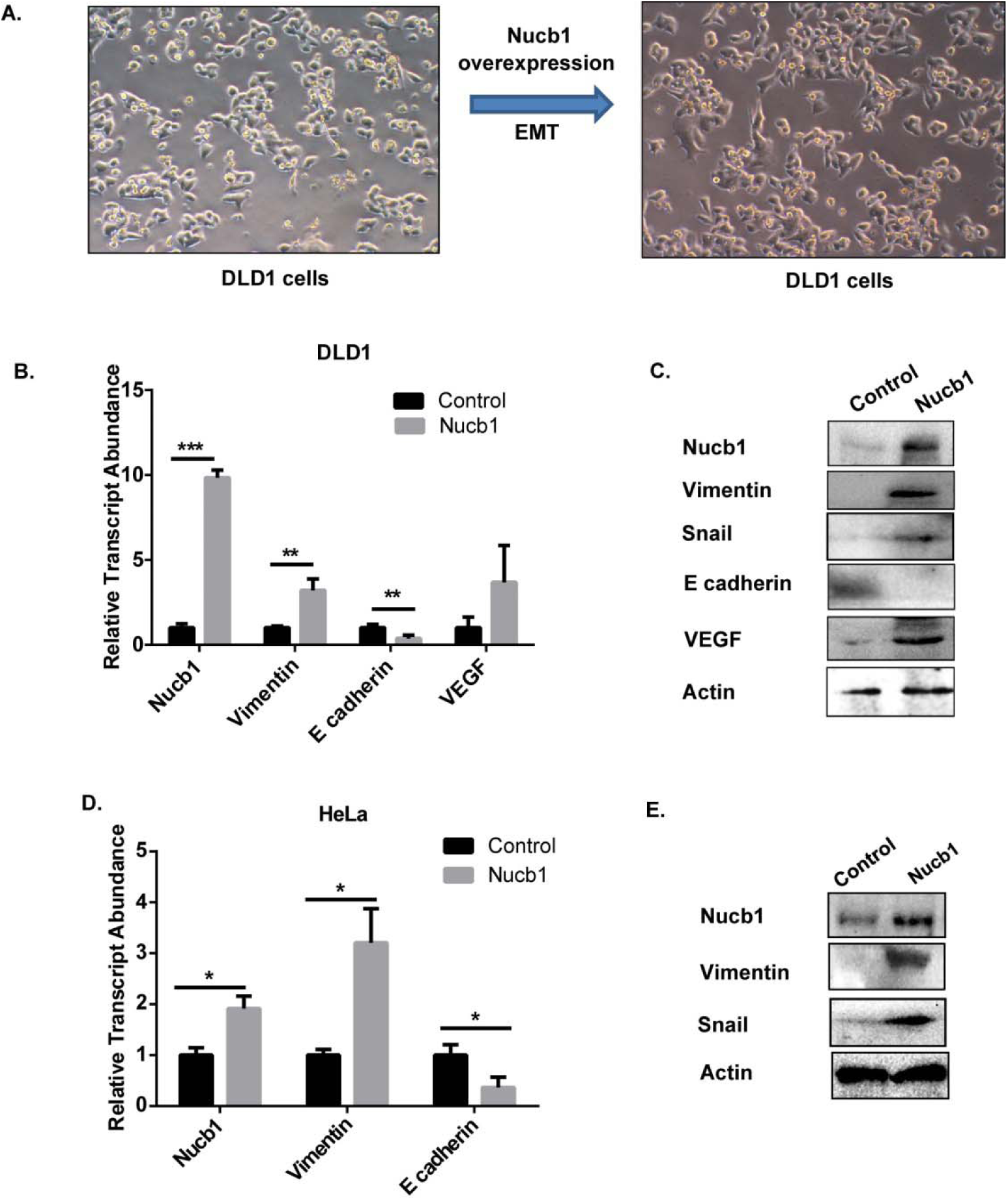
Nucb1 induces EMT in DLD1 and HeLa cells. (A) Changes in cellular morphology of DLD1 cells induced upon Nucb1 overexpression (20X magnification). (B), (D) Bar graphs showing enrichment of vimentin and snail, down regulation of E-cadherin by qRT-PCR (EMT markers) in Nucb1 overexpressing DLD1 and HeLa cells, respectively (***P value<0.0001; ****P value P<0.00001; n=3 two-way ANOVA with Sidak’s multiple comparisons test). (C), (E) Western blots showing enrichment of vimentin and snail, down regulation of E-cadherin (EMT marker proteins) in Nucb1 overexpressing DLD1 and HeLa cells respectively.

### 3.8 Nucb1 binds to an E-Box element on Cripto promoter to mediate EMT

Cripto is known to trigger EMT and tumorigenesis in MCF7 and IMR-32 cells [38, 39]. We observed an increase in the cripto transcript upon Nucb1 overexpression in DLD1 and HeLa cells (Fig.5B, D). Since we have shown that Nucb1 is an E-box binding protein and Presence of E-box elements was observed on cripto promoter also. So we hypothesized Nucb1 may regulate cripto expression by binding to its promoter regionNucb1 was found to bind to E-box (−1945 to −1939 from the transcription start site, TSS) present on Cripto promoter *in vivo* using ChIP-qRT-PCR in DLD1, HeLa and MCF7 cells (Fig.5A, C and Fig.S6.A). These results suggest that Nucb1 may bind to cripto promoter and transcriptionally regulate its expression as a putative transcription factor. This is further supported by luciferase assay where an increase in the luciferase activity of cripto promoter was observed in Nucb1 overexpressing cells (Fig.5E). A converse effect of Nucb1 binding to cripto promoter was observed in MCF7 cells and decreased luciferase activity was observed upon Nucb1 overexpression with no significant changes in the expression of Cripto transcripts (Fig.S6). These results suggest that the regulation of cripto by Nucb1 may be a cell type specific effect.

**Fig.5:**
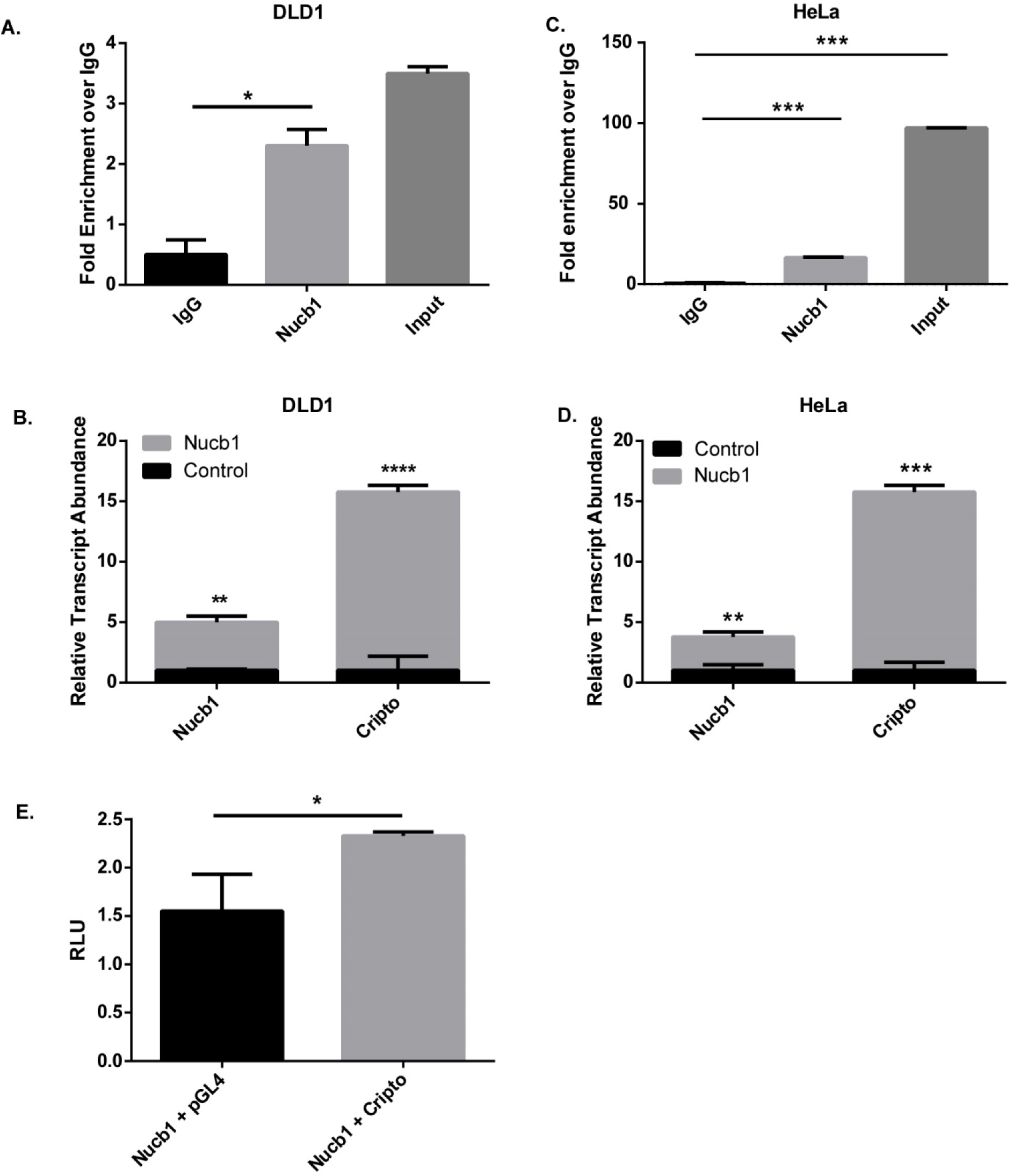
Nucb1 binds to Cripto promoter enhancing its transcription in DLD1 and HeLa cells. (A),(C) Bar graphs from ChIP qRT-PCR demonstrating Nucb1 binding to Cripto promoter in DLD1 and HeLa cells, respectively. IgG antibody was used a negative control and binding of Nucb1 to cripto promoter is shown by enrichment of transcripts corresponding to the region of binding of Nucb1 to Cripto promoter (***P value<0.0001; ****P value P<0.00001; two-way ANOVA with Sidak’s multiple comparisons test). (B), (D) Bar graphs (qRT-PCR) showing enrichment of Cripto transcript upon Nucb1 overexpression in DLD1 and HeLa cells, respectively. (E) Bar graphs from Luciferase assay demonstrating increase putative of Cripto promoter (represented by increase in RLU values) in Nucb1 overexpressing HeLa cells (***P value<0.0001; ****P value P<0.00001; n=3, two-way ANOVA with Sidak’s multiple comparisons test).

## 4. Discussion

Evolutionary analyses can provide insights into the functionality of multidomain proteins and have been used previously to gain functional insights for various multidomain proteins in eukaryotes [6, 8, 19, 40]. For instance, such approaches can explain the conservation and divergent patterns of various domains across the protein family and possibly explain the fate of duplicated genes and probable neo-functionalization. As opposed to other CBPs, Nucleobindins possess other uncharacterized domains (namely, the DBD, LZ and pQ) in addition to EF-hands. Therefore, in order to obtain insights about the conservation and divergence patterns of these domains we studied the molecular evolution of Nucleobindins. Our analysis shows that Nucb proteins in the eukaryotes probably occurred in LECA as CBP. It is evident that divergence of Nucb occurred mainly in the opisthokont lineage of the unikonts in following ways namely, (i) by insertion of specific domains in the terminal regions of EF-hands (DBD and LZ-I in an ancestor to opisthokonts), (ii) variation in intron phases of exons (iii) and by duplication of additional domains in specific lineages of opisthokonts (EF-hand duplication in Urmetazoa, pQ insertion in the ancestor to Metazoa and LZ duplication concomitant to pQ deletion in the ancestor to euteleostomes). Combining results from presence of genes encoding Nucb orthologs in eukaryotes, with the presence of shared features of protein domain architecture, gene structure and phylogeny we suggest that Nucb protein family formerly originated as a CBP in eukaryotes (Fig.6A). Furthermore, the absence of domain-shuffling events suggests that Nucb protein family has evolved under a strong negative selection pressure.

**Fig.6:**
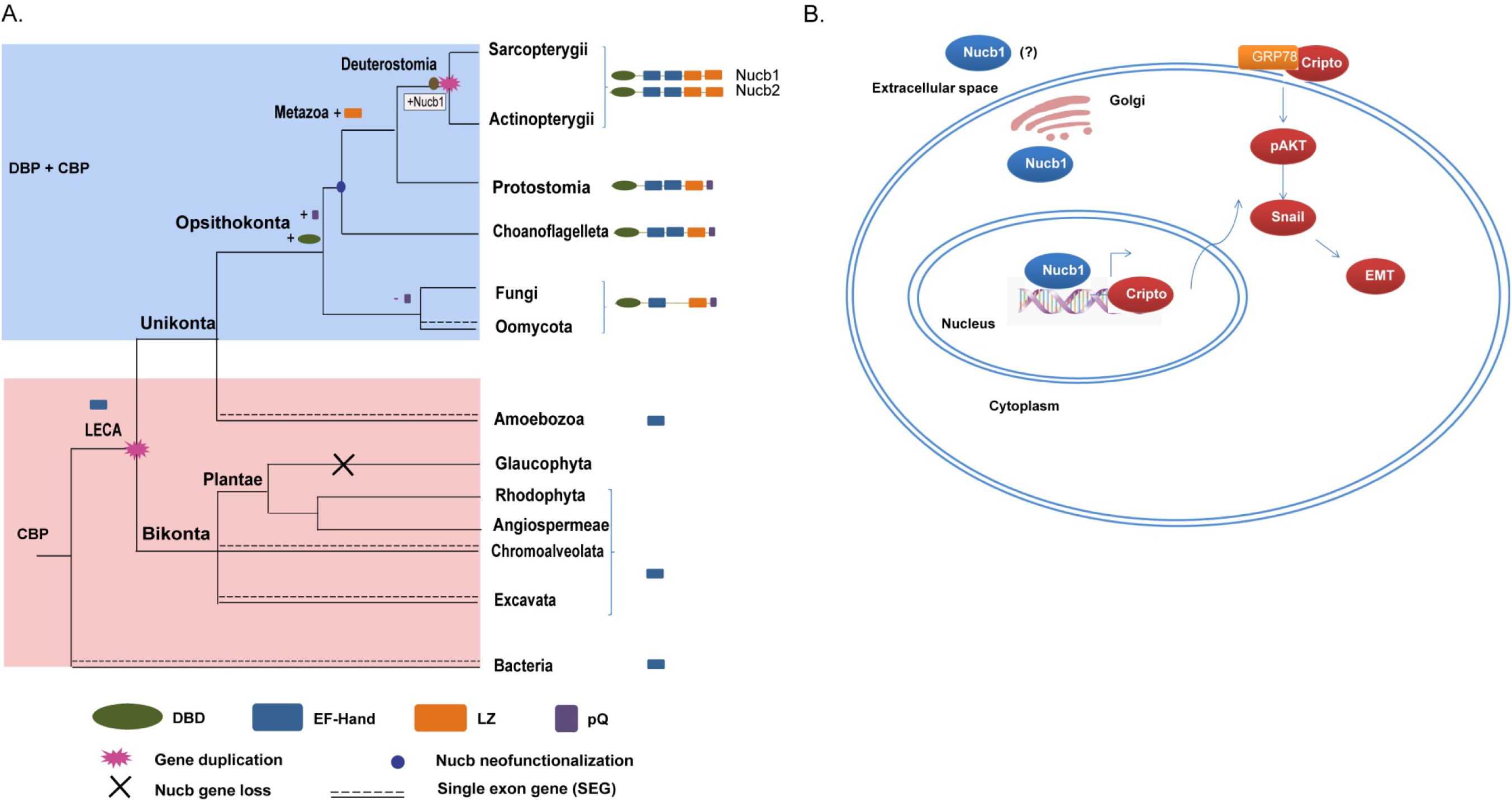
Molecular evolution of Nucleobindins and emergence of Nucb1 as a cannonical E-box binding protein that promotes EMT. (A) Diversification of Nucb proteins in eukaryotes. The presence of Nucb corresponding to each lineage is shown with the corresponding domain architecture. The domain addition events specific to each node is highlighted. Genes encoding Nucb orthologs were absent in the Glaucophyta lineage of the kingdom Plantae suggesting gene loss in this lineage. The branches with single line correspond to lineages that contain intron-containing genes that encode for Nucb orthologs. During the course of evolution Nucb1 evolved as a cannonical E-Box binding protein. The index for domains constituting the DSPs and for the symbols used for gene deletion, domain addition and specific neo-sub-functionalization events are shown in the index below. The lineages containing Nucb orthologs that function as calcium binding proteins (CBPs) are shaded with blue box while the DNA-cum calcium binding proteins (DNA + CBP) are shaded with pink box. (B) Characterization of Nucb1 classifies it as an E-box binding protein promoting EMT. The cartoon represents the cell showing the cell membrane bilayer, nucleus and Golgi compartments. Nucb1 is found in extracellular, nuclear and Golgi fractions. Within the nucleus Nucb1 interacts with E-box element present on cripto promoter inducing its expression. The EGF-CFC family protein cripto localizes with its interacting protein on the plasma membrane promotes EMT.

The Ca^2+^ binding properties of Nucb1 have been explored previously and the EF-hands involved in Ca^2+^ binding have been partly characterized. The solution NMR structure (PDB:1SNL) confirmed that all residues of both the EF-hands possess nearly identical Ф and L angles and bind Ca^2+^ with similar dissociation constants (47 and 40 µM for the non-canonical and the canonical EF-hand, respectively) indicating similar Ca^2+^ binding affinities [41]. This suggests that the presence of non-canonical EF-hand II does not contribute to differences in Ca^2+^ binding properties. Our results from PCA substantiate the above report showing that the loops of the two EF-hands are identical in flexibility (Fig.S2, supplementary text S1) Thus, it is evident that the presence of non-canonical EF-hand does not interfere with Ca^2+^ based conformational changes assumed by these proteins and hence these in-paralogs possess similar Ca^2+^ binding affinities. The loops of the two EF-hands across the in-paralogs however differ in the context of other biophysical properties which cannot be explained by limited dataset used here and need further examination. Moreover, we have predicted functionally divergent residues across EF-hand I and EF-hand II that putatively contribute to the non-canonical nature of the EF-hand II across the in-paralogs. However, the physiological relevance of the observed differences stated here remains to be deciphered.

Next, the presence of an uncharacterized DBD in a canonical CBP is suggestive of putative additional functions acquired by Nucb1/2 following the gene duplication event. In the light of increasing reports suggesting Nucb1/2 as a potential biomarker in cancer, our study was focused to functionally characterize Nucb1 taking cues from molecular evolution studies and *in silico* analyses [11, 13, 16, 18, 42–44].

The DBD has undergone substantial divergence since its emergence in the evolution of Nucb orthologs. The first short DBD appeared in an ancestor to the opisthokonts. Our results strongly suggest that the in-paralogs Nucb1 and Nucb2 differ broadly in the N-terminal regions. Thus, DBD was theoretically characterized in this study. Structural analyses of DBD of Nucb1 and its comparison with Myc suggest that the DBDs are significantly similar in these proteins. Moreover, only Nucb1 possesses crucial residues (Glu175 and Arg178 of human Nucb1) that enable its binding to the E-box sequences of DNA (Fig.2C) [37, 45–47]. Our results have identified divergent features of DBD across the in-paralogs suggesting that these may be functionally divergent (Table.2). In the light of experimental validations, it is clear that Nucb1 binds to canonical E-box sequences *in vitro* and *in vivo*. The presence of these divergent features in the DBD leads to a hypothesis that the divergence in the DBD may be the driving force for gene duplication and subsequent neo-sub-functionalization. Thus, in the context of recent gene duplication of Nucb in the ancestor to euteleostomes, the gain of function of DNA sequence-specific binding ability of Nucb1 suggests neo-sub-functionalization of Nucb1 [40, 48].

Substantiating previous reports, our results suggest that in addition to secretory fractions of the cell, Nucb1 was found in the nucleus and the cytoplasm. It is plausible to assume the differential localization of Nucb1 by virtue of its DNA and Ca^2+^ binding abilities. Nucb1 was found to be deregulated in various cancer types (Fig.S4) hence in order to functionally characterize Nucb1, cell transformation assays were performed. Nucb1 overexpression was found to promote cellular proliferation, migration and possibly EMT in DLD1, HeLa, MCF7 and HEK293T cells. Thus, it is plausible to assume that Nucb1 may play an important role in cellular transformation and possibly in cancer progression [16–18]. On further examination it was found that, Nucb1 binds to Cripto promoter *in vivo* in DLD1 and HeLa cells triggering the expression of Cripto. It is well known that Cripto triggers EMT via AKT/Snail axis [39]. Based on these observations we propose a putative mechanism of Nucb1 induced EMT induction, in Cripto dependent manner. Our results suggest that Nucb1 binds to E-box element on Cripto promoter triggering its expression. The EGF-CFC family protein, Cripto localizes on the plasma membrane and triggers downstream signaling to induce EMT via Nucb1/Cripto/AKT/Snail axis (Fig.6B).

Hence, Nucb1 can be considered a new addition to the category of unconventional Ca^2+^ binding transcriptional regulators as observed for FvCAMLP (*F. velutipes* calmodulin (CaM) like protein) that binds to non-canonical E-box elements on the promoter sequences of target genes thereby modulating their expression [49]. However, some questions still remains unanswered. It remains to be deciphered if the predicted crucial residues from our *in silico* analysis are indeed essential for the DNA binding and impart specificity for binding to E-box sequences on the DNA. Moreover, if Nucb1 is indeed a transcription factor/co-activator/repressor needs further characterization. Whether the secretory fraction or the DNA binding ability of Nucb1 is necessary for its role in EMT is yet to be explored. Also, if Nucb2 indeed needs sequence specificity for binding to DNA needs further experimental characterization.

## Supporting information

none

none

none

none

## Author Contributions

SS and AGK planned experiments; SS and SP performed the analyses; SS performed experiments; SS and SP analyzed data; AGK contributed reagents and essential materials; SS and AGK wrote the paper.

## Acknowledgements, Funding sources and disclosure of conflicts of interest

The authors acknowledge financial support from IIT Madras and infrastructure support from the Bioinformatics Infrastructure Facility and Central Equipment facility, IIT Madras. We also acknowledge Dr. D. Karunagaran, Dr. Nitish R. Mahapatra, Dr. Rama Shankar Verma and Dr. Suresh Kumar Rayala for cell lines and antibodies used in this study. We also acknowledge for Dr. D. Karunagaran and Dr. Suresh Kumar Rayala for useful scientific discussions and in interpretation of data. We acknowledge Mr. Vignesh Ravichandran for providing purified calnuc (Nucb1) protein used for *in vitro* binding studies. Authors declare no conflicts of interest.

## Abbreviations

Nucb: Nucleobindins
CBP: calcium binding protein
DBD: DNA binding domain
DBP+CBP: DNA: cum-calcium binding protein

