## Supplementary material for "Molecular evolution guided functional analyses reveals Nucleobindin-1 as a canonical E-box binding protein promoting Epithelial-to-Mesenchymal transition (EMT)": none

**Supplementary data:**

**Supplementary Text S1: Detailed Methods:**

**Data retrieval, annotation and determination of domain architecture**

Orthologs of Nucb1/2 were retrieved using human Nucb1 and Nucb2 as query sequences for BLAST search. Ssp120, an ortholog of Human Nucb1/2 in *Saccharomyces cerevisiae*, was used as a query sequence to identify orthologs in Fungi, Bacteria Chromoalveolata and Excavata. A single ortholog (named Nucb, Nb) was identified in protostomes, invertebrates, fungi, basal eukaryotes (Amoebozoa, chromalveolates and excavates) and bacteria. However, two copies namely, Nucb1 and Nucb2 were identified in the euteleostomes suggesting a gene duplication event in the ancestor to the euteleostomes. Sequences similar to Nucb1 and Nucb1 were annotated as Nb1 and Nb2, respectively. The identity of sequences comprising the final dataset of 160 predicted Nucb orthologs (including 36 previously unannotated sequences) is shown in Table.S1.

**Functional divergence Test:**

Pairwise coefficients (θ_ij_±SE) and likelihood ratio statistics (LRT) were performed for the euteleostome Nucb1, euteleostome Nucb2 and protostome Nucb orthologs using the Gu99 probabilistic method. A θ-I or θ-II significantly > 0 indicates site-specific altered selective constraints (type I) or a radical shift of amino acid physicochemical properties (type II) after gene duplication and/or speciation. When an amino acid is replaced at one particular position always by a particular another amino acid differeing in terms os biophysical properties, it is termed as type II divergence (site specific property shift) whereas when one amino acid is substituted at the same position by more than one type of amino acid (site specific rate shift), it is termed as type I divergence.

#### Mean evolutionary rate calculations and coevolution analyses

Mean (relative) evolutionary rates (MERs) for each position in the alignment was computed using MEGAv5.0 and the coevolving sites were identified using Coevolution Analysis of Protein Sequences (CAPS) server v2.0 with default parameters (Fares MA, 2006a; Fares MA, 2006b). The alignment file was submitted to the server and the output generated listed the coevolving sites in Nucb proteins for the particular phylogenetic group. The output for CAPS analyses is described in File S3.

**Principal Component Analysis:**

The PCA was performed for EF-hands domains and DBDs. Four biophysical parameters i.e. hydrophobicity, flexibility, isoelectric point and amino acid volume were taken into consideration. Hence four distinct datasets were obtained for each one of the three domains corresponding to the four biophysical parameters. The rows in these four datasets corresponded to the species from which the sequence was obtained and the columns corresponded to the amino acid positions in that particular domain. The data points corresponding to each dataset for a particular biophysical parameter was plotted in the space of first two principal components in the scatter plot. The principal component II (F2/PCII) in the ordinate was plotted against the principal component I (F1/PCI) in the abscissa to analyze the distribution of all the data points in the dataset. The contribution of each residue at a particular position for the specific biophysical parameter was is represented by the vector loadings (length of the arrows) in the corresponding biplots.

***Subcellular fractionation:***

Nuclear and cytoplasmic fractions of the cells were isolated as described previously(Albert S. Baldwin, 1996). Nuclear protein complex isolations buffer (20 mM Tris Cl, 420 mM NaCl, 1.5 mM MgCl2, 0.2 mM EDTA, 1 mM PMSF and 25% (v/v) glycerol, adjusted to pH 8.0) and cytoplasmic extraction buffer (10 mM HEPES, 60 mM KCl, 1 mM EDTA, 0.075% (v/v) NP40, 1mM DTT and 1 mM PMSF, adjusted to pH 7.6.)

#### Structural modeling, ligand prediction and structural superposition:

The human Nucb1 and human Nucb2 proteins was modeled using i-TASSER server (http://zhanglab.ccmb.med.umich.edu/I-TASSER/) (Roy et al., 2010; Yang et al., 2015; Zhang, 2008). The best fit model for both proteins was selected based on C-score and based on further examination of the stereo chemical parameters by SAVES server (SAVES v5.0) (https://www.acronymfinder.com/Structure-Analysis-and-Verification-Server-(University-of-California%2c-Los-Angeles)-(SAVES).html). COACH(Yang and Zhang, 2015) was used to predict ligands for the predicted structure and putative residues crucial for the ligand-protein interaction. FATCAT (http://fatcat.godziklab.org/) was used to superpose modeled structures of human Nucb1 and Nucb2 (Ye and Godzik, 2004).

#### Lysate preparation and western blot analyses:

Cells were lysed in RIPA buffer with protease inhibitor cocktail (Roche life science) to prepare lysates. The lysates were used to probe expression profile of proteins by western blot. 1:1000 dilution of the following primary antibodies in 3% BSA was used: Snail + Slug (Abcam), E-cadherin (Abcam), Vimentin (Cell signalling technology), Fibronectin (Abcam), Nucb1 (Santa Cruz Biotechnology), VEGF (Sigma Aldrich) and Actin (Santa Cruz Biotechnology). Following secondary antibodies tagged with horse radish peroxidase were used at a dilution of 1:5000 – anti-rabbit (Santa Cruz Biotechnology), anti-mouse (Santa Cruz Biotechnology).

#### RNA isolation, cDNA synthesis and qRT-PCR:

RNA isolation was done using RNAIso, Takara according to manufacturer’s intructions. 500ng of RNA was used for cDNA synthesis (cDNA reverse transcriptase kit, Applied Biosystems) which was later used for qRT-PCR analyses using Fast Sybr Green master mix (ROX) (Roche). The data from three independent experiments were used for analyses. The statistical difference of the relative transcript abundance of the specific target gene between control and Nucb1 overexpressing cells was computed two-way ANOVA with Sidak’s multiple comparisons test (alpha of 0.05).

#### ChIP and ChIP qRT-PCR:

Cells were grown to a confluence of 80% in 100mm tissue culture dishes and then cross-linked with 4% paraformaldehyde. Crosslinking, immunoprecipitation, washing and elution was performed as per manufacturer’s instructions. 1ug of Nucb1 antibody (Santa Cruz Biotechnology) was used for the immunoprecipitation. Purified immunoprecipitated DNA was used for qRT-PCR with the promoter primers of cripto. The statistical difference of the Nucb1 transcript abundance between control and Nucb1 overexpressing cells was computed using two-way ANOVA with Sidak’s multiple comparisons test (alpha of 0.05).

#### Promoter luciferase assay:

Promoter luciferase assay was done for cripto promoter. 10,000 cells were seeded in 12 wells plate. 24hrs later cells were co-transfected with pcDNA6b empty or pcDNA6b-Nucb1 expressing plasmid and pGL4.2 or pGL4.2-Cripto promoter and pRLTK plasmid (expressing Renilla luciferase) and incubated for 48 hrs. After incubation, cells were washed with PBS and lysed in 200µl passive lysis buffer. 50 µl of lysate was used for both firefly luciferase activity and Renilla luciferase activity respectively in luminescence plate reader. The luciferase activity was normalised with the Renilla luciferase reading. The statistical difference of the relative luciferase units (RLU) between control and Nucb1 overexpressing cells was computed two-way ANOVA with Sidak’s multiple comparisons test (alpha of 0.05).

#### Functional assays: wound healing assay and proliferation assay

For wound healing assay, 1 x 10^5^ cells were seeded per well in a 12 well culture plate and grown till confluence. After 24hrs (~75% confluence) wounding was done just before transfection using a sterile p20 pipette tip. The cells were cultured in serum free media to minimize proliferation. Images were collected using Olympus microscope using the cell^R software at 4X magnification every 24hrs post wounding for all cells until complete wound healing and the open wound area was calculated to determine % wound closure area. The difference in wound closure area between control and Nucb1 overexpressing cells was tested for statistical significance using two-way ANOVA with Sidak’s multiple comparisons test (alpha of 0.05). For proliferation time assay, 10000 cells were seeded per well in a 12 well culture plate and were incubated at 37^0^C, 5% CO_2_. After 24hrs transfection was done. Post-transfection after a regular period of 24hrs, cells from three wells were harvested and cell count was taken using trypan blue staining. This was done every 24hrs post transfection for 3days following which mean cell count per time point was plotted against hours post seeding for each cell line. The statistical difference in the number of cells for each time point between control and Nucb1 overexpressing cells was computed using two-way ANOVA with Sidak’s multiple comparisons test (alpha of 0.05). Images were taken using Olympus IX51 microscope 24hrs and 48hrs after transfection. Post-imaging cells were used for qRT-PCR and western blot analyses

#### Statistical Analysis:

All the data in result are analyzed quantitatively using GraphPad Prism6.0. Statistically significant data have been marked with asterisks. P-value ≤0.0001 ****, p-value ≤0.001 ***, p-value ≤0.01 ***, p-value ≤0.05 * represented in the graphs.

**Supplementary text S2: Salient features of EF-hands in Nucleobindins**

#### Helices of EF-hands of Nucb1 and Nucb2 possess functionally divergent sites

The sequences of both EF-hands were analyzed to identify functionally divergent residues by computing the coefficient of functional divergence (ϴ) across the in-paralogs. The value of ϴ and the posterior probability (PP) greater than zero signifies functional divergence at a particular site. Type II functionally divergent residues (Asn302Glu) and (Ser320Ala) were identified in the entry and exit helices of EF-hand II. A single type II divergent residue Asn244Asp (PP=4.97) was identified in the entry helix of EF-hand I (Table.S2). Incidentally, MER corresponding to these residues was higher (~2, marked with shaded blue box in Fig.S1C) suggesting probable acquiring of additional functions/properties and reduced evolutionary constraints for allowed substitution in both in-paralogs. We hypothesize that presence of functionally divergent sites in the entry and exit helices of the EF-hands might contribute to differences in biophysical properties of EF-hands between Nucb1 and Nucb2.

Principal Component Analyses (PCA) is a projection analyses that enables one to decipher in what respect one sample is different from another, which variables contribute most to this difference, and whether those variables contribute in the same way (i.e. are correlated) or independently from each other. On similar lines followed previously, each amino acid was assigned a numerical value for the biophysical pattern examined and analyses were performed (Bo Wang •, 2014; Halling et al., 2016). Based on our results for sequence analyses and domain architecture of recent duplication of EF-hands, PCA was used to identify conservation and divergent patterns in the biophysical parameters of DBD and the loop regions between entry and exit helices of the two EF-hands across all Nucb ortholog examined here.

#### EF Hands: Loops of EF-hand I and EF-hand II have distinct biophysical properties

In addition of sequence divergence in the entry and exit helices of EF-hand regions, the loop connecting the corresponding helices also contained variation in sequences between Nucb1 and Nucb2. The twelve residues in the loop regions between entry and exit helices of both EF-hands are known to be involved in Ca^2+^-binding (Alba and Tjandra, 2004; Grabarek, 2006; Lin et al., 2000; Maki et al., 2002). A comparative analysis for the differences in the biophysical parameters between the loop regions of EF-hand I and EF-hand II was observed by PCA. The loop regions of the two EF hands in Nucleobindins differ in their solvation energy or hydrophobicity, isoelectric point and volume (Fig.S2). The differences in these properties may be attributed to the observed sequence divergence in the 2^nd^, 4^th^ and 7^th^ residue of the loop region of EF-hands (shaded with vertical orange bars in Fig.S4). We hypothesize that the substitution of conserved hydrophobic Ile/Leu by polar Thr/Lys in 2^nd^ position and Ser by Glu/Lys in 4^th^ position and the presence of Gly substituting Arg in the 7^th^ position in EF-hand II classifies it as non-canonical EF-hand (as mentioned previously in (Alba and Tjandra, 2004)). Additionally, these substituted residues probably also contribute to the difference in observed differences in the profiles of hydrophobicity, isoelectric point and volume of the EF-hands in the PCA. However, the two EF-hands were found to be identical in terms of flexibility. Since Nucb1 and Nucb2 are calcium sensor proteins and undergo Ca^2+^ dependent conformational changes, hence as expected, the EF-hand loops responsible for Ca^2+^ binding display identical flexibility profiles therefore display nearly identical Ca^2+^-binding affinities. Thus, the results from PCA suggest that the EF-hand I and EF-hand II can be distinguished on the basis of the three biophysical parameters suggesting that evolution of these has been under strong purifying selection pressure (Fig.S2). Taken together, these observations highlight the amino acid residues in type II divergence tests could also be responsible for functional (and/or structural) compensations in spite of sequence level differences in EF-Hand I and EF-hand II (Fig.S1 and Fig.S5). This highlights the possible evolution directed sequence-based structural and functional compensations to both EF-hands for Ca^2+^ binding abilities. Our results highlight the need for further molecular characterization to decipher the physiological consequences of the predicted differences in the biophysical parameters in the context of Ca^2+^ binding ability and biophysical characteristics of both the EF-hands between the two in-paralogs.

**Table S2:** Type-I and Type-II divergent residues identified from loops regions of EF-hand I and EF-Hand II of euteleostome Nucb1 and Nucb2 orthologs by Gu99 method.

| **Type I divergence (Nucb1 vs. Nucb2)** | | **Type II divergence (Nucb1 vs. Nucb2)** | |
| --- | --- | --- | --- |
| Z-score (P value) | -2.867912 (P < 0.05) | α_ML_ | 0.401345 |
| θ_I_ML ± SE | 0.183200± 0.075261 | θ_II_ ± SE | 0.019114± 0.059342 (P < 0.05) |
| LRT θ_I_ (P value) | 5.925283 (P < 0.05) | G_R_/G_C_ | 0.595238/0.016129 |
| Residues(PP) | A319 (0.474803) | N/C/R | 206/17/25 |
|  |  | F00,N/C/R | 0.524194/0.016129/0.018284 |
|  |  | Residues (PP) | N244D (4.973376)  N302K (4.973376)  S320A (4.973376) |

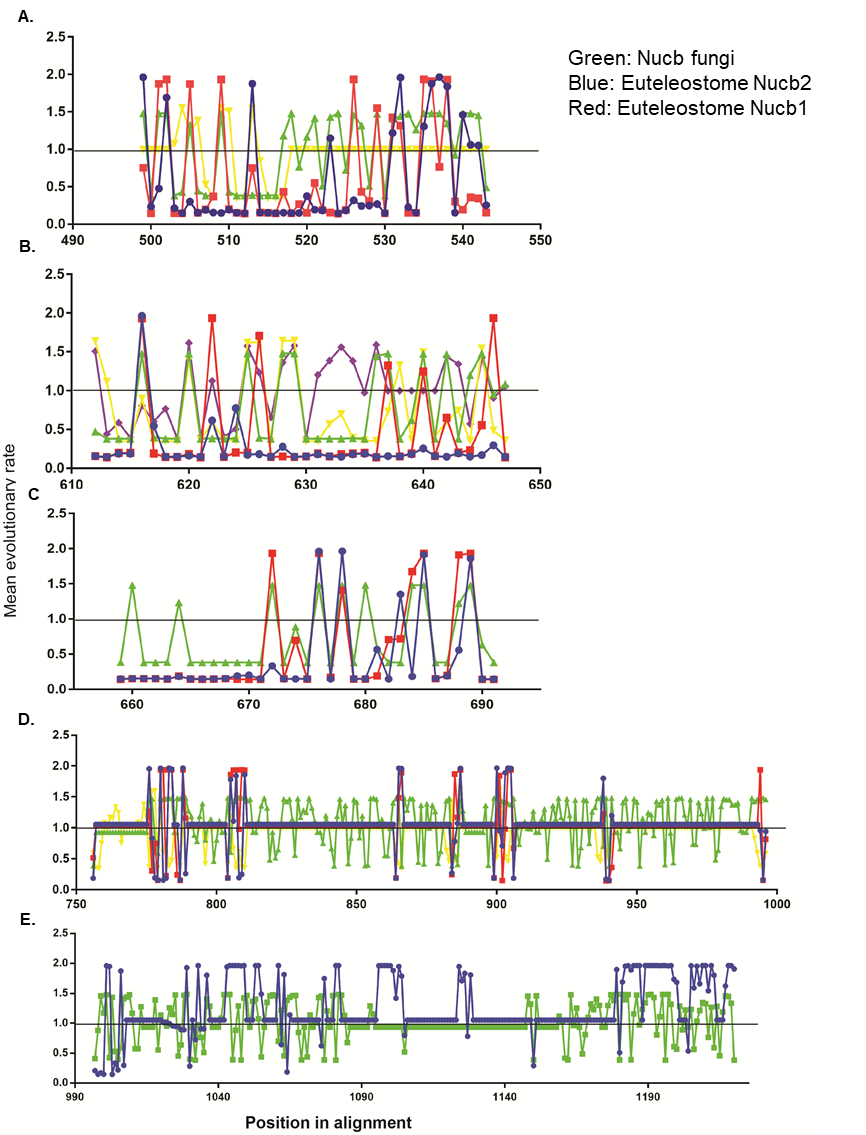

**Fig.S1:** **Domain-wise and phylogenetic group-wise plots of mean evolutionary rates (MERs) computed using MEGAv5.0 (JTT model, gamma distributed sites, 2 categories, max, Log likelihood = -64592.127).** These are shown for each site next to the site number. These rates are scaled such that the average evolutionary rate across all sites is 1. This means that sites evolving slower are showing a rate < 1 are evolving slower than average (MER=1), while those with rates higher than 1 are evolving faster than average. graph representing the computed MER for residues corresponding to (A) DNA binding domain, (B) EF-hand I (C) EF-hand II, (D) Leucine zipper domain. The X-axis represents the position in alignment and Y-axis represents the MER at each position. For all panels in this figure, blue line pertains to vertebrate Nucb1, red to vertebrate Nucb2, green to invertebrate Nucb, yellow to fungal Nucb and violet to bacterial Nucb sequences, respectively.

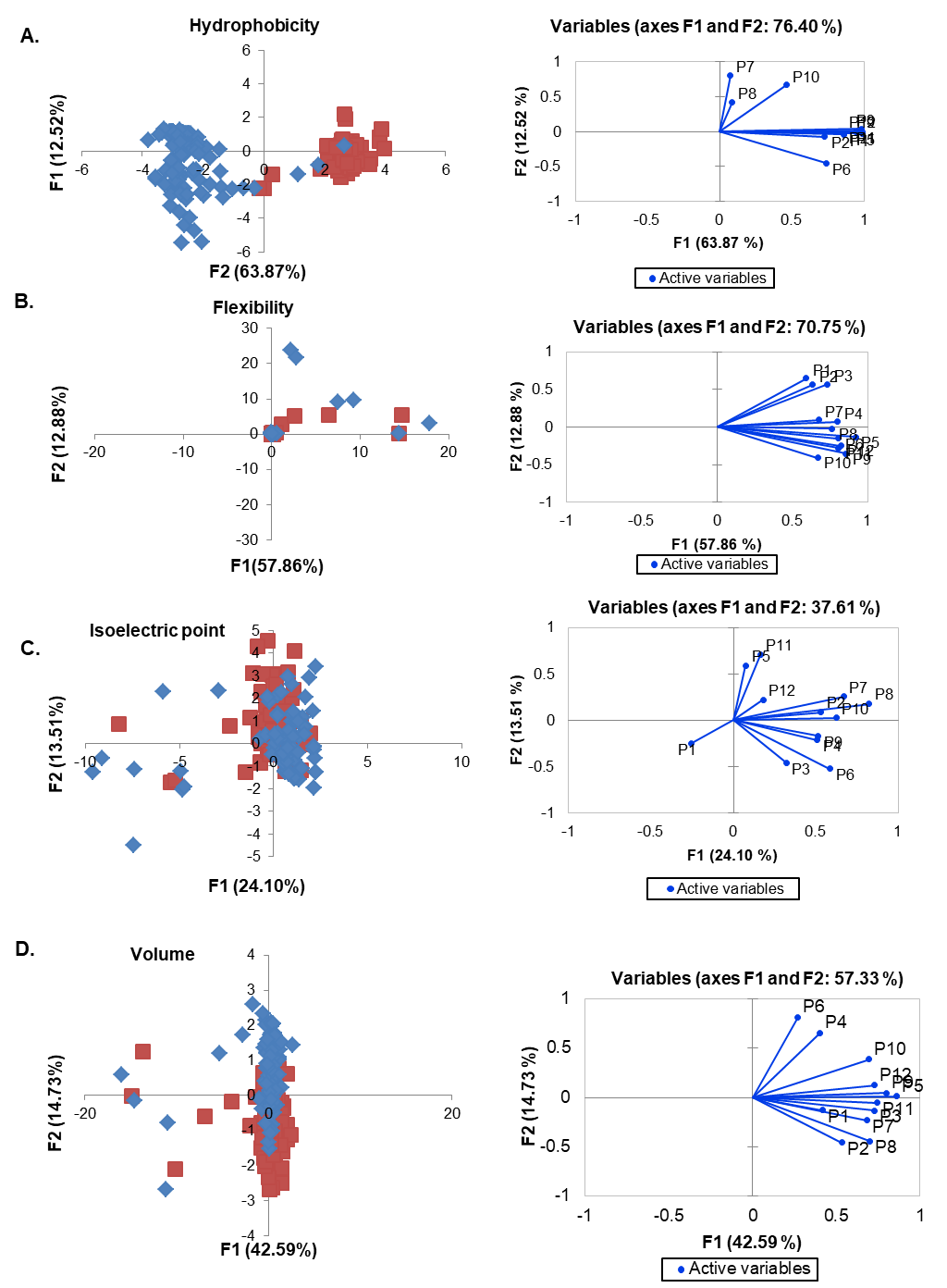

**Fig.S2: Principal Component Analysis of the EF loops**. Each EF-hand from each species is plotted in a plane with the first two principal components. Four different biophysical parameters are plotted from (A-D). Blue and red dots in the scatter plot correspond to F2 (PCII) for all data points for EF-hand II and EF-hand I respectively. The loadings for each position in the loop are plotted from. Loading vector for each position in the corresponding biplots shown next to the PCII scatter plot signifies its contribution to the variance of that specific biophysical parameter.

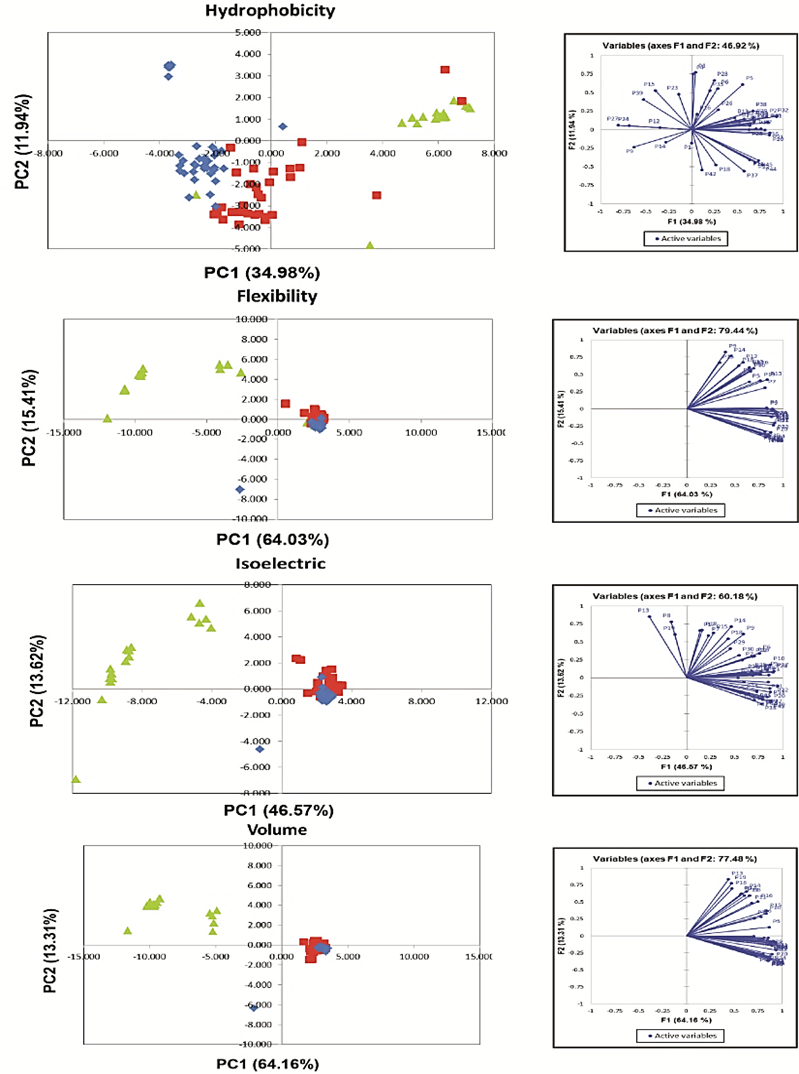

**Fig.S3:** **Biophysical and structural attributes of DNA binding domain of Nucb1/2.** (A) The scatter plot corresponding to the Principal Component Analysis (PCA) of the DNA binding domain for four different biophysical parameters is shown. Blue, red and green dots in the PCA scatter plot correspond to F2 (PCII) for all data points computed from sequences of euteleostome Nucb in-paralogs (Nucb1 and Nucb2), invertebrate Nucb and fungal Nucb paralogs. Loading vector in the corresponding biplots (shown next to the scatter plot) for each position signifies its contribution to the variance of that specific biophysical parameter.

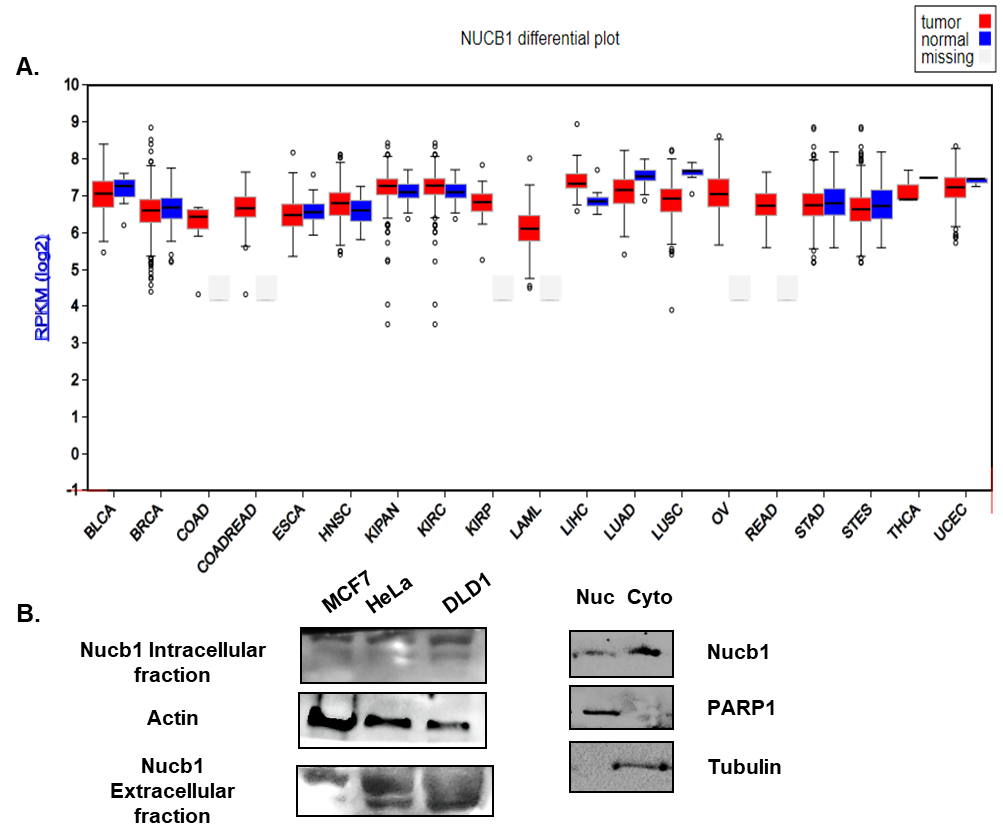

**Fig.S4:** **Nucb1 is differentially regulated in cancer types (TCGA dataset) and is present in multiple cellular compartments.** (A) Nucb1 is differentially regulated in various cancer subtypes as determined from TCGA dataset (). (B) Intracellular and extracellular levels of Nucb1 in MCF7, HeLa and DLD1 cells indicating Nucb1 as a secretory protein predominantly present in the secreted fraction. Western blot results demonstrating localization of Nucb1 in nuclear fractions [predicted nuclear localization signal (NLS) from NES server ExPasy, Leu18-Ala23, Human Nucb1] of HeLa cells.

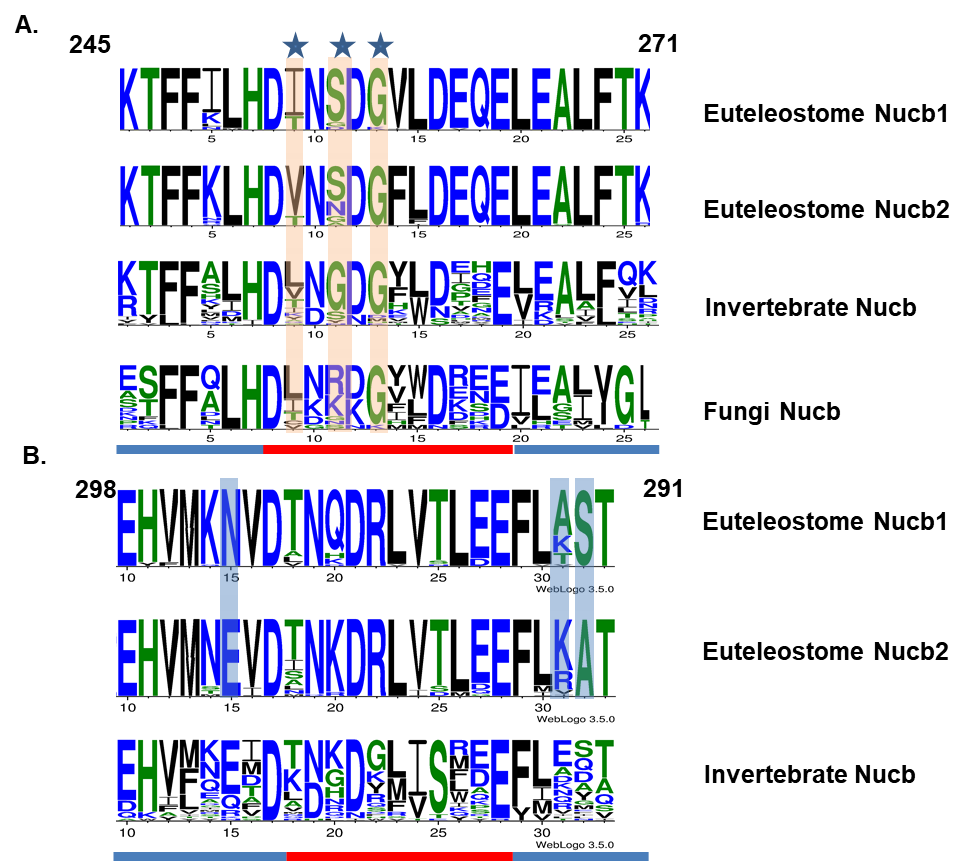

**Fig.S5: Conservation of residues in the helices and loop regions of EF-hands across metazoan Nucb orthologs.** The consensus sequences for (A) EF-hand I and (B) EF-hand II for Nucb orthologs. The loop region and the entry and exit helices are marked with red and blue horizontal thick lines below the Weblogos. The corresponding lineages for weblogos generated from its Nucb orthologs is mentioned in the right. The functionally divergent residues are highlighted with blue vertical boxes between euteleostome Nucb1 and Nucb2 orthologs. The divergent regions in the loop regions between EF-hand I and EF-hand II are shown by vertical orange shaded regions. The blue stars represent divergent residues in the loops of both EF-hands.

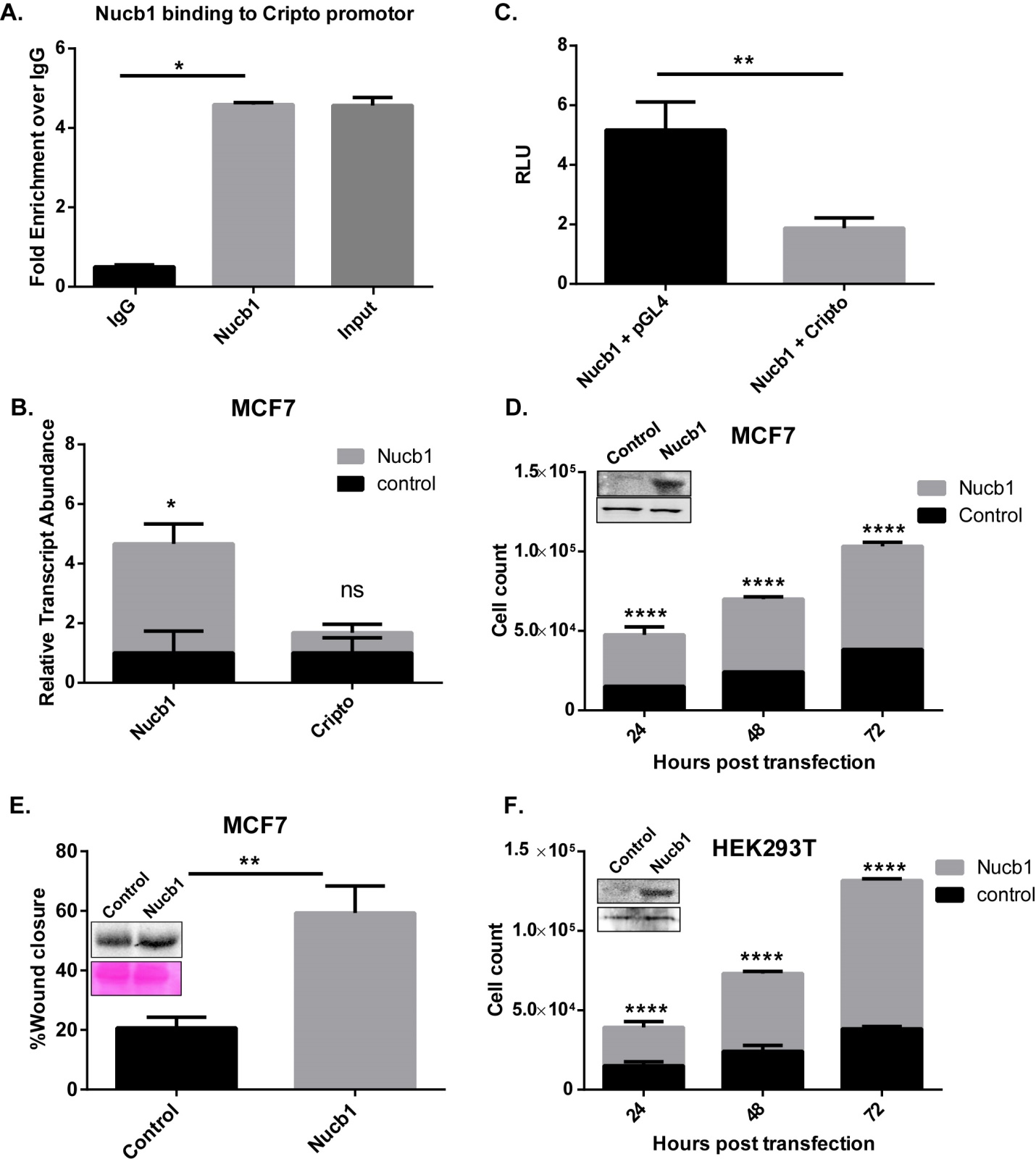

**Fig.S6:** **Characterization of Nucb1 in MCF7 and HEK293T cells.** (A) ChIP results demonstrating that Nucb1 binds to Cripto premotor in MCF7 cells. (B) qRT-PCR result showing that Nucb1 overexpression does not correlate with expression profile of Cripto. (C) Luciferase assay showing decrease in Cripto promoter activity upon Nucb1 overexpression (Holm’s Sidak comparison, n=4). Thus Nucb1 binds to cripto promoter and decreases its activity *in vitro* in MCF7 cells. However no effect was observed on cripto transcript expression following Nucb1 overexpression (as shown in (B)). (D) Proliferation assay result showing that Nucb1 overexpressing leads to enhanced proliferation in MCF7 cells. (E) Representative bar graphs showing %wound closure in the wound healing assay at 0 h and 24 h with corresponding western blots of enrichment of Nucb1 in extracellular fraction of the cell. (*P value<0.05; unpaired with Sidak’s multiple comparisons test (alpha of 0.05). The ponceau images corresponding to the blot is also shown below the western blot for Nucb1. (F) Bar graphs showing significant increase in cell count of Nucb1 overexpressing HEK293T cells Vs. control at corresponding time-points shown in the X-axis in the proliferation assay (***P value<0.0001; ****P value P<0.00001; Sidak’s multiple comparisions test). The corresponding western blot for 24hr post transfection is shown.

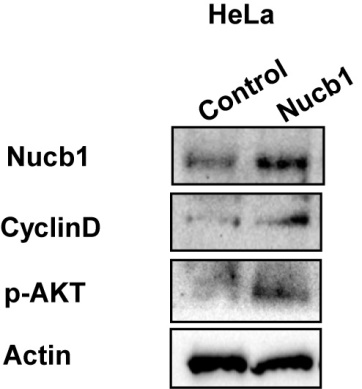

**Fig.S7:** **Nucb1 overexpression is associated enrichment of Cyclin-D1 and p-AKT in Nucb1 overexpressing HeLa cells.** Western blots showing enrichment of Cyclin-D1 and p-AKT in Nucb1 overexpressing HeLa cells.

**Table S1:** List of protein sequences taken up for the study

**Table S4:** List of oligo sequences used for binding assays in Fig.2.

**Supplementary files:**

Supplementary File S1: Full-length protein alignment with marked shared splice site positions (pdf).

Supplementary File S2: ML tree generated using the full-length protein alignment (newick)

Supplementary File S3: CAPS output excel file
