## Supplementary material for "Molecular evolution guided functional analyses reveals Nucleobindin-1 as a canonical E-box binding protein promoting Epithelial-to-Mesenchymal transition (EMT)": none

```
                                                   1                     10
HsapNb1   ........................................MPP.........SGPR....GTL.LLLP..
PtroNb1   ........................................MPP.........SGPR....GTL.LLLP..
PpanNb1   ........................................MPP.........SGPR....GTL.LLLP..
PabeNb1   ........................................MPP.........SGPQ....GTL.LLLP..
GgorNb1   ........................................MPP.........SGPR....GTL.LLLP..
CatyNb1   ........................................MPP.........SGPR....GTL.LLLP..
MmulNb1   ........................................MPP.........SGPR....GTL.LLLP..
RroxNb1   ........................................MPP.........SGPR....GTL.LLLP..
CangNb1   ........................................MPP.........SGPR....GTL.LLLP.L
MfasNb1   ........................................MPP.........SGPR....GAL.LLLP..
MnemNb1   ........................................MPP.........SGPR....GTL.LLLP..
CcapNb1   ........................................MPP.........SGPR....GTL.LLLS..
OgarNb1   ........................................MPP.........SGPR....GAL.LLSPLL
PcoqNb1   ........................................MPP.........SGSR....GAL.LLPPLL
GvarNb1   ........................................MPP.........CGPR....GAL.LFLPRP
TchiNb1   ........................................MPP.........SGPR....GSL.LLPP..
BmutNb1   ........................................MPP.........SGPR....AAL.FLLP..
MjavNb1   ........................................MPP.........SGPR....ATL.LLLP..
VpacNb1   ........................................MPP.........SGPR....AAL.LILP..
CbacNb1   ........................................MPP.........SGPW....AAL.LILP..
VpacNb1   ........................................MPP.........SGPR....AAL.LILP..
CferNb1   ........................................MPP.........SGPW....AAL.LILP..
HarmNb1   ........................................MPL.........SGPR....AAL.LLLP..
RsinNb1   ........................................MPP.........SGPT....AAL.LLLP..
JjacNb1   ........................................MPP.........SGSR....KTL.......
DordNb1   ........................................MPG.........SGPH....RAL.......
NgalNb1   ........................................MPP.........SAPR....TVL.CLTL..
RnorNb1   MKI...............EPAT..................MPT.........SVPR....GAPFLLLP..
MmusNb1   .KTPSAL..........ERPAT..................MPT.........SVPR....GAPFLLLP..
MaurNb1   ........................................MPT.........PVPR....GAP.LLLP..
CasiNb1   ........................................MSP.........TGPR....AAF.......
NparNb    ........................................M.......................KICY..
XlaeNb1   ........................................M.......................KICT..
XtroNb1   ........................................M.......................KICT..
PmucNb1   ........................................M.......................HLPW..
PbivNb1   ........................................M.......................HLLW..
GjapNb1   ........................................M.......................HLTW..
DrerNb1   ........................................M...........TGSF....KAL.......
TrubNb1   ........................................M...........KLRP....GSL.......
TnigNb1   ........................................M...........RLRP....GAL.......
IpunNb    ........................................M...........SWIK.....VL.TAC...
DrerNb2   ........................................M...........SCLK....GLL.TSC...
TrubNb2   ........................................M...........GWIR....TLL.TVW...
HsapNb2   ........................................M...........RWRT....ILL.QYCF..
PpanNb2   ......................................M.M...........RWRT....ILL.QYCF..
GgorNb2   ......................................M.M...........RWRT....ILL.QYCF..
CatyNb2   ..........................................................................
CangNb2   ..........................................................................
RroxNb2   ........................................M...........RWRT....VLL.QYCF..
PabeNb2   ........................................M...........RWRN....ILL.QYCF..
MmulNb2   ........................................M...........RWRT....ILL.QYCF..
MfasNb2   ........................................M...........RWRT....ILL.QYCF..
CcapNb2   ........................................M...........RWRT....LLL.QYCF..
PtroNb2   ........................................M...........RWRT....ILL.QYCF..
OgarNb2   ........................................M...........RWRT....ILL.QYCF..
PcoqNb2   ........................................M...........RWRT....ILL.QYCI..
TchiNb2   ........................................M...........RWRT....ILL.QYCF..
MjavNb2   ........................................M...........RWRT....ILL.QDCF..
HarmNb2   ........................................M...........RWRT....ILL.QYCF..
RsinNb2   ........................................M...........RWRT....ILL.QYCF..
DordNb2   ........................................M...........RWRT....ILL.QNCL..
BmutNb2   ........................................M...........KWRT....ILP.QCCL..
VpacNb2   ........................................M...........KWRT....ILL.QYCF..
CferNb2   ........................................M...........KWRT....ILL.QYCF..
CbacNb2   ........................................M...........KWRT....ILL.QYCF..
JjacNb2   ........................................M...........RWRT....IPP.QCCS..
CasiNb2   ........................................M...........RRRT....FLL.Q.CC..
NgalNb2   ........................................M...........WWRT....IQL.QHCF..
RnorNb2   ........................................M...........RWRT....IQA.RYCF..
MmusNb2   ........................................M...........RWRI....IQV.QYCF..
MaurNb2   ........................................M...........RWKI....IQL.QYCF..
PbivNb2   ........................................MDRCQ...RKQIRWRT....TLL.QQCV..
PmucNb2   ........................................MDPCR...RKQIRWRT....TLL.QQCV..
GjapNb2   ........................................MHRSQ...LKK.RWRT....GLL.QQCA..
LchaNb    ........................................M.....................L.HY....
XlaeNb2   ........................................M.......SSEVRWGR....IFV.ACSS..
XtroNb2   ........................................M....................SL.ICSS..
AcalNb    ........................................MG..........DYRLP...RLL......L
BglaNb    ..........................................................................
LgigNb    ..........................................................................
DmagNb    ........................................M............EGV....STF.LVAA..
DpulNb    ........................................M.............................
HrobNb    ........................................MDRPAISFPSLLYKKS....VLI.FFAV.A
HaztNb    ........................................M.............KF....SSL.SIAI..
TurtNb    ........................................M............MTL....SDI.YTCI..
PtepNb    ........................................M............WKF....FCI.......
SmimNb    ........................................M............RAL....ILT.......
GoccNb    ........................................M............KHL....GAL.QVAV..
DsuzNb    ........................................M...........AQNV....ALL.......
DmelNb    ........................................M...........VQNV....ALL.......
AaegNb    ........................................MYE.........SGKV....SLV.GV...F
CtelNb    ........................................M............RGV....AFL......A
OvicNb    ........................................M............AKW....HYL.......
SkowNb    ........................................M............ASW....KIL.......
SpurNb    ........................................M...................GML.EWKL.P
NvecNb    ........................................M...........DSKL....SLL......S
EpalNb    ........................................M...........AAKS....FIL......G
OfavNb    ........................................M...........WQKY....GLL......C
HvulNb    ........................................M............KEL....LTF.......
CbriNb    ........................................M............IKP....LFI.......
CbreNb    ........................................M............IKP....LVI.......
HconNb    ........................................M............KKA....LLW.......
```

```
TcanNb ........................................MGV.........CREV...WLFF.......
SratNb ........................................M...................KFL.YFSINF
ShaeNb ........................................M.............................
SmanNb ........................................MAN.........KRRIEG...........G
SjapNb ........................................M.............................
AqueNb ........................................M............SSW....VLV......F
SrosNb ........................................M............RTKQ...AAL......L
BdenNb ........................................M...........FIKT...TPLI......S
BsalNb MGLLSIW..........GHESTL.......TSIPRQLVFPM...........FMKAAVTSPGL......G
CcucNb ........................................M............HKS....VFL......F
ArepNb ........................................M............QQR....LTL......F
McirNb ........................................M............RTT....LVL......A
MbreNb ........................................M............GRL....SVL.RLSMAW
DqueNb ........................................M............HLL....PGL......A
FpinNb ........................................M............QLI....PGL......A
FradNb ........................................M............YLL....STL......T
GfroNb MITINIVGVTFLNNSAGGHWQRLECWSCSQASASCQLS.DM............QPF....PGI......A
ShirNb ........................................MH..........LQSLP..F.........S
PostNb ........................................M............KGLA...ASL......L
RsolNb ........................................M............SGI..............
NirrNb ........................................M...........CRSYH...YTL......T
ScerNb ........................................M............WKM....RHH......L
VpolNb .....................................................MKL....SSC......F
TphaNb .....................................................MKL....SNI......L
CglaNb .....................................................MKF....SRS......V
LmirNb .........................................................MKF....QWV......L
EgosNb ........................................M...................RLL......E
LngbNb ......................................................................
McolNb ......................................................................
CchtNb ......................................................................
SmajNb ......................................................................
NnodNb ......................................................................
MproNb ........................................M.............................
PsojNb MGHPNIIKVTELCTGG.GHYSEADCHD.SSMTLNPNIVIAD.............................
PhalNb MGHPNIIKVTELCTGG.GHYSEADCHDCSSMTLNPNIVIAD.............................
PinsNb MGHPNIIKVTELCTGG.GHYSEADCHDCTSISISSNLVIAD.............................
AcanNb MGHPNIIKVMELCTGG.GHYSEADCHECTEMSTSSELVIAD.............................
TclaNb MGHPNIIKVTELCTGG.GHYTEADCHDCIERSLNTNIVIAD.............................
EsilNb MRHPNIIEVTELCTGGEGRYSERDCHDC.......QLVIAN.............................
EguiNb MGHKNVVPVMELCEGG.GHYSERKCHS.PDRALDPAVVIAE.............................
DdisNb ........................................MVLN..........................
AsubNb ........................................MSLPS.........................
AcasNb ........................................MSATA.......TTNT..............
LocuNb ........................................MYQAV........RRL..............
CcarNb ........................................MYQLI........RKF..............
SansNb ........................................MYQLI........RKF..............
GproNb M.......................................R............TNL.............F
RgloNb ........................................MSFDEV...AVVYDSL....RVL.EFANEE
AbacNb ........................................M............RKVF..WPVL......F
TgonNb ME......LTDLEACS.SEDRGTESCR.SHLPSSPTS...MPPASRCKRSRKRRK.....PVD......L
HhamNb ME......LTDLEACS.SEDRGTECDR.SHLPASPTS...MPPASRCKRSRKRRK.....PVD......L
BbesNb MD......PSRRTSAG.GNEKPPESDQ.SPVAAAPS....MPPASRAKRKRKRRR....WPVD......L
CsuiNb ........TGQRREEG.GRQPGRQREP.SLSHSSRG....MPPAARAEKKRKRRRVLTT.PVD......L
CcayNb ........................CDP..............PPARSASESRRRRR....WPKD......L
TbruNb ........................................M............AQ..............C
LbraNb .....................................................MQ..............T
LseyNb ........................................M...........SPR..............D
PfalNb MGSTNISNNNISNNTSTSHNISHNCHNCLKLSEKKSLQSDMSEQDF.FINVFSQKLYIKFHLIPYFKREF
```

```
                   20                  30
                    .                   .
HsapNb1  L..LLLL.LL.........R.AVLAVPLERG.............APNKEE....................
PtroNb1  L..LLLL.LL.........R.AVLAVPLERG.............APHKEE....................
PpanNb1  L..LLLL.LL.........R.AALAVPLERG.............APNKEE....................
PabeNb1  L..LLLL.LL.........R.AVLAVPLERG.............APNKEE....................
GgorNb1  L..LLPL.LF.........R.AVLAVPLERG.............APNKEE....................
CatyNb1  L..LLLV.LL.........R.AVLAVPLERG.............APNKEE....................
MmulNb1  L..LLLV.LL.........R.AVLAVPLERG.............APNKEE....................
RroxNb1  L..LLLV.LL.........R.AVLAVPLERG.............APNKEE....................
CangNb1  L..LLLV.LL.........G.AVLAVPLERG.............APNKEE....................
MfasNb1  L..LLLV.LL.........R.AVLAVPLERG.............APNKEE....................
MnemNb1  L..LLLV.LL.........R.AVLAVPLERG.............APNKEE....................
CcapNb1  P..LLLL.LL.........R.AVLAVPLERG.............VPNKEE....................
OgarNb1  L..LLLLPLL.........R.SVLAVPLERG.............APNREE....................
PcoqNb1  L..LLLL.LL.........R.SVLAVPLERG.............VPNREE....................
GvarNb1  L..LLLL.LL.........R.AVLAVPLERA.............APNKEQ....................
TchiNb1  L..LLLL.LL.........P.AMLAVPLERG.............APNKEE....................
BmutNb1  ....SLL.LL.........R.AVLAVPLERG.............AP.KEE....................
MjavNb1  ....LLL.LL.........R.AVLAVPLERG.............AP.KQE....................
VpacNb1  ....PLL.LL.........R.VILAVPLERG.............AT.QKE....................
CbacNb1  ....PLL.LL.........R.VILAVPLERG.............AT.QKE....................
VpacNb1  ....PLL.LL.........R.VILAVPLERG.............AT.QKE....................
CferNb1  ....PLL.LL.........R.VILAVPLERG.............AT.QKE....................
HarmNb1  ....SLL.LL.........R.AVLTVPLERG.............AP.KEE....................
RsinNb1  ....PLL.LL.........R.AVLTVPLERG.............PS.KEE....................
JjacNb1  F..FLLL.PL.........S.AALAVPLERG.............SPG.QN....................
DordNb1  ...FLLL.LL.........P.AALAVPLERG.............APRKEE....................
NgalNb1  ....PPL.LI.........L.CALAVPLDRA.............VPQQEA....................
RnorNb1  ....PLL.ML.........S.AVLAVPVDRA.............APHQED....................
MmusNb1  ....PLL.ML.........S.AVLAVPVDRA.............APPQED....................
MaurNb1  ....PLL.ML.........S.VVLAVPLDRG.............APHQED....................
CasiNb1  ....LLL.LL.........R.TVLAVPLERG.............PAKKDE....................
NparNb   I..LLFP.IL.........G.LSFAVPIERA.............PPKQEE....................
XlaeNb1  L..LFFS.FL.........G.LGLTVPIERA.............TPKKEE....................
XtroNb1  L..LLFS.FL.........G.LGLAVPIERP.............TPKKEA....................
PmucNb1  F..MGFA.LL.........G.LAICVPIERP.............KEKEEA....................
PbivNb1  F..MGAA.LL.........G.LATCVPIERP.............KEKQEA....................
GjapNb1  L..VSAA.LL.........G.LATCVPIERP.............KEKQEA....................
DrerNb1  ...LLLS.LC.........L.LVWAVPIDRN.............PDPPQE....................
TrubNb1  ...LLIS.VW.........V.GVWSVPIERN.............DVQQEA....................
TnigNb1  L..LLMA.VW.........V.GVCSVPIDRN.............DVEPEA....................
IpunNb   L..VLSS.LL.........F.SAQSVPISVD.............KTKVSV....................
DrerNb2  L..VLAS.VL.........S.WAQAVPISID.............KTKVKI....................
TrubNb2  I..LLLV.QL.........L.YLEAVPISID.............KTKVKQ....................
HsapNb2  L..LITC.LL.........T.ALEAVPIDID.............KTKVQN....................
PpanNb2  L..LITC.LL.........T.ALEAVPIDID.............KTKVQN....................
GgorNb2  L..LITC.LL.........T.ALEAVPIDID.............KTKVQN....................
CatyNb2  ......................................................................
CangNb2  ......................................................................
RroxNb2  L..LITC.LL.........T.ALEAVPIDID.............KTKVQN....................
PabeNb2  L..LITC.LL.........T.ALEAVPIDID.............KTKVQN....................
MmulNb2  L..LITC.LL.........T.ALEAVPIDID.............KTKVQN....................
MfasNb2  L..LITC.LL.........T.ALEAVPIDID.............KTKVQN....................
CcapNb2  L..LITF.LL.........T.ALEAVPIDID.............KTKVQN....................
PtroNb2  L..LITC.LL.........T.ALEAVPIDID.............KTKVQN....................
OgarNb2  L..LVTF.LL.........T.ALEAVPIDID.............KTKVQN....................
PcoqNb2  L..LVTC.LL.........T.ALQAVPIDID.............KTKVQN....................
TchiNb2  L..LVTH.LF.........T.ALEAVPIDVD.............KTKVQN....................
MjavNb2  L..LVTC.LL.........P.SLEAVPIDID.............KTKVKN....................
HarmNb2  L..LVTC.LL.........I.ALEAVPIDID.............KTKVKN....................
RsinNb2  L..LVTC.LL.........V.ALEAVPIDID.............KTKVKN....................
DordNb2  F..LVSC.LL.........T.TFEAVPIDID.............KTKVQN....................
BmutNb2  L..LVTC.AL.........P.ALEAVPIDID.............KTKVKT....................
VpacNb2  L..FITC.VL.........T.ALEAVPIDID.............KTKVKN....................
CferNb2  L..FITC.VL.........T.ALEAVPIDID.............KTKVKN....................
CbacNb2  L..FITC.VL.........T.ALEAVPIDID.............KTKVKN....................
JjacNb2  L..LALC.VL.........T.ALEAVPIDVD.............KTKVQ.....................
CasiNb2  L..LVTC.LL.........I.AFEAVPIDID.............KTKVKS....................
NgalNb2  L..LVSC.VL.........S.ALEAVPIDVD.............KTKVQT....................
RnorNb2  L..LVPC.VL.........T.ALEAVPIDVD.............KTKVHN....................
MmusNb2  L..LVPC.ML.........T.ALEAVPIDVD.............KTKVHN....................
MaurNb2  L..LVPC.ML.........T.ALEAVPIDVD.............KTKVHN....................
PbivNb2  L..LLTY.VV.........I.ILEAVPIDVD.............KTKVQE....................
PmucNb2  L..LLTY.VI.........I.ILEAVPIDVD.............KTKVKE....................
GjapNb2  L..LLTY.LV.........I.TLEAVPIDVD.............KTKVQG....................
LchaNb   F..LLIC.LL.........V.VSDAVPIDID.............KTKVKD....................
XlaeNb2  L..LLIT.SL.........V.SVTAVPIDKD.............KTKVKE....................
XtroNb2  L..LLLT.SL.........A.AVSAVPIDKD.............KAKVKV....................
AcalNb   V..LGLL.CV.........P.LIMGKPVRRG.............PPAPKP....................
BglaNb   ......................................................................
LgigNb   ......................................................................
DmagNb   ...VIGV.LV.........N.NINCAPVDPK.............KSTEKP....................
DpulNb   ......................................................................
HrobNb   F..LFAA.WV.........T.VVVGPPVSEK.............PAGVVV....................
HaztNb   ...FVAL.LC.........S.VIHSLPVKEK.............QPELTE....................
TurtNb   ...IICL.VF.........N.SVLSAPVEED.............KKTDGV....................
PtepNb   L..VLLV.SL.........E.MFVAPPVDKK.............KENNAK....................
SmimNb   F..LALV.AL.........Q.LVVAPPVDQK.............KKDGED....................
GoccNb   F..CLVV.TF.........Q.NIVCPPVDPN.............NQDPGP....................
DsuzNb   .G.LALI.AI.........T.SVVALPVTQN.............KKEPKE....................
DmelNb   .G.LALI.AI.........SASIVALPVTQN.............KKDHKE....................
AaegNb   F..LIVV.LL.........E.NTFSLPVVTQAP...........KTEKKE....................
CtelNb   L..GSML.FL.........D.AISALPVREK.............PAEEEN....................
OvicNb   LG.ILSL.LT.........V.LCNALPVVPKE............PVEEEK....................
SkowNb   L..LLAA.LL.........P.LIFAAPLDPK.............KQRESG....................
SpurNb   L..LLLL.IV.........H.VAQQAPVDPK.............PAEEDR....................
NvecNb   LL.VVLV.LV.........T.LCYAPPVKRG.............KPAEKA....................
EpalNb   LT.LFLV.LL.........T.CCYAPPVQPK.............KEKAEK....................
OfavNb   C..FVIA.FA.........V.LCVAPPVRKD.............RKKADN....................
HvulNb   ..GIFLA.LI.........S.LHVAPPVGKK.............EPELPA....................
CbriNb   ...LGVF.LI.........N.GIVCPPPMRQ.............QAEQAQAHDN................
CbreNb   ...VGVL.LI.........N.GIVCPPPMRQ.............QAEQAQAQDN................
HconNb   ...LSCF.SV...........IVVAPPPQRV.............PVEQQAPP..................
```

```
TcanNb   V..IVTA.SI...........LVSCAPPKRE.............PPEQQQEKVE................
SratNb   F..LLLI.LV.........V.NVIAPPPPRKIE..........EKVEEKI....................
ShaeNb   ......................................................................
SmanNb   FA.TGGSQIM.........SPNSYMAPLMTL.............PKLQSR....................
SjapNb   ......................................................................
AqueNb   LF.LSVV.SL.........S.SSEASPFRNE.............KKEEGK....................
SrosNb   V..LALVGVLAVVGCMQGMT.GAHAAPVRTAVEDTDTDNAAAADAADAAESTAPPADDAPVDADSAQDTT
BdenNb   I..LGV..LL.........T.FFTC....SHA......................................
BsalNb   M..LLA..IL.........A.FYAMLPCTSIA......................................
CcucNb   LV.LAFCALL.........T..................................................
ArepNb   L..LSVITLLCF.......TYSVSAGPGTAH.......................................
McirNb   I..LSL..MI.........A.IAHAGPGT.........................................
MbreNb   VA.LVM..LL.........A.AAYAAPVGTP.............KEEPAP....PAPQ............
DqueNb   VF.VFARLAA.........A..................................................
FpinNb   VF.VFAHLAA.........A..................................................
FradNb   FC.LYLRLAV.........A..................................................
GfroNb   LF.IYLHLVL.........A..................................................
ShirNb   LF.LYLCSVA.........A..................................................
PostNb   LA.AVAHCAY.........G..................................................
RsolNb   ......................................................................
NirrNb   I..FAALAAL............................................................
ScerNb   I.....................................................................
VpolNb   L..F..SLIL......PSQY.IFTSAEAANA.......................................
TphaNb   L..LGSSLVF.........N.IVAADVAANK.......................................
CglaNb   F.....ALLA.........V.LVSANAEVKA.......................................
LmirNb   L.....AGLA.........S.KALTSPTKMA.......................................
EgosNb   LG.LVFGGLL.........G.SVRCMP...........................................
LngbNb   ........ML.........T..................................................
McolNb   ........ML.........T..................................................
CchtNb   ........ML.........T..................................................
SmajNb   ........ML.........T..................................................
NnodNb   ........ML.........T..................................................
MproNb   ........VL.........T..................................................
PsojNb   ........LA.........T..................................................
PhalNb   ........LA.........T..................................................
PinsNb   ........LM.........T..................................................
AcanNb   ........QM.........T..................................................
TclaNb   ........QM.........T..................................................
EsilNb   ........QL.........T..................................................
EguiNb   ........SL.........S..................................................
DdisNb   ......................................................................
AsubNb   ...................T..................................................
AcasNb   ........TA.........T..................................................
LocuNb   ........VF.........T..................................................
CcarNb   ........VF.........T..................................................
SansNb   ........VF.........T..................................................
GproNb   V..LGASAMV.........V..................................................
RgloNb   IE.LALMAPL.........G......PATSAVH.....................................
AbacNb   L..L....ML.........A.AVMASPARLR.............ALDKPE....................
TgonNb   L..G....FL.........A.TVLLRPARIA.............AILRPL....................
HhamNb   L..G....FL.........A.TVLLRPARIA.............AILRPL....................
BbesNb   L..G....FL.........A.RVLRQPARIA.............AVLRPL....................
CsuiNb   L..G....FL.........A.QLLRQPARIA.............AVLKPL....................
CcayNb   L..G....FL.........S.EVLCAPERIA.............AVLQPI....................
TbruNb   FG.......V....LGKA..............................KP....................
LbraNb   VG.......V....SR.AEA.VVAAAPLR...............SKPHRS....................
LseyNb   VA.......V....SRKADA.AIVAAPLP...............ALEHRS....................
PfalNb   FFNIILL.LLYLKETIQKFT.SNFTIPIQKKINNNNNNNVEDFQKENTENGKILYLKEKAFDNNMVQNNN
```

```
                                                                              40
HsapNb1  ..........................................T...........................P
PtroNb1  ..........................................T...........................P
PpanNb1  ..........................................T...........................P
PabeNb1  ..........................................T...........................P
GgorNb1  ..........................................T...........................P
CatyNb1  ..........................................A...........................P
MmulNb1  ..........................................A...........................P
RroxNb1  ..........................................T...........................P
CangNb1  ..........................................T...........................P
MfasNb1  ..........................................A...........................P
MnemNb1  ..........................................A...........................P
CcapNb1  ..........................................P...........................P
OgarNb1  ..........................................S...........................P
PcoqNb1  ..........................................S...........................P
GvarNb1  ..........................................S...........................P
TchiNb1  ..........................................S...........................P
BmutNb1  ..........................................N...........................P
MjavNb1  ..........................................S...........................P
VpacNb1  ..........................................N...........................P
CbacNb1  ..........................................N...........................P
VpacNb1  ..........................................N...........................P
CferNb1  ..........................................N...........................P
HarmNb1  ..........................................S...........................P
RsinNb1  ..........................................S...........................P
JjacNb1  ..........................................S...........................P
DordNb1  ..........................................N...........................P
NgalNb1  ..........................................S...........................P
RnorNb1  ..........................................N...........................Q
MmusNb1  ..........................................S...........................Q
MaurNb1  ..........................................S...........................Q
CasiNb1  ..........................................S...........................P
NparNb   ..........................................E...........................R
XlaeNb1  ..........................................E.P.........................P
XtroNb1  ..........................................EPP.........................P
PmucNb1  .......................................................................
PbivNb1  .......................................................................
GjapNb1  ..........................................E...........................P
DrerNb1  ..........................................E...........................K
TrubNb1  ..........................................KEE.........................V
TnigNb1  ......................................................................K
IpunNb   ..........................................PEEKVQ......................E
DrerNb2  ..........................................PEETVK......................E
TrubNb2  ..........................................PEKEPE......................K
HsapNb2  ..........................................IHPVES......................A
PpanNb2  ..........................................IHPVES......................A
GgorNb2  ..........................................IHPVES......................A
CatyNb2  .......................................................................
CangNb2  .......................................................................
RroxNb2  ..........................................IHPVES......................A
PabeNb2  ..........................................IHPVES......................A
MmulNb2  ..........................................IHPVES......................A
MfasNb2  ..........................................IHPVES......................A
CcapNb2  ..........................................IHPVES......................A
PtroNb2  ..........................................IHPVES......................A
OgarNb2  ..........................................THPVES......................A
PcoqNb2  ..........................................THPVES......................A
TchiNb2  ..........................................IPPVES......................A
MjavNb2  ..........................................AQPVDS......................A
HarmNb2  ..........................................TQPMDS......................A
RsinNb2  ..........................................TQPVDS......................A
DordNb2  ..........................................TQPVES......................A
BmutNb2  ..........................................TQPVDS......................A
VpacNb2  ..........................................TQPVDS......................A
CferNb2  ..........................................TQPVDS......................A
CbacNb2  ..........................................TQPVDS......................A
JjacNb2  ..........................................TQPVES......................A
CasiNb2  ..........................................PEPVES......................A
NgalNb2  ..........................................TPPVES......................A
RnorNb2  ..........................................VEPVES......................A
MmusNb2  ..........................................TEPVEN......................A
MaurNb2  ..........................................TEPVES......................A
PbivNb2  ..........................................EEKEDS......................A
PmucNb2  ..........................................EEK.DS......................A
GjapNb2  ..........................................EEQVES......................A
LchaNb   ..........................................VPPVEA......................E
XlaeNb2  ..........................................EPTEDE......................Q
XtroNb2  ..........................................EPMEEE......................Q
AcalNb   ..........................................VEEN........................S
BglaNb   .......................................................................
LgigNb   .......................................................................
DmagNb   ...........................................TDANG......................V
DpulNb   ......................................................................E
HrobNb   ...........................................DDDEA......................N
HaztNb   ..........................................EQQKEA....LKDEP.............P
TurtNb   ..............................................KD......................P
PtepNb   ............................................N.........................S
SmimNb   .............................................KKE......................T
GoccNb   ..........................................PHEV........................N
DsuzNb   ...........................................AESST......................P
DmelNb   ..........................................AAESST......................P
AaegNb   ..........................................EEQVKD......................S
CtelNb   ..............................................KE......................V
OvicNb   ..........................................PEE.........................E
SkowNb   ..........................................EEEEQE....KDDETDD.........IRE
SpurNb   ..........................................DRDEGE..GIAGEEILN.........IAE
NvecNb   .......................................................................
EpalNb   .......................................................................
OfavNb   .......................................................................
HvulNb   ..............................................NE......................T
CbriNb   ........................................STPQPGAQ....TQEGAPVEIEKN.....EY
CbreNb   ........................................AQQHAQQQ....PAAGQPQEQAQNTEQQGEY
HconNb   ......................................................................E
```

```
TcanNb  .......................................GKPEPGRQ......................P
SratNb  .........................................IEEKEF......................D
ShaeNb  ......................................................................
SmanNb  .........................................REA..........................
SjapNb  ......................................................................
AqueNb  .........................................RERPLH.......................
SrosNb  DAGTDNGAAKPVPVSKDEP..YKAYDPYDPYDYDDYDYGEYEELYNEQYDLYADEQLDL..........K
BdenNb  ......................................................................
BsalNb  ......................................................................
CcucNb  ......................................................................
ArepNb  ......................................................................
McirNb  ......................................................................
MbreNb  ....DTQPSPPDPASEDLPDIHLAADVVDYDEYDDEDDQYYQEQEQEQQEQEQEQQ.............E
DqueNb  ......................................................................
FpinNb  ......................................................................
FradNb  ......................................................................
GfroNb  ......................................................................
ShirNb  ......................................................................
PostNb  ......................................................................
RsolNb  ......................................................................
NirrNb  ......................................................................
ScerNb  ......................................................................
VpolNb  ......................................................................
TphaNb  ......................................................................
CglaNb  ......................................................................
LmirNb  ......................................................................
EgosNb  .....................................................................H
LngbNb  ......................................................................
McolNb  ......................................................................
CchtNb  ......................................................................
SmajNb  ......................................................................
NnodNb  ......................................................................
MproNb  ......................................................................
PsojNb  ......................................................................
PhalNb  ......................................................................
PinsNb  ......................................................................
AcanNb  ......................................................................
TclaNb  ......................................................................
EsilNb  ......................................................................
EguiNb  ......................................................................
DdisNb  ......................................................................
AsubNb  ......................................................................
AcasNb  ......................................................................
LocuNb  ......................................................................
CcarNb  ......................................................................
SansNb  ......................................................................
GproNb  ......................................................................
RgloNb  ......................................................................
AbacNb  ......................................................................
TgonNb  ......................................................................
HhamNb  ......................................................................
BbesNb  ......................................................................
CsuiNb  ......................................................................
CcayNb  ......................................................................
TbruNb  ......................................................................
LbraNb  ......................................................................
LseyNb  ......................................................................
PfalNb  VL..DNNMCNKKNIKKKMKKNNVIKFMVDNYVINDVSNNHHNESLQENNNYSVLQIPNNNKQNLDVQKFR
```

```
                                                         50        60
HsapNb1   ATESP..........D.............................TGLYYHRYLQEVIDVLETDG...HF
PtroNb1   ATESP..........D.............................TGLYYHRYLQEVIDVLETDG...HF
PpanNb1   ATESP..........D.............................TGLYYHRYLQEVIDVLETDG...HF
PabeNb1   ATESP..........D.............................TGLYYHRYLQEVIDVLETDG...HF
GgorNb1   ATESP..........D.............................TGLYYHRYLQEVIDVLETDG...HF
CatyNb1   ATESP..........D.............................TGLYYHRYLQEVIDVLETDG...HF
MmulNb1   ATESP..........D.............................TGLYYHRYLQEVIDVLETDG...HF
RroxNb1   ATESP..........D.............................TGLYYHRYLQEVIDVLETDG...HF
CangNb1   ATESP..........D.............................TGLYYHRYLQEVIDVLETDG...HF
MfasNb1   ATESP.................................................................
MnemNb1   ATESP..........D.............................TGLYYTGNLQDVIDVLETDGA..KF
CcapNb1   ATESP..........D.............................TGLYYHRYLQEVIDVLETDG...HF
OgarNb1   ATESP..........D.............................TGLYYHRYLQEVINVLETDG...HF
PcoqNb1   ATESP..........D.............................TGLYYHRYLQEVINVLETDG...HF
GvarNb1   ATDSP..........D.............................TGLYYHRYLQEVINVLETDG...HF
TchiNb1   ATESP..........D.............................TGLYYHRYLQEVINVLETDG...HF
BmutNb1   ATESP..........D.............................TGLYYHRYLQEVINVLETDG...HF
MjavNb1   ATESP..........D.............................TGLYYHRYLQEVINVLETDG...HF
VpacNb1   ATESPVSGPALLASQD.............................TGLYYHRYLQEVINVLETDG...HF
CbacNb1   ATESPVSGPALLASQD.............................TGLYYHRYLQEVINVLETDG...HF
VpacNb1   ATESP..........D.............................TGLYYHRYLQEVINVLETDG...HF
CferNb1   ATESP..........D.............................TGLYYHRYLQEVINVLETDG...HF
HarmNb1   ATESP..........D.............................TGLYYHRYLQEVINVLETDG...HF
RsinNb1   ATESP..........D.............................TGLYYHRYLQEVINVLETDG...HF
JjacNb1   ATESP..........D.............................TGLYYHRYLQEVITVLETDG...HF
DordNb1   ATESP..........D.............................TGLYYHRYLQEVINVLETDG...HF
NgalNb1   ATESP..........D.............................TGLYYHRYLQEVINVLETDG...HF
RnorNb1   ATETP..........D.............................TGLYYHRYLQEVINVLETDG...HF
MmusNb1   ATETP..........D.............................TGLYYHRYLQEVINVLETDG...HF
MaurNb1   ATESP..........D.............................TGLYYHRYLQEVINVLETDG...HF
CasiNb1   ATESP..........D.............................TGLYYHRYLQEVINVLETDG...HF
NparNb    PAESA..........D.............................TGLYYDRYLREVIDVLETDN...HF
XlaeNb1   PPEQT..........D.............................TGLYYDRYLREVIDVLETDG...HF
XtroNb1   PPEQP..........D.............................TGLYYDRYLREVIDVLETDG...HF
PmucNb1   APEPP..........D.............................TGLYYHRYLQEVINVLETDG...HF
PbivNb1   APEPP..........D.............................TGLYYHRYLQEVINVLETDG...HF
GjapNb1   PPDPA..........D.............................TGLFYHRYLQEVINVLETDG...HF
DrerNb1   AEENV..........D.............................TGLYYDRYLREVIEVLETDP...HF
TrubNb1   QEETE..........D.............................TGLYYDRYLREVIEVLETDP...HF
TnigNb1   EDETG..........D.............................TGLYYDRYLREVIEVLETDP...HF
IpunNb    PPPSV..........D.............................TGLHYDRYLREVIDFLEKDQ...HF
DrerNb2   PPQSV..........D.............................TGLHYDRYLREVIDFLEKDQ...HF
TrubNb2   PPASV..........D.............................TGLHYDHYLREVIDFLEKDE...HF
HsapNb2   KIEPP..........D.............................TGLYYDEYLKQVIDVLETDK...HF
PpanNb2   KIEPP..........D.............................TGLYYDEYLKQVIDVLETDK...HF
GgorNb2   KIEPP..........D.............................TGLYYDEYLKQVIDVLETDK...HF
CatyNb2   ......................................................................
CangNb2   ......................................................................
RroxNb2   KIEPP..........D.............................TGLYYDEYLKQVIDVLETDK...HF
PabeNb2   KIEPP..........D.............................TGLYYDEYLKQVIDVLETDK...HF
MmulNb2   KIEPP..........D.............................TGLYYDEYLKQVIDVLETDK...HF
MfasNb2   KIEPP..........D.............................TGLYYDEYLKQVIDVLETDK...HF
CcapNb2   KIESP..........D.............................TGLYYDEYLKQVIDVLETDK...HF
PtroNb2   KIEPP..........D.............................TGLYYDEYLKQVIDVLETDK...HF
OgarNb2   KIEPP..........D.............................TGLYYDEYLKQVIDVLETDK...HF
PcoqNb2   KIGPP..........D.............................TGLYYDEYLKQVIEVLETDQ...HF
TchiNb2   KIEPP..........D.............................TGLYYDEYLKQVIDVLETDQ...HF
MjavNb2   KIEPP..........D.............................TGLYYDEYLKQVIDVLETDS...HF
HarmNb2   KIEPP..........D.............................TGLYYDEYLKQVIDVLETDN...HF
RsinNb2   KIEPP..........D.............................TGLYYDEYLKQVIDVLETDS...HF
DordNb2   RIEPP..........D.............................TGLYYDEYLKQVIDVLETDQ...HF
BmutNb2   KIEPP..........D.............................TGLYYDEYLKQVIDVLETDS...HF
VpacNb2   KIEPP..........D.............................TGLYYDEYLKQVIDVLETDS...HF
CferNb2   KIEPP..........D.............................TGLYYDEYLKQVIDVLETDS...HF
CbacNb2   KIEPP..........D.............................TGLYYDEYLKQVIDVLETDS...HF
JjacNb2   KIEPP..........D.............................TGLYYDEYLKEVIDVLETDK...HF
CasiNb2   KIEPP..........D.............................TGLYYDEYLKQVIDVLETDK...HF
NgalNb2   KLEPP..........D.............................TGLYYDEYLKQVIDVLETDQ...HF
RnorNb2   RIEPP..........D.............................TGLYYDEYLKQVIEVLETDP...HF
MmusNb2   RIEPP..........D.............................TGLYYDEYLKQVIEVLETDP...HF
MaurNb2   KIEPP..........D.............................TGLYYDEYLKQVIDVLETDQ...HF
PbivNb2   KVDNT..........D.............................TGLYYDAYLRQVIDVLETDK...HF
PmucNb2   TVENP..........D.............................TGLYYDVYLRQVIEVLETNK...HF
GjapNb2   KIDNP..........D.............................TGLYYDAYLRQVIDVLETDK...HF
LchaNb    KIETP..........D.............................TGLYYDRYLREVIDVLETDK...HF
XlaeNb2   QPESP..........D.............................TGLFYDRYLREVVEVLETDR...HF
XtroNb2   KSQSA..........D.............................TGLFYDRYLREVVEVLETDR...HF
AcalNb    PEDVD..........N.............................TGLAYDSYLRQVVEVLESDS...EF
BglaNb    ......................................................................
LgigNb    ...............D.............................TGLEYDRYLREIVSVLEEDE...DF
DmagNb    GDAGG..........E.............................TGLEYNRYLQEVVQVLESDP...DF
DpulNb    LEMQQ..........E.............................TGLEYNRYLQEVVQLLESDP...DF
HrobNb    STEEE..........S.............................LGLEYERYMREVVKLLESDE...EF
HaztNb    VLQDW..........E.............................LSLEYGRYLKEVIQALESDP...SF
TurtNb    DAALN..........D.............................LGLEYGRYLQEVVAALEEDK...EF
PtepNb    TEEMD..........D.............................FGLEYGRYLAQVVQTLEEDK...EF
SmimNb    PEELD..........D.............................FGLEYGRYLQQVVQALEEDK...EF
GoccNb    NETDP..........A.............................FNLEYGRYLAEVVKVLESDR...DF
DsuzNb    ATADV..........E.............................TALEYERYLREVVEALEADP...EF
DmelNb    ATADV..........E.............................TALEYERYLREVVEALEADP...EF
AaegNb    NVEHI..........E.............................NAIEYNKYLQEVVKVLESDP...VF
CtelNb    DQEELV...................................DKLSYEKYLREMSVLMTDT...DF
OvicNb    LLEDE..........D.............................TGLEYDRYLRQVIKVLEKDP...DM
SkowNb    QEEDY..........D.............................TGLEYDRYLQEVINVLEKDP...DF
SpurNb    GDGIA..........D.............................LGLEYEGYLREVIQVLEADP...EM
NvecNb    .PEPN..........K.............................DDPEYARYLRQVIEILEKDE...DY
EpalNb    ..........D....E.............................NEPEYARYLRQVVEILEKDD...DY
OfavNb    .ESEG..........K.............................GDPEYARYLKQVIEILENDD...DY
HvulNb    GTVED..........K.............................QDAEYFRYLSQVVEVLEKDP...EF
CbriNb    DGRHP..........S.............................YQFAYTKYLEEVVKILETDP...KF
CbreNb    EGRHP..........G.............................YQFAYTKYLEEVVKILESDP...KF
HconNb    NQNQK..........M.............................YEFSYSKYLEEVVKILESEP...KF
```

```
TcanNb   PDQQQ..........L.............................YHFAYSKYLEQVVKVLESDP...KF
SratNb   EDDLP..........K.............................YEFHYSKYLEKIVQILENDP...RF
ShaeNb   ......................................................................
SmanNb   ...............E.............................EAAQYKKYLEEAPKAINVDP...KT
SjapNb   ......................................................................
AqueNb   ...............DTIMERMKKIRES............LSMKGQDEEYKRYWEELI...ANNP...DL
SrosNb   DEESYVGGDEDDDA.DPDVMSVRKEAAE............KRRKADEEEYQRYLDEMMEEMARDS...DL
BdenNb   ....................................................................TL
BsalNb   ....................................................................NL
CcucNb   .......................................................QAI..........EF
ArepNb   ....................................................................RF
McirNb   ....................................................................HR
MbreNb   QQEQQAQALGNAKA.DQ..APVRAEPKPKKLWPSVEEMRLLRVKEFDKEYERYEEETADQLVKDP...NM
DqueNb   ......................................................................
FpinNb   ......................................................................
FradNb   ......................................................................
GfroNb   ......................................................................
ShirNb   ......................................................................
PostNb   ......................................................................
RsolNb   ......................................................................
NirrNb   ......................................................................
ScerNb   ......................................................................
VpolNb   ...............G......................................................
TphaNb   ...............N......................................................
CglaNb   ...............K......................................................
LmirNb   ...............D......................................................
EgosNb   ...............G......................................................
LngbNb   ...............E......................................................
McolNb   ...............E......................................................
CchtNb   ...............E......................................................
SmajNb   ...............D......................................................
NnodNb   ...............D......................................................
MproNb   ...............E......................................................
PsojNb   ...............E......................................................
PhalNb   ...............E......................................................
PinsNb   ...............E......................................................
AcanNb   ...............E......................................................
TclaNb   ...............G......................................................
EsilNb   ...............E......................................................
EguiNb   ...............E......................................................
DdisNb   .............................................................END......
AsubNb   ...............EA.........................................QHVNQD......
AcasNb   ...............EA.........................................ATIDNGG.....
LocuNb   ...............DS.........................................RCFDAEN.....
CcarNb   ...............ES.........................................HCLEEDN.....
SansNb   ...............ES.........................................HCLEEDN.....
GproNb   ...............D........................I.......EAVLYSLELI............
RgloNb   .............................................TRVQNEKREEESLRALLINAGAWGL
AbacNb   ................................................AAAQKAAESWLVLTDAG...KY
TgonNb   ................................................VADVCGKQNW.VCTDSG...CR
HhamNb   ................................................VADVCGKQNW.VCTDSG...CR
BbesNb   ................................................VADVCGKQNWLLCTDSG...CR
CsuiNb   ................................................VADVCGKQNW.SFDDTG...SR
CcayNb   ................................................VADVAGKQNW.LHDNSG...FR
TbruNb   .............................................SDEERWNLLERVYDEMNAQQQEIAA
LbraNb   .............................................TTEQREAFLEQLYDELQGQRDEIDS
LseyNb   .............................................TADQRQAFLEHLYDELHIQQGEIHE
PfalNb   NTEETMNKIESNKN.D...........................KEERLQFEKEKKQILEILEKEIRKKKL
```

```
          70            80                    90                  100
          .             .                     .                   .
HsapNb1  REK....LQAANAEDIKS.............GKLSRELD..........FVSHHVRTKLDE.........
PtroNb1  REK....LQAANAEDIKS.............GKLSRELD..........FVSHHVRTKLDE.........
PpanNb1  REK....LQAANAEDIKS.............GKLSRELD..........FVSHHVRTKLDE.........
PabeNb1  REK....LQAANAEDIKS.............GKLSRELD..........FVSHHVRTKLDE.........
GgorNb1  REK....LQAANAEDIKS.............GKLSRELD..........FVSHHVRTKLDE.........
CatyNb1  REK....LQAANAEDIKS.............GKLSRELD..........FVSHHVRTKLDE.........
MmulNb1  REK....LQAANAEDIKS.............GKLSRELD..........FVSHHVRTKLDE.........
RroxNb1  REK....LQAANAEDIKS.............GKLSRELD..........FVSHHVRTKLDE.........
CangNb1  REK....LQAANAEDIKS.............GKLSRELD..........FVSHHVRTKLDE.........
MfasNb1  .................S.............GKLSRELD..........FVSHHVRTKLDE.........
MnemNb1  PERS...CKAANARRTSS.............GKLSRELD..........FVSHHVRTKLDE.........
CcapNb1  REK....LQAANAEDIKS.............GKLSRELD..........FVSHHVRTKLDE.........
OgarNb1  REK....LQAANAEDIKS.............GKLSRELD..........FVSHHVRTKLDE.........
PcoqNb1  REK....LQAANAEDIKS.............GKLSRELD..........FVSHHVRTKLDE.........
GvarNb1  REK....LQAANAEDIKS.............GKLSRELD..........FVSHHVRTKLDE.........
TchiNb1  REK....LQAANAEDIKS.............GKLSQELD..........FVSHHVRTKLDE.........
BmutNb1  REK....LQAANAEDIKS.............GKLSRELD..........FVSHHVRTKLDE.........
MjavNb1  REK....LQAANAEDIKS.............GKLSRELD..........FVSHHVRTKLDE.........
VpacNb1  REK....LQAANAEDIKS.............GKLSRELD..........FVSHHVRTKLDE.........
CbacNb1  REK....LQAANAEDIKS.............GKLSRELD..........FVSHHVRTKLDE.........
VpacNb1  REK....LQAANAEDIKS.............GKLSRELD..........FVSHHVRTKLDE.........
CferNb1  REK....LQAANAEDIKS.............GKLSRELD..........FVSHHVRTKLDE.........
HarmNb1  REK....LQAANAEDIKS.............GKLSRELD..........FVSHHVRTKLDE.........
RsinNb1  REK....LQAANAEDIKS.............GKLSRELD..........FVSHHVRTKLDE.........
JjacNb1  REK....LQSANAEDIKS.............GKLSRELD..........FVSHHVRTKLDE.........
DordNb1  REK....LQAANAEDIKS.............GKLSRELD..........FVSHHVRTKLDE.........
NgalNb1  REK....LQAANAEDIKS.............GKLSRELD..........FVSHHVRTKLDE.........
RnorNb1  REK....LQAANAEDIKS.............GKLSQELD..........FVSHNVRTKLDE.........
MmusNb1  REK....LQAANAEDIKS.............GKLSQELD..........FVSHNVRTKLDE.........
MaurNb1  REK....LQAANAEDIKS.............GKLSRELD..........FVSHHVRTKLDE.........
CasiNb1  REK....LQAANAEDIKS.............GKLSRELD..........FVSHHVRTKLDE.........
NparNb   REK....LQTANADDIKS.............GKLSKELD..........FVSHHVRTRLDE.........
XlaeNb1  REK....LQAANADDIKS.............GKLSKELD..........FVSHNVRTKLDE.........
XtroNb1  REK....LQAANADDIKS.............GKLSKELD..........FVSHHVRTKLDE.........
PmucNb1  REK....LQAANAEDIKS.............GKLSKELD..........FVGHSVRTKLDE.........
PbivNb1  REK....LQAANAEDIKS.............GKLSKELD..........FVSHHVRTKLDE.........
GjapNb1  REK....LQAANAEDIKS.............GKLSKELD..........FVSHHVRTKLDE.........
DrerNb1  REK....LQTANTEDIKN.............GRLSKELD..........LVGHHVRTRLDE.........
TrubNb1  REK....LQTANTEDIKN.............GHLAKELD..........LVSHHVRTRLDE.........
TnigNb1  REK....LQTANTEDIKN.............GHLAKELD..........LVSHHVRTRLDE.........
IpunNb   REK....LHNTDMEDIKQ.............GKLAKELD..........FVSHHVRTKLDE.........
DrerNb2  REK....LHNTDMEDIKQ.............GKLAEELD..........FVSHHVRSKLDE.........
TrubNb2  REK....LRNTDMEDIKQ.............GKLAKELD..........FVGHHIRTKLDE.........
HsapNb2  REK....LQKADIEEIKS.............GRLSKELD..........LVSHHVRTKLDE.........
PpanNb2  REK....LQKADIEEIKS.............GRLSKELD..........LVSHHVRTKLDE.........
GgorNb2  REK....LQKADIEEIKS.............GRLSKELD..........LVSHHVRTKLDE.........
CatyNb2  ......................................................................
CangNb2  ......................................................................
RroxNb2  REK....LQKADIEEIKS.............GRLSKELD..........LVSHHVRTKLDE.........
PabeNb2  REK....LQKADIEEIKS.............GRLSKELD..........LVSHHVRTKLDE.........
MmulNb2  REK....LQKADIEEIKS.............GRLSKELD..........LVSHHVRTKLDE.........
MfasNb2  REK....LQKADIEEIKS.............GRLSKELD..........LVSHHVRTKLDE.........
CcapNb2  REK....LQKADIEEIKS.............GRLSKELD..........LVSHHVRTKLDE.........
PtroNb2  REK....LQKADIEEIKS.............GRLSKELD..........LVSHHVRTKLDE.........
OgarNb2  REK....LQKADIEEIKS.............GRLSKELD..........LVSHHVRTKLDE.........
PcoqNb2  REK....LQKADVEEIKS.............GRLSKELD..........LVSHHVRTKLDE.........
TchiNb2  REK....LQKADIEEIKS.............GKLSKELD..........LVSHNVRTRLDE.........
MjavNb2  REK....LQKADIEEIKS.............GRLSKELD..........LVSHHVRTKLDE.........
HarmNb2  REK....LQKADIEEIKS.............GRLSKELD..........LVSHHVRTKLDE.........
RsinNb2  REK....LQKADIEEIKS.............GRLSKELD..........LVSHHVRTKLDE.........
DordNb2  REK....LQKADIEEIKS.............GRLSKELD..........LVSHHVRTKLDE.........
BmutNb2  REK....LQKADIEEIKS.............GRLSKELD..........LVSHHVRTKLDE.........
VpacNb2  REK....LQKADIEDIKS.............GRLSKELD..........LVSHHVRTKLDE.........
CferNb2  REK....LQKADIEDIKS.............GRLSKELD..........LVSHHVRTKLDE.........
CbacNb2  REK....LQKADIEDIKS.............GRLSKELD..........LVSHHVRTKLDE.........
JjacNb2  REK....LQKADIEEIKS.............GRLSKELD..........LVSHHVRTKLDE.........
CasiNb2  REK....LQKADIEEIKS.............GRLSKELD..........LVSHHVRTKLDE.........
NgalNb2  RER....LQKADIEEIKS.............GRLSKELD..........LVSHHVRTKLDE.........
RnorNb2  REK....LQKADIEEIRS.............GRLSQELD..........LVSHKVRTRLDE.........
MmusNb2  REK....LQKADIEEIRS.............GRLSQELD..........LVSHKVRTRLDE.........
MaurNb2  REK....LQKADIEEIRS.............GRLSKELD..........LVSHHVRTKLDE.........
PbivNb2  REK....LQSADIEEIKS.............GKLSKELD..........LVSHHVRTKLDE.........
PmucNb2  REK....LQTADIEEIKS.............GKLSKELD..........LVGHKVRTKLDE.........
GjapNb2  REK....LQTADIEEIKS.............GKLSKELD..........LVSHHVRTKLDE.........
LchaNb   REK....LQTANLEEIKS.............GKLSKELD..........LVSHHVRTKLDE.........
XlaeNb2  REK....LQTADIEDIKS.............GKISKELD..........LVGHHIRTKLDE.........
XtroNb2  REK....LQTADIEDIKS.............GKISKELD..........LVSHHIRTKLDE.........
AcalNb   RKK....LETANVTDIKS.............GKIAMYLE..........MVNHTIRTSLDE.........
BglaNb   .................................................MVNHTVRTSLDE.........
LgigNb   RKK....LEAANVSEIKD.............GSIARHLE..........FVGHNVRTKLDE.........
DmagNb   RQK....LEKSDPEDIRT.............GKVARELE..........YVNHHVRNKLDE.........
DpulNb   RQK....LEKSDPEDIRT.............GKVAKELE..........YVNHHVRNKLDE.........
HrobNb   RKK....LESANSSDIKS.............GAIAQHLN..........FVSKSIRSKLDE.........
HaztNb   RKK....LEDAEADDIKS.............GKIARELH..........FVDHNVRTKLDE.........
TurtNb   AKK....LENVSAEHIQS.............GAIANELN..........LVSHNIRTKLDE.........
PtepNb   AKR....LENITTDQIRS.............GAIAKELE..........FVKHNVRSKLDE.........
SmimNb   AKK....LENVSAEQIKS.............GHIAKELE..........FVKHNVRSKLDE.........
GoccNb   AKK....LENASRDEITS.............GHVAKHLD..........LVDHKIRTKLDE.........
DsuzNb   RKK....LDKAPEADIRS.............GKIAQELD..........YVNHHVRTKLDE.........
DmelNb   RKK....LDKAPEADIRS.............GKIAQELD..........YVNHHVRTKLDE.........
AaegNb   RDK....LDKAAESDIRS.............GKIAQELE..........YVNHNVRSQLDE.........
CtelNb   AQK....MMKIQDIAGQS.............GEIAKHLE..........FVSHGVRSKLDE.........
OvicNb   RKR....MEEMSLEDLKE.............GNFARELG..........FLSSNIRSKLDE.........
SkowNb   KKK....LEEADIEHIKS.............GRIADELN..........LVDHQVRSKLDE.........
SpurNb   KSH....MDEMDTNDLLT.............GKFGKQLN..........NVGSHIRGQLDE.........
NvecNb   VRK....LMNASDDDLRS.............GRIAEDID..........LVKHDVRSKLDE.........
EpalNb   VKK....LMNASDEDLRS.............GKVADDLD..........FVKHDVRTKLDE.........
OfavNb   VKR....LLNASDEELRT.............GRVADDID..........LVKHDVRTKLDE.........
HvulNb   KSK....LHNASEEDIRS.............GKIANYLD..........LVGHGVRLKLDE.........
CbriNb   NER....LKNMNEEDIKA.............GKIADHID..........DLPQEVFDKLHR.........
CbreNb   NDR....LKNMNEEEIKA.............GKIADHID..........DLPAEVFDKLHK.........
HconNb   SER....LRGMAEEDIKA.............GKIADHID..........DLDSHVFDKLTK.........
```

```
TcanNb   TER....LKNMPEGDIKS.............GKIADHIE..........ELSSHTFEQLTK.........
SratNb   NEK....LKTLSEEDIRS.............GKIADHFD..........VISKDVAEELTK.........
ShaeNb   ......................................................................
SmanNb   LDS....ARKVFVENQLP.............SFMKFQLDRQIPFNDLLLWNTSLRDKDDE.........
SjapNb   ......................................................................
AqueNb   AKR....FREQKAEDLAA.............G.LGDRLS..........HRDGGIRSQMEE.........
SrosNb   RED....L....IEDIQGVQEAERAPGDKDILKIALRLKN...........EERVRNIIEE.........
BdenNb   SSN...................................................................
BsalNb   SEK...................................................................
CcucNb   KNK...................................................................
ArepNb   NEE...................................................................
McirNb   YKE...................................................................
MbreNb   AKSVLEALRANKDDSVAEKAVE.........AQVAQKLQ.............RGFRSRLDEQPRLAGQAQ
DqueNb   ......................................................................
FpinNb   ......................................................................
FradNb   ......................................................................
GfroNb   ......................................................................
ShirNb   ......................................................................
PostNb   ......................................................................
RsolNb   ......................................................................
NirrNb   ......................................................................
ScerNb   ......................................................................
VpolNb   ...............................STVNVEVE...............................
TphaNb   ...............................PEVNVEVE...............................
CglaNb   ...............................DDVNVEVE...............................
LmirNb   ...............................PYLNVEVD...............................
EgosNb   ...............................DKIHTEVE...............................
LngbNb   ......................................................................
McolNb   ......................................................................
CchtNb   ......................................................................
SmajNb   ......................................................................
NnodNb   ......................................................................
MproNb   ......................................................................
PsojNb   ......................................................................
PhalNb   ......................................................................
PinsNb   ......................................................................
AcanNb   ......................................................................
TclaNb   ......................................................................
EsilNb   ......................................................................
EguiNb   ......................................................................
DdisNb   ...................................................KEFVKKLFDS.........
AsubNb   ...................................................TQYIRTLFQS.........
AcasNb   ..................................................VMSKVREVFAK.........
LocuNb   ...................................................KKKYEELFKK.........
CcarNb   ...................................................TKTFADLFEK.........
SansNb   ...................................................TKTFAELFEK.........
GproNb   ......................................................................
RgloNb   VSR....KRRGSSQNLPETTSMPANP....AATKSIKLQDF.......RKVFHRVSPWKSS.........
AbacNb   AESW.....NTASRMFKA.............GVTEEQWQ..........SAMHRYRDPLGA.........
TgonNb   RSAW.....DVQSRLGYV.............GDLREQSV..........SALRNFREPPGA.....GSTD
HhamNb   RSAW.....DVQSRLGYV.............GDLREQSV..........SALRNFREPLGA.....GSTD
BbesNb   RSAW.....DIQSRLGYV.............GDLREQSI..........SALKNFREPLGQ.....GMAD
CsuiNb   LSAW.....DVQSRLGYV.............GDLREQSV..........SALKSF...MGG.....GGA.
CcayNb   RSAW.....EAQSRLGYV.............GDLRELPL..........SAVKSY..P............
TbruNb   RKRELDEFYDGNIEKIEEALHFNENR....RGSREGDWDTY.......EKVEHIVRAKSEVQE.......
LbraNb   RKRQMDEVYESNHHDIHDLLFANLDA....AHGALGDWDTW.......EKVEAAVKPQKNEMC.......
LseyNb   RKNQMDDMFESNQQQIHDILFAKVDT....AGGALGDWDTW.......EKVEMLMRPQREEAA.......
PfalNb   KEK..EFLKNESIDDIKSDSLSFISP...TFNNIEKEQEDRNEINDLHKSKSKNVRNKIDD.........
```

```
                                        110       120
                                         .         .
HsapNb1   ........................LKRQEVSRLRMLLKAKMDAEQD.PNVQ...................
PtroNb1   ........................LKRQEVSRLRMLLKAKMDAEQD.PNVQ...................
PpanNb1   ........................LKRQEVSRLRMLLKAKMDAEQD.PNVQ...................
PabeNb1   ........................LKRQEVSRLRMLLKAKMDAEQD.PNVQ...................
GgorNb1   ........................LKRQEVSRLRMLLKAKMDAEQD.PNVQ...................
CatyNb1   ........................LKRQEVSRLRMLLKAKMDAEQD.PNVQ...................
MmulNb1   ........................LKRQEVSRLRMLLKAKMDAEQD.PNVQ...................
RroxNb1   ........................LKRQEVSRLRMLLKAKMDAEQD.PNVQ...................
CangNb1   ........................LKRQEVSRLRMLLKAKMDAEQD.PNVQ...................
MfasNb1   ........................LKRQEVSRLRMLLKAKMDAEQD.PNVQ...................
MnemNb1   ........................LKRQEVSRLRMLLKAKMDAEQD.PNVQ...................
CcapNb1   ........................LKRQEVSRLRMLLKAKMDAQQE.PNVQ...................
OgarNb1   ........................LKRQEVSRLRMLLKAKIDAEQE.PNVQ...................
PcoqNb1   ........................LKRQEVSRLRMLLKAKMDAEQE.PNVQ...................
GvarNb1   ........................LKRQEVSRLRMLLKAKMDAEQE.PNVQ...................
TchiNb1   ........................LKRQEVSRLRMLLKAKMDAAQE.PNAQ...................
BmutNb1   ........................LKRQEVSRLRMLLKAKMDAQQE.PNIQ...................
MjavNb1   ........................LKRQEVSRLRMLLKAKMDAEQE.PNIQ...................
VpacNb1   ........................LKRQEVSRLRMLLKAKMDAEQE.PNVQ...................
CbacNb1   ........................LKRQEVSRLRMLLKAKMDAEQE.PNVQ...................
VpacNb1   ........................LKRQEVSRLRMLLKAKMDAEQE.PNVQ...................
CferNb1   ........................LKRQEVSRLRMLLKAKMDAEQE.PNVQ...................
HarmNb1   ........................LKRQEVSRLRMLLKAKMDAEQE.PNVQ...................
RsinNb1   ........................LKRQEVSRLRMLLKAKMDAEQE.PSVQ...................
JjacNb1   ........................LKRQEVSRLRMLLKAKMDAEQE.PNVQ...................
DordNb1   ........................LKRQEVSRLRMLLKAKMDAEQE.PNVQ...................
NgalNb1   ........................LKRQEVSRLRMLLKAKMDAEQE.PNMQ...................
RnorNb1   ........................LKRQEVSRLRMLLKAKMDAKQE.PNLQ...................
MmusNb1   ........................LKRQEVSRLRMLLKAKMDAKQE.PNLQ...................
MaurNb1   ........................LKRQEVSRLRMLLKAKMDAKQE.PNLQ...................
CasiNb1   ........................LKRQEVSRLRMLLKAKLDAEQG.PNVQ...................
NparNb    ........................LKRQEVSRLRMLIKAKMDATME.ENVQ...................
XlaeNb1   ........................LKRQEVSRLRMLIKAKMDATME.ENVQ...................
XtroNb1   ........................LKRQEVSRLRMLIKAKMDATME.ENVQ...................
PmucNb1   ........................LKRQEVSRLRMLLKAKMDATME.QDVQ...................
PbivNb1   ........................LKRQEVSRLRMLLKAKMDAAME.QDVQ...................
GjapNb1   ........................LKRQEVSRLRMLLKAKMDATME.QDVQ...................
DrerNb1   ........................LKRQEVSRLRMLLKAKLDSTNA.QSVQ...................
TrubNb1   ........................LKRQEVSRLRMLFKAKLDSTNI.QTQQ...................
TnigNb1   ........................LKRQEVSRLRMLFKAKLDSTNV.QTQQ...................
IpunNb    ........................LKRQEVSRLRTLIRAKQDIEGG.NGVV...................
DrerNb2   ........................LKRQEVSRLRTLIKAKQDIEGG.NDIA...................
TrubNb2   ........................LKRQEVNRLRTLIKAKQDLEGG.NDIS...................
HsapNb2   ........................LKRQEVGRLRMLIKAKLD..SL.QDIG...................
PpanNb2   ........................LKRQEVGRLRMLIKAKLD..SL.QDIG...................
GgorNb2   ........................LKRQEVGRLRMLIKAKLD..SL.QDIG...................
CatyNb2   ......................................................................
CangNb2   ......................................................................
RroxNb2   ........................LKRQEVGRLRMLIKAKLD..SL.QDIG...................
PabeNb2   ........................LKRQEVGRLRMLIKAKLD..SL.QDIG...................
MmulNb2   ........................LKRQEVGRLRMLIKAKLD..SL.QDIG...................
MfasNb2   ........................LKRQEVGRLRMLIKAKLD..SL.QDIG...................
CcapNb2   ........................LKRQEVGRLRMLIKAKLD..SI.QDIG...................
PtroNb2   ........................LKRQEVGRLRMLIKAKLD..SL.QDIG...................
OgarNb2   ........................LKRQEVGRLRMLIKAKLD..SF.QGVG...................
PcoqNb2   ........................LKRQEVGRLRMLIKAKLD..SF.QDKG...................
TchiNb2   ........................LKRQEVGRLRMLIKAKSD..SH.QDVG...................
MjavNb2   ........................LKRQEVARLRMLIKAKLD..SL.QDTG...................
HarmNb2   ........................LKRQEVGRLRMLIKAKLD..SL.QDTG...................
RsinNb2   ........................LKRQEVGRLRMLIKAKLD..SL.HDTG...................
DordNb2   ........................LKRQEVGRLRMLIKAKLD..SL.QDVG...................
BmutNb2   ........................LKRQEVARLRMLIKAKLD..SF.QDTG...................
VpacNb2   ........................LKRQEVARLRMLIKAKLD..SL.QDTG...................
CferNb2   ........................LKRQEVARLRMLIKAKLD..SL.QDTG...................
CbacNb2   ........................LKRQEVARLRMLIKAKLD..SL.QDTG...................
JjacNb2   ........................LKRQEVGRLRMLIKAKLD..SL.QDVG...................
CasiNb2   ........................LKRQEVARLRMLIKAKLD..SH.QDVG...................
NgalNb2   ........................LKRQEVGRLRMLIKAKLD..SL.QDVG...................
RnorNb2   ........................LKRQEVGRLRMLIKAKLD..AL.QDTG...................
MmusNb2   ........................LKRQEVGRLRMLIKAKLD..AL.QDTG...................
MaurNb2   ........................LKRQEVARLRMLIKAKLD..SL.QDTG...................
PbivNb2   ........................LKRQEVARLRMLIKAKID..AY.QDSG...................
PmucNb2   ........................LKRQEVARLRMLIKAKID..AY.QDSG...................
GjapNb2   ........................LKRQEVARLRMLIKAKMD..SF.QDSG...................
LchaNb    ........................LKRQEVARLRMLIRAKMD..AQ.QELG...................
XlaeNb2   ........................LKRQEVSRLRMLIRAKMD..LG.DNTG...................
XtroNb2   ........................LKRQEVARLRMLIRAKMD..QG.DDTG...................
AcalNb    ........................IKRREVARLQDLARLEMQ..AV.SGTE...................
BglaNb    ........................IKRREISRLQELARLQMQ..GM.SAHD...................
LgigNb    ........................IKRREVDRLQELARLKMK..QM.KGTE...................
DmagNb    ........................LKRQEMERLRHLAMEEYE..RA.RGLG...................
DpulNb    ........................LKRQEMERLRHLAMEEYE..RA.RGLG...................
HrobNb    ........................LKRNELERLRTVLRLKVR..QA.HPDS...................
HaztNb    ........................LKRRELERIRHLAVQANE..LS.NGLD...................
TurtNb    ........................LKRIEIERLRKLTKLEHD..IKEFGSFSN.........DGRRWRTL
PtepNb    ........................LKRIEVDRLRKLIQEQME..RTELGLYRDGDGHLLNTPDGRRWRTI
SmimNb    ........................LKRMEVDRLRKLIREQME..RTELGLN...................
GoccNb    ........................LKRIEIDRLRKLTQKKNE..LE.HGID...................
DsuzNb    ........................IKRREVERLRELANQAYE..LS.NDID...................
DmelNb    ........................IKRREVERLRELANQAYE..LS.NDID...................
AaegNb    ........................LKRIELQRLRELATRQFE..LS.NNID...................
CtelNb    ........................IKRKEVQRLKQITKLQEK..LK.NGMN...................
OvicNb    ........................LKRMEVQRLRTVAKQRME..AA.TGTGL..................
SkowNb    ........................LKRHEINRLRLVAREQLD..QK.NGVK...................
SpurNb    ........................LKKKEIQRLRIIARKAME..QQ.AHDD...................
NvecNb    ........................LKRQEVERQRMIRRQMND..HL.NGIK...................
EpalNb    ........................LKRQEIERQRQIRRQMND..HQ.AGLT...................
OfavNb    ........................LKRQEVERQRMIRRQMND..HL.NGLK...................
HvulNb    ........................IKRTEVEYQRELLRQRQD..FM.SGIE...................
CbriNb    ........................AKFEEIERLRKQIEEQIK..AD.GG.....................
CbreNb    ........................AKFEEIERLRKQIEEQIK..AD.GG.....................
HconNb    ........................AKLDEIERLRAQIQKQIE..AD.GG.....................
```

```
TcanNb  ........................LKLAEIERLREVIAKQIE..AD.GG.....................
SratNb  ........................AKLQEVERLRQAILKEVE..EG.KK.....................
ShaeNb  ......................................................................
SmanNb  ........................TKRYLVEKLIKTHLEKAA............................
SjapNb  ......................................................................
AqueNb  ........................LQRRNIEELRDLHRQKMA..GE.SHEP...................
SrosNb  ........................KERQLLEAQRHEARRRAE..AR.MHVP...................
BdenNb  ......................................................................
BsalNb  ......................................................................
CcucNb  ......................................................................
ArepNb  ......................................................................
McirNb  ......................................................................
MbreNb  AIFLGMLNTCLSSTGTGLIRCAISQKRQVIERMRRAAKRADN..TE........................
DqueNb  ...............................................HGDH...................
FpinNb  ...............................................HGEH...................
FradNb  ...............................................HG.....................
GfroNb  ...............................................HG.....................
ShirNb  ...............................................HGAH...................
PostNb  ...............................................HGGH...................
RsolNb  ......................................................................
NirrNb  ......................................................................
ScerNb  ......................................................................
VpolNb  ......................................................................
TphaNb  ......................................................................
CglaNb  ......................................................................
LmirNb  ......................................................................
EgosNb  ......................................................................
LngbNb  ........................LQNRKLLKLFCM..................................
McolNb  ........................LQKRKLIKFFSM..................................
CchtNb  ........................LQTRKLTKFFSM..................................
SmajNb  ........................FQKRKLMKLFSM..................................
NnodNb  ........................LQQRKLTKLFSM..................................
MproNb  ........................LQKRKLKKLFDL..................................
PsojNb  ........................SEIAELKNQFMA..................................
PhalNb  ........................GEIAELKNQFMA..................................
PinsNb  ........................SEISELKNQFMA..................................
AcanNb  ........................SDIQELKRQFMA..................................
TclaNb  ........................AEITELKKQFEM..................................
EsilNb  ........................ADIGHLRDAFRE..................................
EguiNb  ........................EEIAGLREMFQA..................................
DdisNb  ......................................................................
AsubNb  ......................................................................
AcasNb  ......................................................................
LocuNb  ......................................................................
CcarNb  ......................................................................
SansNb  ......................................................................
GproNb  ........................LKKGINERL.....................................
RgloNb  ...........NSSG.........LASTQIQAPSTPSLKSNK.....SGSS...................
AbacNb  ........................VKSRKLSSA.....................................
TgonNb  RV......TAFRTRKKALKTF...PDDKKVDGE.....................................
HhamNb  SA......AVFRTRKKALKTF...PENRKVDGE.....................................
BbesNb  GR......TAHSRRSDPLDA....PSIRRASA......................................
CsuiNb  ........................PSTRDTTGE.....................................
CcayNb  ........................PRLKSAAAT.....................................
TbruNb  ........................VLEERIRLLGDIEASQKQRETTLSSVQSE................L
LbraNb  ........................AMQSHIEKLREVLQAKSATTGNLDSVQKE................L
LseyNb  ........................AMRSRIEALNEILNERNATTQTCKATEQE................L
PfalNb  .FLINNMNTIIKKYKKGLILKG..LKSFDIKNLKSLQKNKIETLNKHEGIKNK.........DGVKEKVK
```

```
                130                            140                  150
                  .                             .                    .
HsapNb1   .....VDH....................LNLLKQFEHL.........DPQNQHTFE.....ARDLELLIQ
PtroNb1   .....VDH....................LNLLKQFEHL.........DPQNQHTFE.....ARDLELLIQ
PpanNb1   .....VDH....................LNLLKQFEHL.........DPQNQHTFE.....ARDLELLIQ
PabeNb1   .....VDH....................LNLLKQFEHL.........DPQNQHTFE.....ARDLELLIQ
GgorNb1   .....VDH....................LNLLKQFEHL.........DPQNQHTFE.....ARDLELLIQ
CatyNb1   .....VDH....................LNLLKQFEHL.........DPQNQHTFE.....ARDLELLIQ
MmulNb1   .....VDH....................LNLLKQFEHL.........DPQNQHTFE.....ARDLELLIQ
RroxNb1   .....VDH....................LNLLKQFEHL.........DPQNQHTFE.....ARDLELLIQ
CangNb1   .....VDH....................LNLLKQFEHL.........DPQNQHTFE.....ARDLELLIQ
MfasNb1   .....VDH....................LNLLKQFEHL.........DPQNQHTFE.....ARDLELLIQ
MnemNb1   .....VDH....................LNLLKQFEHL.........DPQNQHTFE.....ARDLELLIQ
CcapNb1   .....VDH....................LNLLKQFEHL.........DPQNQHTFE.....ARDLELLIQ
OgarNb1   .....VDH....................LNLLKQFEHL.........DPQNQHTFE.....ARDLELLIQ
PcoqNb1   .....VDH....................LNLLKQFEHL.........DPQNQHTFE.....ARDLELLIQ
GvarNb1   .....VDH....................LNLLKQFEHL.........DSQNQHTFE.....ARDLELLIQ
TchiNb1   .....MDH....................LNLLKHFEHL.........DPQNQHTFE.....ARDLELLIQ
BmutNb1   .....LDH....................LNLLKQFEHL.........DPQNQHTFE.....ARDLELLIQ
MjavNb1   .....LDH....................LSLLKQFEHL.........DPQNQHTFE.....ARDLELLIQ
VpacNb1   .....LDH....................LSLLKQFEHL.........DPQNQHTFE.....ARDLELLIQ
CbacNb1   .....LDH....................LSLLKQFEHL.........DPQNQHTFE.....ARDLELLIQ
VpacNb1   .....LDH....................LSLLKQFEHL.........DPQNQHTFE.....ARDLELLIQ
CferNb1   .....LDH....................LSLLKQFEHL.........DPQNQHTFE.....ARDLELLIQ
HarmNb1   .....LDH....................LSLLKQFEHL.........DPQNQHTFE.....ARDLELLIQ
RsinNb1   .....LDH....................LSLLKQFEHL.........DPQNQHTFE.....ARDLELLIQ
JjacNb1   .....VDH....................LNLLKQFEHL.........DPQNQHTFE.....ARDLELLIQ
DordNb1   .....LDH....................LNLLKQFEHL.........DPQNQHTFE.....ARDLELLIQ
NgalNb1   .....MDH....................LNLLKQFEHL.........DPQNQHTFE.....ARDLELLIQ
RnorNb1   .....VDH....................MNLLKQFEHL.........DPQNQHTFE.....ARDLELLIQ
MmusNb1   .....VDH....................MNLLKQFEHL.........DPQNQHTFE.....ARDLELLIQ
MaurNb1   .....VDH....................LNLLKQFEHL.........DPQNQHTFE.....ARDLELLIQ
CasiNb1   .....VDH....................MSLLKQFEHL.........DPQNQHTFE.....ARDLELLIQ
NparNb    .....IDH....................LALLKQFEHL.........DPQNQHTFE.....ARDLELLIQ
XlaeNb1   .....IDH....................MSLLKQFEHL.........DPQNQHTFE.....ARDLELLIQ
XtroNb1   .....IDH....................MSLLKQFEHL.........DPQNQHTFE.....ARDLELLIQ
PmucNb1   .....VDH....................LALLKQFEHL.........DSQNQHTFE.....ARDLELLIK
PbivNb1   .....VDH....................LSLLRQFEHL.........DSQNQHTFE.....AQDLELLIK
GjapNb1   .....IDH....................RTLLKQFEHL.........DPQNQHTFE.....ARDLELLIQ
DrerNb1   .....MDH....................ASLLKQFEHL.........DPHNQNTFE.....AKDLELLIA
TrubNb1   .....MDH....................ASLLKQFEHL.........DPHNQNTFE.....AKDLELLIS
TnigNb1   .....MDH....................ASLLKQFEHL.........DPHNQNTFE.....AKDLELLIS
IpunNb    .....VDH....................QALLKQFEYL.........NHMNPHTFE.....VDDLDRLIK
DrerNb2   .....VDH....................QALLKQFEYL.........NHMNPHTFE.....VEDLDRLIK
TrubNb2   .....VDH....................QALLKQFEYL.........NHMNPHTFE.....VDDLDLLIK
HsapNb2   .....MDH....................QALLKQFDHL.........NHLNPDKFE.....STDLDMLIK
PpanNb2   .....MDH....................QALLKQFDHL.........NHLNPDKFE.....STDLDMLIK
GgorNb2   .....MDH....................QALLKQFDHL.........NHLNPDKFE.....STDLDMLIK
CatyNb2   .....MDH....................QALLKQFDHL.........NHLNPDKFE.....STDLDMLIK
CangNb2   ......................................................................
RroxNb2   .....MDH....................QALLKQFDHL.........NHLNPDKFE.....STDLDMLIK
PabeNb2   .....MDH....................QALLKQFDHL.........NHLNPDKFE.....STDLDMLIK
MmulNb2   .....MDH....................QALLKQFDHL.........NHLNPDKFE.....STDLDMLIK
MfasNb2   .....MDH....................QALLKQFDHL.........NHLNPDKFE.....STDLDMLIK
CcapNb2   .....MDH....................QALLKQFDHL.........NHLNPDKFE.....STDLDMLIK
PtroNb2   .....MDH....................QALLKQFDHL.........NHLNPDKFE.....STDLDMLIK
OgarNb2   .....MDH....................QALLKQFDHL.........NHMNPDKFE.....STDLDMLIK
PcoqNb2   .....MDH....................QSLLKQFDHL.........NHLNPDKFE.....STDLDMLIK
TchiNb2   .....IDH....................QALLKQFDHL.........NHLNPDTFE.....STDLDMLIK
MjavNb2   .....IDH....................HALLKQFDHL.........NHLNPDKFE.....STDLDMLIK
HarmNb2   .....IDH....................HALLKQFDHL.........NHLNPDKFE.....STDLDMLIK
RsinNb2   .....IDH....................HALLKQFDHL.........NHLNPDKFE.....STDLDMLIK
DordNb2   .....IDH....................QALLKQFDHL.........NHMNPDKFE.....STDLDMLIK
BmutNb2   .....MDH....................HALLKQFDHL.........NHLNPDKFE.....STDLDMLIK
VpacNb2   .....IDH....................HALLKQFDHL.........NHLNPDKFE.....STDLDMLIK
CferNb2   .....IDH....................HALLKQFDHL.........NHLNPDKFE.....STDLDMLIK
CbacNb2   .....IDH....................HALLKQFDHL.........NHLNPDKFE.....STDLDMLIK
JjacNb2   .....IDH....................QALLKQFDHL.........NHQNPDKFE.....STDLDMLIK
CasiNb2   .....INH....................QALLKQFDHL.........NHMNPDTFE.....STDLDMLIK
NgalNb2   .....LNH....................QALLKQFDHL.........NHQNPDTFE.....STDLDMLIK
RnorNb2   .....MNH....................HLLLKQFEHL.........NHQNPDTFE.....SKDLDMLIK
MmusNb2   .....MNH....................HLLLKQFEHL.........NHQNPNTFE.....SRDLDMLIK
MaurNb2   .....MNH....................QLLLKQFEHL.........NHQNPDTFE.....SRDLDMLIK
PbivNb2   .....IDH....................QALLKQFEHL.........NHHNPHTFE.....VKDLDMLIK
PmucNb2   .....VDH....................QALLKQFGHL.........NHHNPHTFE.....VKDLDMLIK
GjapNb2   .....IDH....................QSLLKQFEHL.........NHQNPHTFE.....AKDLDMLIR
LchaNb    .....VDH....................QALLKQFQHL.........NHHNPDTFE.....PKDLDMLIK
XlaeNb2   .....MDH....................RALLKQFEHL.........NHNNPHSFE.....PQDLDLLIK
XtroNb2   .....MDH....................RALLRQFEHL.........NQNNPHSFE.....PQDLDLLIR
AcalNb    .....LIMNAL.............LGGGIKNMEIPSHL.........DVRNPHSFE.....MKDLEALIK
BglaNb    ...........................GVKKFEIPSYL.........DVRNPHSFE.....VKDLENLII
LgigNb    .....ILHN................IAYHGDIRIPGHL.........DVMNPHSFE.....MKDLESLIK
DmagNb    .....IPH....................DGRLKIPGHL.........DHKSP.SFE.....SEDLRKLIV
DpulNb    .....IPH....................DGRLKIPGHL.........DHKSP.SFE.....SEDLRKLIV
HrobNb    .....GDRLPP..............PALHELEKLFGHL.........DSSNQHSFE.....TKDLVKLIK
HaztNb    ............................HNALKVPVHL.........DHKNPTRFE.....VEDLRKLIK
TurtNb    GSPNAAKM....................DKHMSKFEHL.........DHKNPHSFE.....VQDLHKLII
PtepNb    PSYKGLDR....................KNGVKIPKHL.........DIDNPHSFE.....MEDLKKLIQ
SmimNb    ............................RKGVKIPKHL.........DIENPHSFE.....IEDLKKLIQ
GoccNb    .....RTH........................VINDHV.........DKDNHHTFE.....IDDLKKLIQ
DsuzNb    ............................RKHLKVSQHL.........DHDNEHTFE.....IEDLRKLIQ
DmelNb    ............................RKHLKVSQHL.........DHDNEHTFE.....IEDLRKLIQ
AaegNb    ............................REHLKIAEHL.........DHENQHTFE.....IDDLKKLIL
CtelNb    .....MALSPAAAAARGIDSAFQEKAGILDSDKHLKHI.........DHANSNTFE.....VEDLAKLIQ
OvicNb    ...RRMDQ....................KALEGMVGHV.........DPATMDKFT.....DTDFEKLIK
SkowNb    ....RMDK....................KSITDLLGHV.........DLGNTKSFE.....EEDLAKLIQ
SpurNb    .....VSQ....................THLMDMIAHI.........DFGNEHTFE.....EQDLNNLLQ
NvecNb    .................................EREYWNPL.....FDDENPDFFG.....ADDFKKLLW
EpalNb    .................................EREHWNPV.....FDDDNPDFFG.....PDDFKKLLA
OfavNb    .................................EREYWNPL.....FDDDNPNFFG.....PEDFKKLLR
HvulNb    .....RNY..........................WNPI.........HHDNKDSFE.....MEDLKKLLS
CbriNb    ............................AHNVQMPDHL.........DVQELEKFH.....KEDLRKLIQ
CbreNb    ............................AHNIKMPDHL.........DIQQLEKFH.....KEDLRKLIQ
HconNb    ............................AHNVKVPEHL.........DASNWERFN.....KEDLRKLIR
```

```
TcanNb  ............................AHNIKVPEHI.........DVNDWEKFG.....KEDLRKLIV
SratNb  ............................PHNIKAPEHI.........DIDQLDKFH.....AEDLRKLVK
ShaeNb  ...................................GGI.........VYNSIPGYE.....TESPKKVVD
SmanNb  ............................DSHRNTMGGV.........VHKTLPGYE.....AESPNKIVD
SjapNb  ......................................................................
AqueNb  .....IEI....................RRQRERGRHLS......................DKEMEEKIR
SrosNb  ............................RPHDAMPEGG.........EEEDEDLFLDPEAAFDKLTEIVK
BdenNb  ......................................................................
BsalNb  ......................................................................
CcucNb  ......................................................................
ArepNb  ......................................................................
McirNb  ......................................................................
MbreNb  ............................PERPDQ..........AKFGDKGEGSHN.....EALVRNVVE
DqueNb  ...............................................GHDGPAEGE..............
FpinNb  ...............................................GHDGPASGE..............
FradNb  ...............................................DHDAPASGE..............
GfroNb  ...............................................GHDGPASGE..............
ShirNb  ...............................................EESKEPLTG..............
PostNb  ...............................................EHSGPSQGE..............
RsolNb  ......................................................................
NirrNb  ......................................................................
ScerNb  ......................................................GL..............
VpolNb  ..................................................RPPEGL..............
TphaNb  ..................................................TPPEGL..............
CglaNb  ..................................................KPPNGM..............
LmirNb  ..................................................KPPEGQ..............
EgosNb  ..................................................RPPEGK..............
LngbNb  ..............................................YDCDRDGFLV.....CKDFENIAT
McolNb  ..............................................YDANCDGALV.....CQDFENLVK
CchtNb  ..............................................HDAKFDGALA.....YQDFENIIK
SmajNb  ..............................................YDVNCDGFIV.....EQDFEDVAA
NnodNb  ..............................................YDSDYTGVLV.....KKDFELMFD
MproNb  ..............................................YDVDGSGVIT.....EADYEKMAQ
PsojNb  ..............................................IDTDGNGVIT.....VSELAEALR
PhalNb  ..............................................IDTDGNGVIT.....VSELAEALR
PinsNb  ..............................................IDKDGNGVIT.....VAELADALR
AcanNb  ..............................................IDADQNGVIT.....ITELATALR
TclaNb  ..............................................IDADGNGVIT.....MQELAAAVR
EsilNb  ..............................................IDADNNGSIC.....VDELKMVVK
EguiNb  ..............................................MDTDNSGAIT.....FDELKEGLK
DdisNb  ..............................................LDKDNNGKLT..............
AsubNb  ..............................................LDNNNDGKLS..............
AcasNb  ..............................................LDANGDGHLT..............
LocuNb  ..............................................LDTNKDGKVD..............
CcarNb  ..............................................LDANKDGKVD..............
SansNb  ..............................................LDANKDGKVD..............
GproNb  ......................................................................
RgloNb  ...........................LPPRMNLPRGLSAVKTLAASGSGLQSTMSM..SSVSNIQPLED
AbacNb  ...........................................T..........................
TgonNb  ...........................................T..........................
HhamNb  ...........................................T..........................
BbesNb  ...........................................N..........................
CsuiNb  ...........................................D..........................
CcayNb  ...........................................T..........................
TbruNb  AEATRLLCMSEARAKALKAECVANTMKARQLQRVCDGL........RGSGKKDESE...VF.RDMRLLIE
LbraNb  AGVLRLLHLQENEIRKLKSQNSTTKLQLRSVERECAVL........RGFRTENELS...ALKEDLRFLIE
LseyNb  GGVLKLLHMRGNEVKTLSNQHHTATLQLRYIQRECAAL........RGLRNEDETS...LLKKDLSYLIE
PfalNb  TGINNVYHNIEDNQKLYTDVLI..IGAGISGLAASYYLNKCNFANCTEFNNEKSMRKEKTKKKKYKVIVK
```

```
          160       170       180       190       200       210       220
HsapNb1  TATRDLAQYDAAHHEEFKRYEMLKEHERRRYLESLGEEQRKEAERKLEEQQRRHREHPK.VNVPGSQA..
PtroNb1  TATRDLAQYDAAHHEEFKRYEMLKEHERRRYLESLGEEQRKEAERKLEEQQRRHREHPK.VNVPGSQA..
PpanNb1  TATRDLAQYDAAHHEEFKRYEMLKEHERRRYLESLGEEQRKEAERKLEEQQRRHREHPK.VNVPGSQA..
PabeNb1  TATRDLAQYDAAHHEEFKRYEMLKEHERRRYLESLGEEQRKEAERKLEEQQRRHREHPK.VNVPGSQA..
GgorNb1  TATRDLAQYDAAHHEEFKRYEMLKEHERRRYLESLGEEQRKEAERKLEEQQRRHREHPK.VNVPGSQA..
CatyNb1  TATRDLAQYDAAHHEEFKRYEMLKEHERRRYLESLGEEQRKEAERKLEEQQRRHREHPK.VNVPGSQA..
MmulNb1  TATRDLAQYDAAHHEEFKRYEMLKEHERRRYLESLGEEQRKEAERKLEEQQRRHREHPK.VNVPGSQA..
RroxNb1  TATRDLAQYDAAHHEEFKRYEMLKEHERRRYLESLGEEQRKEAERKLEEQQRRHREHPK.VNVPGSQA..
CangNb1  TATRDLAQYDAAHHEEFKRYEMLKEHERRRYLESLGEEQRKEAERKLEEQQRRHREHPK.VNVPGSQA..
MfasNb1  TATRDLAQYDAAHHEEFKRYEMLKEHERRRYLESLGEEQRKEAERKLEEQQRRHREHPK.VNVPGSQA..
MnemNb1  TATRDLAQYDAAHHEEFKRYEMLKEHERRRYLESLGEEQRKEAERKLEEQQRRHREHPK.VNVPGSQA..
CcapNb1  TATRDLAQYDAAHHEEFKRYEMLKEHERRRYLESLGEEQRKEAERKLEEQQRRHREHPK.VNVPGSQA..
OgarNb1  TATRDLAQYDAAHHEEFKRYEMLKEHERRRYLESLGEEQRKEAERKLEEQQRRHREHPK.VNVPGSQA..
PcoqNb1  TATRDLAQYDAAHHEEFKRYEMLKEHERRRYLESLGEEQRKEAERKLEEQQRRHREHPK.VNVPGSQA..
GvarNb1  TATRDLAQYDAAHHEEFKRYEMLKEHERRRYLESLGEEQRKEAERKLEEQQRRHREHPK.VNVPGSQA..
TchiNb1  TATRDLAQYDAAHHEEFKRYEMFKEHERRRYLESLGEEQRKEAERKLEEQQRRHREHPK.VNVPGSQA..
BmutNb1  TATRDLAQYDAAHHEEFKRYEMLKEHERRRYLESLGEEQRKEAERKLEEQQRRHREHPK.VNVPGSQA..
MjavNb1  TATRDLAQYDAAHHEEFKRYEMLKEHERRRYLESLGEEQRKEAERKLEEQQRRHREHPK.VNVPGSQA..
VpacNb1  TATRDLAQYDAAHHEEFKRYEMLKEHERRRYLESLGEEQRKEAERKLEEQQRRHREHPK.VNVPGSQA..
CbacNb1  TATRDLAQYDAAHHEEFKRYEMLKEHERRRYLESLGEEQRKEAERKLEEQQRRHREHPK.VNVPGSQA..
VpacNb1  TATRDLAQYDAAHHEEFKRYEMLKEHERRRYLESLGEEQRKEAERKLEEQQRRHREHPK.VNVPGSQA..
CferNb1  TATRDLAQYDAAHHEEFKRYEMLKEHERRRYLESLGEEQRKEAERKLEEQQRRHREHPK.VNVPGSQA..
HarmNb1  TATRDLAQYDAAHHEEFKRYEMLKEHERRRYLESLGEEQRKEAERKLEEQQRRHREHPK.VNVPGSQA..
RsinNb1  TATRDLAQFDAAHHEEFKRYEMLKEHERRRYLESLGEEQRKEAERKLEEQQRRHREHPK.INVPGSQA..
JjacNb1  TATRDLAQYDEAHHEEFKRYEMLKEHERRRYLESLGEEQRKEAERKLEEQQRRHREHPK.VNVPGSQA..
DordNb1  TATRDLAQYDAAHHEEFKRYEMLKEHERRRYLESLGEEQRKEAERKLEEQQRRHREHPK.VNVPGSQA..
NgalNb1  TATRDLAQFDAAHHEEFKRYEMLKEHERRRYLESLGEEQRKEAERKLEEQQRRHREHPK.VNVPGSQA..
RnorNb1  TATRDLAQYDAAHHEEFKRYEMLKEHERRRYLESLGEEQRKEAERKLQEQQRRHREHPK.VNVPGSQA..
MmusNb1  TATRDLAQYDAAHHEEFKRYEMLKEHERRRYLESLGEEQRKEAERKLQEQQRRHREHPK.VNVPGSQA..
MaurNb1  TATRDLAQYDAAHHEEFKRYEMLKEHERRRYLESLGEEQRKEAERKLQEQQRRHREHPK.VNVPGSQA..
CasiNb1  TATRDLAQYDAAHHEEFKRYEMLKEHERRRYLESLGEEQRKEAERKLEEQQRRHREHPK.VNVPGSQA..
NparNb   AATKDLENYDAARHEEFKRYEMMKEHERREYLKSLDEDKRKMEEAHYEELKKKHREHPK.VNVPGSKD..
XlaeNb1  AATKDLENYDAARHEEFKRYEMMKEHERREYLKSLDEEKRKMEEAHYEEMKKKHREHPK.VNVPGSMD..
XtroNb1  AATKDLENYDAARHEEFKRYEMMKEHERREYLKSLDEEKRKMEEAHFEEMKKKHKEHPK.VNVPGSID..
PmucNb1  AATKDLENYDAAHHEEFKRYEMMKEHERREYLKSLDEEKRREEEAHFEELKKKHKEHPK.VNVPGSRD..
PbivNb1  AATKDLENYDAAHHEEFKRYEMMKEHERREYLKSLDEEKRREEEAHFEELKKKHKEHPK.VNVPGSRD..
GjapNb1  TATKHLENFDADHHNEFKRYEMMKEHERREYLKSLDEEKRREEEARFEELKKKHKEHPK.VNAPGSQD..
DrerNb1  TATKDLENYDAERHEEFKRYEMLKEHERREYLKSLDQEKREKEEKRMDELKEKHRQHPK.VNSPGSVD..
TrubNb1  TATKDLENYDAERHEEFKRYEMLKEHERREYLKSLDQEKREHEEKRLQELKEKHRQHPK.VNAPGSVA..
TnigNb1  T.VRSRTMTLCQRHEEFKRYEMLKEHERREYLRSLDQEKREQEEKRLQELREKHRQHPK.VNAPGSVA..
IpunNb   SATKDLENYDKERHEEFKRYEMMKEHERREHLKTLDEEGRKKEEEHYEEMKKKHADHPK.INHPGSQN..
DrerNb2  SATKDLENYDKERHEEFKKYEMMKEHERREHLKTLDEDGRKKEEEHYEEMKKKHADHPK.VNHPGSKD..
TrubNb2  SATNDLENYDKERHDEFKRYEMMKEHNRQEHLKTLDNEERRKEEEHYEEMKKKHADHPK.INHPGSQN..
HsapNb2  AATSDLEHYDKTRHEEFKKYEMMKEHERREYLKTLNEEKRKEEESKFEEMKKKHENHPK.VNHPGSKD..
PpanNb2  AATSDLEHYDKTRHEEFKKYEMMKEHERREYLKTLNEEKRKEEESKFEEMKKKHENHPK.VNHPGSKD..
GgorNb2  AATSDLEHYDKTRHEEFKKYEMMKEHERREYLKTLNEEKRKEEESKFEEMKKKHENHPK.VNHPGSKD..
CatyNb2  AATSDLEHYDKTRHEEFKKYEMMKEHERREYLKTLNEEKRKEEESKFEEMKKKHENHPK.VNHPGSKD..
CangNb2  .....................MMKEHERREYLKTLNEEKRKEEESKFEEMKKKHENHPK.VNHPGSKD..
RroxNb2  AATSDLEHYDKTRHEEFKKYEMMKEHERREYLKTLNEEKRKEEESKFEEMKKKHENHPK.VNHPGSKD..
PabeNb2  AATSDLEHYDKTRHEEFKKYEMMKEHERREYLKTLNEEKRKEEESKFEEMKKKHENHPK.VNHPGSKD..
MmulNb2  AATSDLENYDKTRHEEFKKYEMMKEHERREYLKTLNEEKRKEEESKFEEMKKKHENHPK.VNHPGSKD..
MfasNb2  AATSDLENYDKTRHEEFKKYEMMKEHERREYLKTLNEEKRKEEESKFEEMKKKHENHPK.VNHPGSKD..
CcapNb2  AATSDLEHYDKTRHEEFKKYEMMKEHERREYLKTLSEEKRKEEESKFQEMKKKHENHPK.VNHPGSKD..
PtroNb2  AATSDLEHYDKTRHEEFKKYEMMKEHERREYLKTLNEEKRKEEESKFEEMKKKHENHPK.VNHPGSKD..
OgarNb2  AATSDLEHYDKTRHEEFKKYEMMKEHERREYLKTLNEEKRKEEESRFEEMKKKHENHPK.VNHPGSKD..
PcoqNb2  AATSDLEDYDKTRHEEFKKYEMMKEHERREYLKTLNEEKRKEEESKFEEMKKKHENHPK.VNHPGSKD..
TchiNb2  AATNDLEHYDKTRHEEFKKYEMMKEHERREYLKTLSEEKRKEEESKFEEMKKKHENHPK.VNHPGSKD..
MjavNb2  AATSDLEHYDKTRHEEFKKYEMMKEHERREYLKTLNEEKRKEEESKFEEMKKKHENHPK.VNHPGSKD..
HarmNb2  AATSDLEHYDKTRHEEFKKYEMMKEHERREYLKTLNEEKRKEEESKFEEMKKKHENHPK.INHPGSKD..
RsinNb2  AATSDLEHYDKTRHEEFKKYEMMKEHERREYLKTLSEEKRKEEESKFEEMKKKHENHPK.INHPGSKD..
DordNb2  AATSDLEHFDKTRHEEFKKYEMMKEHERREYLKTLNEEKRKEEESKFEEMKKKHEDHPK.VNHPGSKD..
BmutNb2  AATSDLEHYDKARHEEFKKYEMMKEHERREYLKTLSEEKRKEEESKFAEMKKKHENHPK.VNHPGSKD..
VpacNb2  AATSDLEHYDKARHEEFKKYEMMKEHERREYLKTLNEEKRKEEESKFEEMKKKHENHPK.VNHPGSKD..
CferNb2  AATSDLEHYDKARHEEFKKYEMMKEHERREYLKTLNEEKRKEEESKFEEMKKKHENHPK.VNHPGSKD..
CbacNb2  AATSDLEHYDKARHEEFKKYEMMKEHERREYLKTLNEEKRKEEESKFEEMKKKHENHPK.VNHPGSKD..
JjacNb2  AATSDLENYDKTRHEEFKKYEMMKEHERREYLKTLNEEKRKEEESKFEEMKKKHENHPK.VNHPGSKD..
CasiNb2  AATSDLEHYDKTRHEEFKKYEMLKEHERREYLKTLNEEKRKEEESKYEEMKKKHENHPK.VNHPGSKD..
NgalNb2  AATTDLEHYDKTRHEEFKKYEMMKEHERREYLKTLNEEKRKEEEAKFAEMKKKHEDHPK.VNHPGSKD..
RnorNb2  AATADLEQYDRTRHEEFKKYEMMKEHERREYLKTLSEEKRKEEEAKFAEMKRKHEDHPK.VNHPGSKD..
MmusNb2  AATADLEQYDRTRHEEFKKYEMMKEHERREYLKTLSEEKRKEEESKFEEMKRKHEDHPK.VNHPGSKD..
MaurNb2  AATADLEQYDRTRHEEFRKYEMMKEHERREYLKTLNEEKRKEEESKFEEMKRKHENHPK.VNHPGSKD..
PbivNb2  AATNDLENYDKARHEEFKKYEMMKEHERREYLKTLDEEKRHQEEAKYEEMKKKHSNHPK.VNHPGSKD..
PmucNb2  TATHDLENYDNDRHEEFKKYEMMKEHERREYLKTLDEEKRHQEEAKYEDLKKKHNDHPK.VNHPGSKD..
GjapNb2  TATSDLENYDKARHDEFKRYEMMKEHERREYLKTLDEEKRHQEESKYEEMKKKHGEHPK.VNHPGSKD..
LchaNb   TATNDLENYDQERHEEFKNYEMKKEHERREYLKTLDEEKRKEEETRYEEMRKKHSEHPK.VNHPGSKD..
XlaeNb2  TATKDLDSYDKTRHDEFKRYEMMKEHERREYLKTLDEEKRKEEEEKYEEMKKKHKEHPK.MNHPGSKD..
XtroNb2  TATKDLENYDKTRHDEFKRYEMMKEHERREYLKTLDEEKRKEEEEKYEEMKKKHREHPK.INHPGSKD..
AcalNb   KTTSDLEELDKKRRGEFKEYEMEKEFEKQEHLRELPEEERKKEEHRLEELKKKHAEHPK.LNHPGSRD..
BglaNb   KTTSDLEELDKQRKEEFKEYEMEKTFEQQEHLKALKEEERKREEARLEELKKKHAQHPK.VNHPGSKD..
LgigNb   KTTNDLEELDKQRRDEFKEYEMEKQYEEKEKLKHLSEEERKKEEERLAELKKKHGEHEK.INHPASKD..
DmagNb   QTSKDLEEADRQRKEEFKEYEMQKEFEHQNKLKTLDEEKRKAEQKAWEDAQAKHKQHPK.MHHPGSRQ..
DpulNb   QTSKDLEEADKQRKEEFKEYEMQKEFEHQNKLKGLDEEKRKTEEKEWEEAQAKHKQHPK.MHHPGSKQ..
HrobNb   RTTDDLEDIDRRRREDFKRYEMEKEHLRREEMKGLPTEEREKKEKEHEEMEKKHREHPK.VHHPGNEE..
HaztNb   QTSDDLEKMDAQRRKDFSQYEMQKEFEYKQQLEHMTAEERKKAEEEHEMRIKKHKDHPK.LHHPGSKQ..
TurtNb   KATRDLEELDKKRREEFKKYEMEKEHRYRESLKNMTEEQKTEAVKKHSEMNQKHKEHPR.VHHPGSKQ..
PtepNb   SATTDLEEIDRQRREEFKTYELEKEAAYRESLQNLTSEEKKKAEEHHKELIEKHKDHPK.VHHPGSKP..
SmimNb   TATKDLEEIDRQRREEFKKYELEKEAKYRESLRNMTEEQRKETEKHHEEMIQKHKDHPK.VHHPGSKP..
GoccNb   AATRDLEKLHEAQKEEFKNYEMEKEFEFQQGIKNMTEDQKKDELKKHEE..EKHRKHPS.AHHPGSKA..
DsuzNb   KTSDDLAEADRKRRGEFKEYEMQKEFEREAQKKEMDEESRKKFEAEVKEKEQKHKDHEK.LHHPGNKA..
DmelNb   KTSDDLAEADRKRRGEFKEYEMQKEFEREAQKKEMDEESRKKFETELKEKEEKHKDHEK.LHHPGNKA..
AaegNb   KTSQDLSENDRRRREEFKEYELQKEFEKQEKLRLLDEEHRKQYEEDLKKQKETHEKHEK.IHHPGNKA..
CtelNb   QATTDLEDLDKERRSDFKTYEMEKEHLQKEELKALDEQARAAKEKELHDKKEKHKKHAK.VNHPGNME..
OvicNb   KAAADLDEADQERRKEFKRYEMEKELKRRQDLKTLDQAAQVKAEAEFNERRDRLKSHPN.IKHPGSKA..
SkowNb   KATKDLDEMDRRRKTDFKKYEMEKEIKRREKLKQMSDAKREQAKQEWEESQKKRKQHDK.VKHPGSKK..
SpurNb   KATKDLEENDRERRKEFKRYEMEKELKRRSEMRRMNEEERAEAMENYEKERKEMKQHAK.LNHPGSKK..
NvecNb   KHHEEMDKQDRERREEFKKHEMQKEHERKIKMKDMDEKHRKEAEEHFRELQEKHHNQSKQLHHPGSKA..
EpalNb   KHHNEMDKRDKERKEDYKKYEMEKEHKRREKLKEMDEAHRKEAEEKFKKEQERHXEEAKKLHHPGSKA..
OfavNb   KHHEEMDKQDNERRQEFKKHEMEKEHKRKEHLKEMDEKARKEEELKFEALRQKHLNASKNLHHPGSKA..
HvulNb   KHNDMMSAQDAKRHEEFKAYEMEKEHERREKLHNMTAEERAKEEALYKEHREQRSKHEK.IHEPGHKA..
CbriNb   KTVADMNKMDDQRKEDFKQYEMKKQAEEDHKLAQMSPEEREKAKREHEEAEKRHNDHEK.LKHPGSKD..
CbreNb   KTVADMNAYDEIRKEEFKQYEMKKQAEEDHRLAQMTAEEREKAKQEHEESVKRHNDHEK.LKHPGSKD..
HconNb   RTVEDMEIIDRNRQKDFKEYEMRKKAEEDLKLAKMTEEERKKAIHDAEEAKKRHNEHEK.LKHPGSRD..
```

```
TcanNb   KTVADMEKIDEQRREQFKEYEMQKKAEADHKMANMTPEERKKFEQDLEQMKKRHNEHEK.VKHPGSRD..
SratNb   KTFEDMDEIDRQRKEKFKKYEMEKRAKYDHELAKLPEEERKKLQEEHEAAKKRHQDHEK.LNHPGERR..
ShaeNb   Q....LKKLDTDQEKKYQEHVAKEKSDMVRKLSKFTEEERARMLKSLKEQAEKHKSHPR.IYQPGSQD..
SmanNb   K....LKKLDDEQERKYQEHVAKEKSDMVQKFSKFTEEERARMLKSLKELAEKHKRHPR.IHQPGSQD..
SjapNb   ...........................................LRHFKESAERHRRRPK.IYQPGSED..
AqueNb   EHQRIQEQLDHARREDFKNFEMKLEHRRRQKLKEMDETQRLREEARLKKLSDERKVHEP.LLHPASKD..
SrosNb   RRAAVLGEIDTKRRAAFLQKEMMRELEFRRSVAHEKDPEKKK......QLIKSHKQ......KIGHAK..
BdenNb   ..........YGEDSNYGDVKPSASHF.RERLEAIGYGQD....................SNEKG.....
BsalNb   ..........YGESTNYGDVSPSAAHF.SERLRAAGYGQDI...................PTESS.....
CcucNb   ............DHSNYGDSKQNKEHI.REHLRKLSASSG..........................
ArepNb   ............DHSNYGSEKTNRQHI.QEHLDILAKDG.............................
McirNb   ............PHSNYGDEKKNKHHI.QEHMKAMSKDDD............................
MbreNb   KRVRLMGQLDERSRQAFVEHMMQRELEFRERMKATTNAADRRRLNEEHQKAWADLKKRR.DFQPGHRD..
DqueNb   ............TIQQYAERHMAKEHH...........................................
FpinNb   ............TIQQYAERHMAKEHH...........................................
FradNb   ............SIQQYAERHMAKEHH...........................................
GfroNb   ............TVQEYAQKHMASEHH...........................................
ShirNb   ............DAAQYAQRHMATEHH...........................................
PostNb   ............TIQQYAQRHMSSEHH...........................................
RsolNb   ..................WLQMASEHH...........................................
NirrNb   ............SAADWENMHMQDEHN...........................................
ScerNb   ...............SWEEWHMDHEHQ...........................................
VpolNb   ...............SWEEWHMQHEHQ...........................................
TphaNb   ...............SWEEWHMQHEHQ...........................................
CglaNb   ...............SWEEWHMDHEHE...........................................
LmirNb   ...............EWEDWHMQHEHQ...........................................
EgosNb   ...............NWEQWHMEHEHQ...........................................
LngbNb   KLAKVKNLGARSPKVLALKERFLQAWKSLLSGADSNSDKKI.............................
McolNb   KLADLRNWGSRSPKYQVMLNRCMDDWKSLKGDADTSQDKKV.............................
CchtNb   KLAALRHWGSRSPRYQILLNNCNRNWEELKRDADITHNKKV.............................
SmajNb   KLADRAGWSVRSPKGITLKNQLAQDWQGLQKASDTNRDRQI.............................
NnodNb   KLSTLRNWSLRSPRCLVLQDKLMRKWQGLEKKADTAKNKQV.............................
MproNb   SQAKVQGYQPGSLEYNIICSQFKTLWKNLQKEVDINDDGEV.............................
PsojNb   D.....TGLGMIEEEVL..........ELVKGIDIDGDGLV.............................
PhalNb   D.....TGLGMIEEEVL..........DLVKGIDIDGDGLV.............................
PinsNb   S.....MGHGMIEEEVM..........ELLQGIDIDGDGLV.............................
AcanNb   G.....MGHGMIETEVL..........ELLQGIDIDGDGLI.............................
TclaNb   D.....MGHGMLESEVL..........QLLKGLDIDGDGNL.............................
EsilNb   KEVGRDARARGVLEPWTASSKAVTGYKRLLMSIQLSVPMQC.............................
EguiNb   R.....YGSTLKESEIR..........ALMDAADVDNSGTI.............................
DdisNb   ..................REEIKEGFF.................................LRIPSSEK..
AsubNb   ..................RDELKQGFR.................................LHIPSSER..
AcasNb   ..................RDELHDGLK.................................LKLPATEA..
LocuNb   ..................VSELKEGLE.................................MGTTLGKG..
CcarNb   ..................VSELKTGLA.................................MGFSMGKG..
SansNb   ..................VSELKTGLA.................................MGFTMGKG..
GproNb   ......................................................................
RgloNb   VHITLVDRIYFYSSFNYSSFHANRAAP...............QGGMGADYMKANGANPAASNKPQETSTY
AbacNb   .........................................................PR.KSLPGVPDG.
TgonNb   .........................................................QE.DLLPGLPDKK
HhamNb   .........................................................PK.DVLPGLSDKK
BbesNb   .........................................................PD.DCQPGLPDQK
CsuiNb   .........................................................LE.AC..GIPDKN
CcayNb   .........................................................PK.ASIPDIPDSA
TbruNb   A.................NRQMEKDCQLRVALNMGPKQKTVKAAQRVDASQKQTKTLPQFMSQPGSFP..
LbraNb   A.................NKTLEQENALRRHLNARDAA......................HEPGVMD..
LseyNb   A.................NKKLEAENQLRARQQVVEASP.................PQ..LQPSTAD..
PfalNb   KRKYNYNGNDKNKMGNFSIYDVQKEYVKEEYSKLINNNQRYYSDEDYRRIKNRRKIKLRSVLYPTNKSN.
```

```
                230         240                                             250       260
HsapNb1   ..QLKEVWEELDGL..DPNRFNP...........................KTFFILHDINSDGVLDEQE
PtroNb1   ..QLKEVWEELDGL..DPNRFNP...........................KTFFILHDINSDGVLDEQE
PpanNb1   ..QLKEVWEELDGL..DPNRFNP...........................KTFFILHDINSDGVLDEQE
PabeNb1   ..QLKEVWEELDGL..DPNRFNP...........................KTFFILHDINSDGVLDEQE
GgorNb1   ..QLKEVWEELDGL..DPNRFNP...........................KTFFILHDINSDGVLDEQE
CatyNb1   ..QLKEVWEELDGL..DPNRFNP...........................KTFFILHDINSDGVLDEQE
MmulNb1   ..QLKEVWEELDGL..DPNRFNP...........................KTFFILHDINSDGVLDEQE
RroxNb1   ..QLKEVWEELDGL..DPNRFNP...........................KTFFILHDINSDGVLDEQE
CangNb1   ..QLKEVWEELDGL..DPNRFNP...........................KTFFILHDINSDGVLDEQE
MfasNb1   ..QLKEVWEELDGL..DPNRFNP...........................KTFFILHDINSDGVLDEQE
MnemNb1   ..QLKEVWEELDGL..DPNRFNP...........................KTFFILHDINSDGVLDEQE
CcapNb1   ..QLKEVWEELDGL..DPNRFNP...........................KTFFILHDINSDGVLDEQE
OgarNb1   ..QLKEVWEELDGL..DPNRFNP...........................KTFFLMHDINSDGVLDEQE
PcoqNb1   ..QLKEVWEELDGL..DPNRFNP...........................KTFFILHDINSDGVLDEQE
GvarNb1   ..QLKEVWEELDGL..DPNRFNP...........................KTFFLLHDINSDGVLDEQE
TchiNb1   ..QLKEVWEELDGL..DPNRFNP...........................KTFFILHDINSDGVLDEQE
BmutNb1   ..QLKEVWEELDGL..DPNRFNP...........................KTFFILHDINSDGVLDEQE
MjavNb1   ..QLKEVWEELDGL..DPNRFNP...........................KTFFILHDINSDGVLDEQE
VpacNb1   ..QLKEVWEELDGL..DPNRFNP...........................KTFFILHDINSDGVLDEQE
CbacNb1   ..QLKEVWEELDGL..DPNRFNP...........................KTFFILHDINSDGVLDEQE
VpacNb1   ..QLKEVWEELDGL..DPNRFNP...........................KTFFILHDINSDGVLDEQE
CferNb1   ..QLKEVWEELDGL..DPNRFNP...........................KTFFILHDINSDGVLDEQE
HarmNb1   ..QLKEVWEELDGL..DPNRFNP...........................KTFFILHDINSDGVLDEQE
RsinNb1   ..QLKEVWEELDGL..DPNRFNP...........................KTFFILHDINSDGVLDEQE
JjacNb1   ..QLKEVWEELDGL..DPNRFNP...........................KTFFILHDINSDGVLDEQE
DordNb1   ..QLKEVWEELDGL..DPNRFNP...........................KTFFILHDINSDGILDEQE
NgalNb1   ..QLKEVWEELDGL..DPNRFNP...........................KTFFILHDINSDGVLDEQE
RnorNb1   ..QLKEVWEELDGL..DPNRFNP...........................KTFFILHDINSDGVLDEQE
MmusNb1   ..QLKEVWEELDGL..DPNRFNP...........................KTFFILHDINSDGVLDEQE
MaurNb1   ..QLKEVWEELDGL..DPNRFNP...........................KTFFILHDINSDGVLDEQE
CasiNb1   ..QLKEVWEELDGL..DPNRFNP...........................KTFFILHDINSDGVLDEQE
NparNb    ..QLKEVWEETDGL..DPNDFNP...........................KTFFNLHDTNGDGVLDEQE
XlaeNb1   ..QLKEVWEETDGL..DPNEFNP...........................KTFFKLHDTNGDGVLDEQE
XtroNb1   ..QLKEVWEETDGL..DPNEFNP...........................KTFFKLHDTNGDGVLDEQE
PmucNb1   ..QLKEVWQETDGL..DPNEFNP...........................KTFFKLHDTNSDGVLDEQE
PbivNb1   ..QLKEVWQETDGL..DPSEFNP...........................KTFFKLHDTNSDGVLDEQE
GjapNb1   ..QLKEVWQETDGL..DPNEFNP...........................KTFFNLHDTNSDGVLDEQE
DrerNb1   ..QLREVWEETDGL..DPQEFNP...........................KTFFKLHDTNSDGVLDVQE
TrubNb1   ..QLREVWEETDGL..DPQEFNP...........................KTFFKLHDTNDDKVLDEQE
TnigNb1   ..QLQEVWEETDGL..DPLEFNP...........................KTFFKLHDTNDDKVLDEQE
IpunNb    ..QLKEVWEEADGL..DPDDFDP...........................KTFFNLHDTNGDGFFDEQE
DrerNb2   ..QLKEVWEEADGL..DPEDFDP...........................KTFFNLHDTNGDGFFDEQE
TrubNb2   ..QLKEVWEEADGL..DPDDFDP...........................KTFFKLHDTNGDGFFDEQE
HsapNb2   ..QLKEVWEETDGL..DPNDFDP...........................KTFFKLHDVNSDGFLDEQE
PpanNb2   ..QLKEVWEETDGL..DPNDFDP...........................KTFFKLHDVNSDGFLDEQE
GgorNb2   ..QLKEVWEETDGL..DPNDFDP...........................KTFFKLHDVNSDGFLDEQE
CatyNb2   ..QLKEVWEETDGL..DPNDFDP...........................KTFFKLHDVNSDGFLDEQE
CangNb2   ..QLKEVWEETDGL..DPNDFDP...........................KTFFKLHDVNSDGFLDEQE
RroxNb2   ..QLKEVWEETDGL..DPNDFDP...........................KTFFKLHDVNSDGFLDEQE
PabeNb2   ..QLKEVWEETDGL..DPNDFDP...........................KTFFKLHDVNSDGFLDEQE
MmulNb2   ..QLKEVWEETDGL..DPNDFDP...........................KTFFKLHDVNSDGFLDEQE
MfasNb2   ..QLKEVWEETDGL..DPNDFDP...........................KTFFKLHDVNSDGFLDEQE
CcapNb2   ..QLKEVWEETDGL..DPNDFDP...........................KTFFKLHDVNSDGFLDEQE
PtroNb2   ..QLKEVWEETDGL..DPNDFDP...........................KTFFKLHDVNSDGFLDEQE
OgarNb2   ..QLKEVWEETDGL..DPNDFDP...........................KTFFKLHDVNSDGFLDEQE
PcoqNb2   ..QLKEVWEETDGL..DPDDFDP...........................KTFFKLHDVNSDGFLDEQE
TchiNb2   ..QLKEVWEETDGL..DPNDFDP...........................KTFFQLHDVNSDGFLDEQE
MjavNb2   ..QLKEVWEETDGL..DPNDFEP...........................KTFFKLHDVNNDGFLDEQE
HarmNb2   ..QLKEVWEETDGL..DPNDFDP...........................KTFFKLHDVNNDGFLDEQE
RsinNb2   ..QLKEVWEETDGL..DPNDFDP...........................KTFFRLHDVNNDGFLDEQE
DordNb2   ..QLKEVWEETDGL..DPNDFDP...........................KTFFKLHDVNGDGFLDEQE
BmutNb2   ..QLKEVWEEADGL..DPNDFDP...........................KTFFKLHDVNSDGFLDEQE
VpacNb2   ..QLKEVWEEADGL..DPNDFDP...........................KTFFKLHDVNSDGFLDEQE
CferNb2   ..QLKEVWEEADGL..DPNDFDP...........................KTFFKLHDVNSDGFLDEQE
CbacNb2   ..QLKEVWEEADGL..DPNDFDP...........................KTFFKLHDVNSDGFLDEQE
JjacNb2   ..QLKEVWEETDGL..DPNDFDP...........................KTFFRLHDVNNDGFLDEQE
CasiNb2   ..QLKEVWEEADGL..DPDDFDP...........................KTFFKLHDVNNDGFLDEQE
NgalNb2   ..QLKEVWEETDGL..DPNDFDP...........................KTFFKLHDVNSDGFLDEQE
RnorNb2   ..QLKEVWEETDGL..DPNDFDP...........................KTFFKLHDVNNDGFLDEQE
MmusNb2   ..QLKEVWEETDGL..DPNDFDP...........................KTFFKLHDVNNDGFLDEQE
MaurNb2   ..QLKEVWEEADGL..DPNDFDP...........................KTFFKLHDVNNDGFLDEQE
PbivNb2   ..QLKEVWEEADGL..DPNDFDP...........................KTFFKLHDVNADGFLDEQE
PmucNb2   ..QLKEVWEEADGL..DPNDFDP...........................KTFFKLHDVNADGFLDEQE
GjapNb2   ..QLKEVWEEADGL..DPNDFDP...........................KTFFKLHDVNNDGFLDEQE
LchaNb    ..QLKEVWEETDGL..DPNDFDP...........................KTFFKLHDTNSDGFLDEQE
XlaeNb2   ..QLKEVWEETDGL..DPSEFDP...........................KTFFKLHDTNSDGFLDEQE
XtroNb2   ..QLKEVWEETDGL..DPTDFDP...........................KTFFKLHDTNSDGFIDEQE
AcalNb    ..QFEEVWDKVDHL..EDQEFNP...........................KTFFFKHDLNGDMEWSIEE
BglaNb    ..QFEEVWEKVDHL..EDQEFNP...........................KTFFYTHDVNGDMEWSVDE
LgigNb    ..QFEEVWENTDHL..DKEDFDP...........................KTFFLLHDANGDGVWSIQE
DmagNb    ..QLEEVWEEQDHL..SPDSFNP...........................KTFFQLHDLDGNGFWDPDE
DpulNb    ..QLEEVWEEQDHL..SPDSFNP...........................KTFFALHDLDGNGYWDPDE
HrobNb    ..QLKDVWEEGDGL..ERDEFNP...........................RTFFYLHDTNSDGFLDPVE
HaztNb    ..QLEQVWEEEDHM..NKQEFNP...........................KTFFHLHDLDGNGHWDENE
TurtNb    ..QLEEVWEEQDRM..PKEEFNP...........................KTFFSLHDVNGDGYLDPEE
PtepNb    ..QLEQVWEEQDHL..PKDEFNL...........................RTFFAMHDLNGDGHLDYQE
SmimNb    ..QLEQVWEDQDHM..PKEEFNP...........................RTFFAMHDVNGDGHLDVEE
GoccNb    ..QLEQVWEEEDHM..EKQDFDP...........................KTFFAMHDLNGDGVLDEQE
DsuzNb    ..QLEDVWEKQDHM..DKNDFDP...........................KTFFSIHDVDSNGYWDEAE
DmelNb    ..QLEDVWEKQDHM..DKNDFDP...........................KTFFSIHDVDSNGYWDEAE
AaegNb    ..QLEEVWEKQDHM..DGQDFDP...........................KTFFMLHDIDGNGYWDENE
CtelNb    ..QLEEVWEETDNL..DRDDFNP...........................KTFFHLHDVDGDGYWDEQE
OvicNb    ..QMRDVWEETDKL..DPEDFNP...........................KTFFALHDTNNDKKLDMFE
SkowNb    ..QLEEVWEETDHL..DKDDFDP...........................RTFFMLHDKNGDGKLDPME
SpurNb    ..QLEEVWENTDHL..DRKDFNP...........................KTFLYLHDSNSDGFLDEFE
NvecNb    ..QLEQVWEESDGL..DAKDFDP...........................KTFFKLHDVNGDGFLDTGE
EpalNb    ..QLEQVWEESDGL..DKNEFEP...........................KTFFKLHDINGDGYLDTGE
OfavNb    ..QLEQVWEETDGL..DANDFEP...........................RTFFKLHDTNGDDYLDTGE
HvulNb    ..QLEETWEKEDGL..DPESFDP...........................RTFFNLHDKNSDHYLDLYE
CbriNb    ..QLQEVWEETDHL..EKDQYDP...........................KTFFALHDLNGDGFWNDFE
CbreNb    ..QLQEVWEESDHL..EKDQYDP...........................KTFFALHDLNGDGFWNDFE
HconNb    ..QLQEVWEDTDKM..DKDNFDP...........................RTFFALHDLNGDGYWNADE
```

```
TcanNb  ..QLEEVWEDSDKM..EKENFDP............................RTFFALHDLNGDGFWSAEE
SratNb  ..QLEEVWEEKDHM..DKENYDP............................KTFFALHDLNGDGYWNIQE
ShaeNb  ..QLREYWEKYEGL..DKESFNS............................RTLFADIDLDGDGYLNIHE
SmanNb  ..QLKEYWEKYEGL..DEQSFNS............................RTLFADIDLDGDGYLNIHE
SjapNb  ..QLKEYWEKYEGL..DRESFNP............................RTLFADIDLDGDGYLNVNE
AqueNb  ..QFEEVWKTDDGF..KDEKFDI............................RTFFHLHDTNGDGTLDIRE
SrosNb  ..QLNEVWEKQDGM...QSKFNP............................KVFFSLHDINGDDFWDHKE
BdenNb  .................DSNQEV............................IYFFSIHDYNYDGMLDGHE
BsalNb  .................THNQDV............................IYFFSIHDYNYDGKLDGHE
CcucNb  ..EEPQDLSDEDMI.....................................YYLFLIHDLNGDGFLDGHE
ArepNb  ..EKAAQVSEEDMI.....................................FYLFVLHDQNGDGHLDGHE
McirNb  ..DTVPELSEQDMI.....................................YYLFVIHDTNGDGYLDGHE
MbreNb  ..QLQEVWDNIDGM..KGQKFTP..............................................K
DqueNb  .................IDMFDP............................ASFFLLHDLNRDGIWDREE
FpinNb  .................IDSFDP............................PSFFQLHDLNRDGIWDREE
FradNb  .................IDSFDA............................SSFFQLHDLNRDGVWDREE
GfroNb  .................IDSFDV............................DSFFQLHDLNRDKFWDREE
ShirNb  .................IDSFDL............................ASFFQLHDLNRDGVWDKEE
PostNb  .................IDSFDI............................RSFFQLHDLNRDGMWDKEE
RsolNb  .................IDAFNP............................STFFQLHDLDRNGVLDRNE
NirrNb  .................IYAYDA............................NSFFTLHDLNNDGVWDSHE
ScerNb  .................LKXYTP............................ETFFALHDIKKKGFLDEND
VpolNb  .................MNKYDP............................ETFFALHDTKKKGYFDDSD
TphaNb  .................MKEYDP............................ESFFALHDIKKKGYFDDDD
CglaNb  .................LKKYTP............................EQFFALHDVKKKGYWDEND
LmirNb  .................MEKYDP............................ETFFALHDIGKKGYLDKKD
EgosNb  .................MAEYTP............................EKFFNLHDTNGKGYMDERD
LngbNb  ..SLQEWLTYYDDVLGNKDRYFK......................EVQSLMKLIFEVFDTNGDGQLCTEE
McolNb  ..NLEEWLNYYDAILSDEKKYEQ......................KVRFFMELVFDVFDGDEDGKISQQE
CchtNb  ..SLEEWLSYYSIILNDEEKYLH......................NIHFFMELVFEVFDKDEDGKISQAE
SmajNb  ..SLDEWLAYYDAVLADETLYNE......................RVKALTQLVFDVFDQDEDNTLSAAE
NnodNb  ..SLAEWLAYYDEGLSDTEPRSE......................DIFGLMELVFDVFDQDEDGKVNQKE
MproNb  ..TLEEFLEHKAKQLSFKEQYRPLWLERQSGIKTSQSYERSYEEDVIAKLTNLIFERLDVDGNGEISRQE
PsojNb  ..DYPEFLAATKRNLANQEEHLI.............................NAFNYFDTTNTGQITKAD
PhalNb  ..DYPEFLAATKRNLSNQKEHLI.............................NAFNYFDTTNTGHITKAD
PinsNb  ..DYPEFLAATKRNQTNKQEYLI.............................NAFNYFDTKKQGVITKAD
AcanNb  ..DYPEFLAATKRNLANKKEYLI.............................NAFNYFDTTKKGVITKAD
TclaNb  ..DYNEFLAATKRNLFNKEEYLV.............................NAFNYFDTKKQGVITKED
EsilNb  ..TAQEFLAATDRNVFIREDNVR.............................RAFQHFDIEGSGSITLAN
EguiNb  ..DYGEFIAATHLNKLEREEHLV.............................AAFSYFDKDGSGYITVDE
DdisNb  ..DIESFLTNVDKDKVSFKEFED..................FTIENIKKL.KIVFEELDTNKSGTLDIHE
AsubNb  ..NIDQFLEAVDSNHVSFDEFSQ..................FVNKNLVQL.KTLFNDLDTDKSGYLDIKE
AcasNb  ..DVDALLARLDIDKVSLLEFEA..................FAMAQSKLL.RKVFDDLDADKSGTIDVEE
LocuNb  ..AAQKIVSSGDRNKLDLKEFSR..................YLVEHEKKL.RLTFKSLDKNKDGCIDASE
CcarNb  ..EAQKIFASGDTDKLDFEEFSK..................YLKEHEKKL.RLTFKSLDKNQDGRIDARE
SansNb  ..EAQKMFVSGDTDKLDFEEFSK..................YLKEHEKKL.RLTFKSLDKNQDGRIDAKE
GproNb  ...................................................KFLFDLHDCDSDGHLNFSE
RgloNb  VVDLASIVHTLDIM..LKQPLNT....RL......................RFLFDLHDLDGDGFLDKNE
AbacNb  .D.................................................YVVFRFAS...........
TgonNb  VD.................................................FIVTRFAG...........
HhamNb  VD.................................................FIVTRFAG...........
BbesNb  VD.................................................YLVTRFAG...........
CsuiNb  VD.................................................YVVTRFAG...........
CcayNb  VD.................................................FVVSRFCG...........
TbruNb  ..DLDQSQR..DVV.MSSKAIYK..LERIQ........ER...........ASTLVMLDYDA.ACVGASK
LbraNb  ..D........D.L.MEEDDY..................R...........RKLMHMLDRDADAQADARE
LseyNb  ..S........DVI.LYDDAF..................R...........SEFLKMLDFDAAAPVGAAK
PfalNb  ..KLLKILSFNDT..RRRREYDKSLKPRVSLVCGKDNWE............STFYASDDTNEDKQINNNI
```

```
            270       280         290       300       310       320
HsapNb1  LEALFTKELEKVYDPKN..EEDDMRE..MEEERLRMREHVMKNVDTNQDRLVTLEEFLAST....QRKE.
PtroNb1  LEALFTKELEKVYDPKN..EEDDMRE..MEEERLRMREHVMKNVDTNQDRLVTLEEFLAST....QRKE.
PpanNb1  LEALFTKELEKVYDPKN..EEDDMRE..MEEERLRMREHVMKNVDTNQDRLVTLEEFLAST....QRKE.
PabeNb1  LEALFTKELEKVYDPKN..EEDDMRE..MEEERLRMREHVMKNVDTNQDRLVTLEEFLAST....QRKE.
GgorNb1  LEALFTKELEKVYDPKN..EEDDMRE..MEEERLRMREHVMKNVDTNQDRLVTLEEFLAST....QRKD.
CatyNb1  LEALFTKELEKVYDPKN..EEDDMRE..MEEERLRMREHVMKNVDTNQDRLVTLEEFLTST....QRKE.
MmulNb1  LEALFTKELEKVYDPKN..EEDDMRE..MEEERLRMREHVMKNVDTNQDRLVTLEEFLTST....QRKE.
RroxNb1  LEALFTKELEKVYDPKN..EEDDMRE..MEEERLRMREHVMKNVDTNQDRLVTLEEFLTST....QRKE.
CangNb1  LEALFTKELEKVYDPKN..EEDDMRE..MEEERLRMREHVMKNVDTNQDRLVTLEEFLTST....QRKE.
MfasNb1  LEALFTKELEKVYDPKN..EEDDMRE..MEEERLRMREH....VDTNQDRLVTLEEFLTST....QRKE.
MnemNb1  LEALFTKELEKVYDPKN..EEDDMRE..MEEERLRMREHVMKNVDTNQDRLVTLEEFLTST....QRKE.
CcapNb1  LEALFTKELEKVYDPKN..EEDDMRE..MEEERLRMREHVMKNVDTNQDRLVTLEEFLAST....QRKE.
OgarNb1  LEALFTKELEKVYDPKN..EDDDMRE..MEEERLRMREHVMKNVDTNQDRLVTLEEFLAST....QRKE.
PcoqNb1  LEALFTKELEKVYDPKN..EEDDMRE..MEEERLRMREHVMKNVDTNQDRLVTLEEFLAST....QRKE.
GvarNb1  LEALFTKELEKVYDPKN..EEDDMRE..MEEERLRMREHVMKNVDTNQDRLVTLEEFLAST....QRKE.
TchiNb1  LEALFTKELEKVYDPKN..EEDDMRE..MEEERLRMREHVMKNVDTNQDRLVTLEEFLAST....QRKE.
BmutNb1  LEALFTKELEKVYDPKN..EDDDMRE..MEEERLRMREHVMKNVDTNQDRLVTLEEFLAST....QRKE.
MjavNb1  LEALFTKELEKVYDPKN..EEDDMRE..MEEERLRMREHVMKNVDTNQDRLVTLEEFLAST....QRKE.
VpacNb1  LEALFTKELEKVYDPKN..EEDDMRE..MEEERLRMREHVMKNVDTNQDRLVTLEEFLAST....QRKE.
CbacNb1  LEALFTKELEKVYDPKN..EEDDMRE..MEEERLRMREHVMKNVDTNQDRLVTLEEFLAST....QRKE.
VpacNb1  LEALFTKELEKVYDPKN..EEDDMRE..MEEERLRMREHVMKNVDTNQDRLVTLEEFLAST....QRKE.
CferNb1  LEALFTKELEKVYDPKN..EEDDMRE..MEEERLRMREHVMKNVDTNQDRLVTLEEFLAST....QRKE.
HarmNb1  LEALFTKELEKVYDPKN..EEDDMRE..MEEERLRMREHVMKNVDTNQDRLVTLEEFLAST....QRKE.
RsinNb1  LEALFTKELEKVYDPKN..EEDDMRE..MEEERLRMREHVMKNVDTNQDRLVTLEEFLAST....QRKE.
JjacNb1  LEALFTKELEKVYDPKN..EEDDMRE..MEEERLRMREHVMKNVDTNQDRLVTLEEFLAST....QRKE.
DordNb1  LEALFTKELEKVYDPKN..EEDDMRE..MEEERLRMREHVMKNVDTNQDRLVTLEEFLAST....QRKE.
NgalNb1  LEALFTKELEKVYDPKN..EEDDMRE..MEEERLRMREHVMKNVDTNQDRLVTLEEFLVST....QRKE.
RnorNb1  LEALFTKELEKVYDPKN..EEDDMRE..MEEERLRMREHVMKNVDTNQDRLVTLEEFLAST....QRKE.
MmusNb1  LEALFTKELEKVYDPKN..EEDDMRE..MEEERLRMREHVMKNVDTNQDRLVTLEEFLAST....QRKE.
MaurNb1  LEALFTKELEKVYDPKN..EEDDMRE..MEEERLRMREHVMKNVDTNQDRLVTLEEFLAST....QRKE.
CasiNb1  LEALFTKELEKVYDPKN..EEDDMRE..MEEERLRMREHVMKNVDTNQDRLVTLEEFLAST....QRKE.
NparNb   LEALFTKELEKVYDPKN..EEDDMVE..MEEERLRMREHVMKNVDANHDRLVTLEEFLKST....ENKD.
XlaeNb1  LEALFTKELEKVYDPKN..EEDDMVE..MEEERLRMREHVMKNVDTNQDRLVTLDEFLKST....ERKE.
XtroNb1  LEALFTKELEKVYDPKN..EEDDMVE..MEEERLRMREHVMKNVDANHDRLVTLDEFLKST....ERKE.
PmucNb1  LEALFTKELDKVYDPRN..EEDDMLE..MEEERLRMREHVMKNVDLNKDRLVTLEEFVKST....QRKE.
PbivNb1  LEALFTKELEKVYDPRN..EEDDMLE..MEEERLRMREHVMKNVDLNKDRLVTLDEFLKST....QRKE.
GjapNb1  LEALFTKELEKVYDPRN..EEDDMLE..MEEERLRMREHVMKNVDLNRDRLVNLDEFLKST....QKKE.
DrerNb1  LEALFTKELEKVYDPKN..EEDDMVE..MEEERLRMREHVMQNVDVNHDRLVSLEEFLKST....EKRE.
TrubNb1  LEALFTKELEKVYDPKN..EEDDMME..MEEERLRMREYFMKNVDVNQDRLVTLEEFLKST....EKKD.
TnigNb1  LEALFTKELEKVYDPKN..EEDDMME..MEEERLRMREYFMKNVDANKDRVVSLEEFLKST....EKKD.
IpunNb   LEALFTKELEKIYDPAN..EEDDMVE..MEEERLRMREHVMNEVDMDKDRLVSLGEFLIAT....KKKE.
DrerNb2  LESLFTKELEKIYDPTN..EEDDMVE..MEEERLRMREHVMNEVDSNKDRLVSLDEFLVAT....KKKE.
TrubNb2  LEALFTKELEKIYDPNN..EEDDMIE..MEEERLRMREHVMNEVDSNKDRLVSLEEFLVAT....RKKE.
HsapNb2  LEALFTKELEKVYDPKN..EEDDMVE..MEEERLRMREHVMNEVDTNKDRLVTLEEFLKAT....EKKE.
PpanNb2  LEALFTKELEKVYDPRN..EEDDMVE..MEEERLRMREHVMNEVDTNKDRLVTLEEFLKAT....EKKE.
GgorNb2  LEALFTKELEKVYDPKN..EEDDMVE..MEEERLRMREHVMNEVDTNKDRLVTLEEFLKAT....EKKE.
CatyNb2  LEALFTKELEKVYDPKN..EEDDMVE..MEEERLRMREHVMNEVDTNKDRLVTLEEFLKAT....EKKE.
CangNb2  LEALFTKELEKVYDPKN..EEDDMVE..MEEERLRMREHVMNEVDTNKDRLVTLEEFLKAT....EKKE.
RroxNb2  LEALFTKELEKVYDPKN..EEDDMVE..MEEERLRMREHVMNEVDTNKDRLVTLEEFLKAT....EKKE.
PabeNb2  LEALFTKELEKVYDPKN..EEDDMVE..MEEERLRMREHVMNEVDTNKDRLVTLEEFLKAT....EKKE.
MmulNb2  LEALFTKELEKVYDPKN..EEDDMVE..MEEERLRMREHVMNEVDTNKDRLVTLEEFLKAT....EKKE.
MfasNb2  LEALFTKELEKVYDPKN..EEDDMVE..MEEERLRMREHVMNEVDTNKDRLVTLEEFLKAT....EKKE.
CcapNb2  LEALFTKELEKVYDPKN..EEDDMVE..MEEERLRMREHVMNEVDTNKDRLVTLEEFLKAT....EKKE.
PtroNb2  LEALFTKELEKVYDPKN..EEDDMVE..MEEERLRMREHVMN............................
OgarNb2  LEALFTKELEKVYDPKN..EEDDMVE..MEEERLRMREHVMNEVDINKDRLVTLEEFLKAT....EKKE.
PcoqNb2  LEALFTKELEKVYDPKN..EEDDMVE..MEEERLRMREHVMNEVDTNKDRLVTLEEFLKAT....EKKE.
TchiNb2  LEALFTKELEKVYDAKN..EEDDMVE..MEEERLRMREHVMNEVDTNKDRLVTLEEFMRAT....ERKE.
MjavNb2  LEALFTKELEKVYDPKN..EEDDMVE..MEEERLRMREHVMNEVDTNKDRLVTLEEFLKAT....EKKE.
HarmNb2  LEALFTKELEKIYDPKN..EEDDMVE..MEEERLRMREHVMNEVDANKDRLVTLEEFLKAT....EKKE.
RsinNb2  LEALFTKELEKVYDPKN..EEDDMVE..MEEERLRMREHVMNEVDANKDRLVTLEEFLKAT....EKKE.
DordNb2  LEALFTKELEKVYDPRN..EEDDMVE..MEEERLRMREHVMNEVDTNKDRLVTMEEFLKAT....EKKE.
BmutNb2  LEALFTKELEKVYDPKN..EEDDMVE..MEEERLRMREHVMNEVDTNKDRLVTLDEFLKAT....EKKE.
VpacNb2  LEALFTKELEKVYDPKN..EEDDMVE..MEEERLRMREHVMNEVDTNKDRLVTLEEFLKAT....EKKE.
CferNb2  LEALFTKELEKVYDPKN..EEDDMVE..MEEERLRMREHVMNEVDTNKDRLVTLEEFLKAT....EKKE.
CbacNb2  LEALFTKELEKVYDPKN..EEDDMVE..MEEERLRMREHVMNEVDTNKDRLVTLEEFLKAT....EKKE.
JjacNb2  LEALFTKELEKVYDPKN..EEDDMVE..MEEERLRMREHVMNEVDSNKDRLVTLEEFLKAT....EKKE.
CasiNb2  LEALFTKELEKVYDPKN..EEDDMIE..MEEERLRMREHVMTEIDANKDRLVTLEEFLKAT....EKKE.
NgalNb2  LEALFTKELEKVYDPRN..AEDDMME..MEEERLRMREHVMNEVDSNKDRLVTLEEFLKAT....EKKE.
RnorNb2  LEALFTKELDKVYNPQN..AEDDMIE..MEEERLRMREHVMNEIDNNKDRLVTLEEFLRAT....EKKE.
MmusNb2  LEALFTRELEKVYNPQN..AEDDMIE..MEEERLRMREHVMSEIDNNKDRLVTLEEFLRAT....EKKE.
MaurNb2  LEALFTKELEKVYDPRN..AEDDMIE..MEEERLRMREHVMNEIDSNKDRLVTLEEFLRAT....EKKE.
PbivNb2  LEALFTKELEKVYDPKN..EEDDMVE..MEEERLRMREHVMNEVDINRDRLVTLEEFMRAT....EKKE.
PmucNb2  LEALFTRELEKVYDPRN..EEDDMVE..MEEERLRMREHVMNEVDINKDRLVTLEEFMRAT....EKRE.
GjapNb2  LEALFTKELEKVYDPKN..EEDDMVE..MEEERLRMREHVMSEVDINKDRLVTLDEFLRAT....EKRE.
LchaNb   LEALFTKELAKVYDPKN..EEDDMAE..MEEERLRMREHVMNEVDINKDRLITMEEFLKAT....EKKE.
XlaeNb2  LEALFTKELEKVYDPNN..EEDDMVE..MEEERLRMREHVMMEVDLNKDRLIDLQEFIRAT....EKRE.
XtroNb2  LEALFTKELEKVYDPNN..EEDDMVE..MEEERLRMREHVMTEVDLNRDRLIDLQEFIRAT....EKRE.
AcalNb   VDAVLQIELDKVYDARNSPDEDDPME..RQEEMNRMREHVFQEMDKDHNQRISFEEFIEYTGAMGDNEE.
BglaNb   VDAVLQLELDKVYDAKNSPDEDDPVE..RQEEMNRMREHVFQEMDKDKNWRISFQEFIDYTGS.......
LgigNb   VEALLQHELDKAYDEKNA.DEDDPVE..RVEEMNRMREHIFTEIDQNHDKFITRDEFLIYTGPKGDRKE.
DmagNb   VKALFIKELDKAYDPNA..PEDDMAE..RYEEMERMREHVFNETDTNRDYLISFPEFLEQT....RKQE.
DpulNb   VKALFSKELDKAYDPNA..PEDDMAE..RYEEMERMREHVFNETDTNRDYLISFPEFLEQT....RKQE.
HrobNb   LEALFQKELDKVYNENA..TEDDMME..RYEEMNRMREHVMKEIDANRDSVISLEEFVNST....QREE.
HaztNb   VRALFLKELEKAYDPNA..SEDDLHE..RDEEMERMREHIFAEADANDDDLISYDEFLEQT....RRNE.
TurtNb   VEALLTLEIKKMYDPNN..PEDDPNE..MEEEYHRMREHVYKEADKNKDGLISQKEFIELT....QKTE.
PtepNb   VEAVLIPEVRKVYNPNN..EEDDPVE..MMEDMYRMREHIFNQSDTNKDHLISLAEFLKMT....EDDS.
SmimNb   VEAILLPEVKKLYNPNN..EEDDPAE..MEEEFHRMREHIFNETDAHLIGSTERYRYILII........
GoccNb   IEAILQLEAKKMYMSDPQ.HPGDPRE..MEEELNRMREHVFKEVDKDKDGLISKKEFLEMT....EVDS.
DsuzNb   VKALFVKELDKVYQSGL..PEDDMRE..RAEEMERMREHYFQETDLNHDGLVSIEEFMVQT....NKEE.
DmelNb   VKALFVKELDKVYQSDL..PEDDMRE..RAEEMERMREHYFQETDMNHDGLISIDEFMVQT....NKEE.
AaegNb   VKVLFINELNKMYEAGH..PEDDMKE..RAEEMERMREHVFQEADTNKDGLISYEEFIEQT....KRDE.
CtelNb   IEALFQKELDKVYDPNA..EEDDMME..RFEEMNRMREHVMKEIDTDGDKLVSMEEFVAST....NRDE.
OvicNb   LEALFIKEVEKIYQD....KEADPRE..KFEEISRMREHVLREIDLDGDKMVSREEFLKAA....DQTQ.
SkowNb   LEALFVNEIEKIYG.....DDSDPRE..KMEEMSRMREHVMKEIDTDKDFMVSMREFMDAT....RDKD.
SpurNb   LEALFIKEVEKVYKDQ...KTADVQE..KFEELSRMREHVVNQIDNNKDKMVSLAEFMEAA....SASD.
NvecNb   LEALFVKEVTKLYNPKD..EDYDPKE..RDEEISRMREHVMNEIDKDKDGFVSQDEFLQSS....KGEE.
EpalNb   LEALFVKEVGRLHDPKS..EDFDKRE..RDEELARMREHVMNEIDKDKDGFVSKDEFLAAS....KSDD.
OfavNb   LEALFVKEVSV...........................HVIYKPD.........................
HvulNb   LETIFLADIDKVYNESN..PEVDLRE..RSEEIERMREHVMKNMDKDKDGLISFTEFMDET....KSED.
CbriNb   LESLFQLELEKMYNETN..PDDDMKE..RAEEMYRMREHVMKQIDVNQDRMISMQEFLNDA....DSQNA
CbreNb   LESLFQLELEKMYNETN..PDDDMKE..RAEEMYRMREHVMKQIDVNQDRMISMQEFLNDA....DAQNA
HconNb   LEALFEKEIEKVYNATD..PDDDPRE..RLEEMHRMREHVTKQMDKNGDRLISMEEFLQDT....EAQT.
```

```
TcanNb   LEALFQLELQKVYNESD..PDDDPRE..RIEEMYRMREHVVQQMDKNNDRMISLAEFLADA....EAQT.
SratNb   LDALFQLELGKMYNESN..PDDDPKE..KMEEMHRMREHVLNQMDTNKDKMISMEEFLADS....EAQAS
ShaeNb   VEALLQRELEKVYNPED..PDYDPWE..EKYDQSRMRQKFMERFDTDGDYFVSRDEFLRGV....NQPY.
SmanNb   VEAFLQTELEKVYNPED..PDYDPWE..ERHDQKKMRQKFMDRFDTDGNYLVSRDEFLRGV....NQPY.
SjapNb   IEAFLQREIEKVYNPED..PDFDPWE..EKYDQIKMRQKFMERFDTDKDYFISREEFLKNV....KVPY.
AqueNb   VEALFVNEIRKYYGKD...FEND.RE..AHEDMSRMREHVMSEVDRDHDSLITLEEFVRYA....NTDL.
SrosNb   LEAIFHRSAVKLHTID...GETDTDA..VEEEMLDMRAWVMKEVDTDKDQLVSRDEFITFA....NSPE.
BdenNb   LRLAFQ.....GYEQDT..GKEKAH..IDLADLETMIDHALAEDDTNNDGMISWEEYLESQ.........
BsalNb   LRLAFQ.....GFERDT..GKESVPK.IDMADLETMIDHALIEDDTDNDGMISWEEYLESQ.........
CcucNb   LRAAFTD.....FGD....EHEDVTHYFSLEEVTDMVDHVLEEDDLNGDGLISWSEYLESQ.........
ArepNb   LRAAFSD.....FDHD...SEEDPTKHVSLEDITLMVDHVLGEDDLDGDGMISWEEYMQSQ.........
McirNb   LRAAFTD.....FDED...EQHGEDQDITLQEVTDMVDHVLMEDDKNGDGLISWSEYLESQ.........
MbreNb   VEALVNAEATRLHTIEDF...VDEQA..VAYEAVRMRQAFMADVDTNKDDMISLQEFMLFA....ASNS.
DqueNb   IEAIYG..VHHVYSQKK..SKDE.IE..HQQKADHIVNTVLEKIDKNKDGKISLEEFLEVG....VDGLP
FpinNb   IEAIYG..VHHAYSQKK..SKDE.IE..HQKKADNIVNTVLEKLDKNKDGKISLDEFLAVG....LDGLP
FradNb   IEAIYG..VHHVYSQKK..SKDD.IE..HQKKADHIVNTVLGRLDKDKDGKVTLEEFLSVG....LDGLP
GfroNb   IEAIYG..VHHVYSQKK..SKDE.VE..HQKKADHIVNTVLKLLDKDKDGKISMEEFAAVG....LDGLP
ShirNb   VEAIYG..VHHVYSQKK..SKDD.EE..HKAKAKVIVDTIMNALDKNKDEVVTMEEFESVG....LAALP
PostNb   IEAVYG..VHHVYSQKK..SKDD.IE..HQEKAEQIVKTVLARIDANNDGVITAEELVAVG....LDGLP
RsolNb   IEAIYG..VHHIYSKRK..TPSE.EA..QAAKAKQVADAVLAAMDTNGDGFITMEEFLAKG....LDALP
NirrNb   IRALYG.............TDHDANI..PEHISQSIEKTVLDAIDRNRDGVISLNEWLEYK....KNGGS
ScerNb   ILSLYGLNREEIVGAGDGMGQHDESEKIDNEMAKRVVSLIMRLLDVDDNTKITKEEYLQFA....KRGNK
VpolNb   ILGLYGLNRGEVVGNGDGMGDHDNSESIDEELAKRVVKFIKKLIDVNDDSKITKEEYLSFA....KRGNK
TphaNb   ILALYGLNRGEVVGKGDGMGEHDSSELIDEENSKRVVTFIKKLLDVDDDSRIAKEEYLAFA....KKGHK
CglaNb   ILGLYGLNRDEVIGAGDGMGEHDNSEKIDETTAKNVVKFIMKLLDVNDDTRILRDEYLEFA....RRGKK
LmirNb   VLTMYGLDREEVVGAGDGMGSHDNSEMIDETLADRVVQLIMALLDVDDDTKIEKEEYMAFA....KRGKK
EgosNb   ILSLYGLSRPEVVGTGDGMGQHDDSERIDEETANRIVKLIGELLDTNDDGKVTKAEYLNFA....KRGKS
LngbNb   WAELFQ.....VYNVHPA.............YAPLAFEQLDMNGDGFLSKDEILFLI....DDFF.
McolNb   WAGLLS.....VYNVSPV.............YAPLVFPKIDTNQDGFVSKEEALQLI....RDFF.
CchtNb   WAGILS.....VYNVSPV.............YAPRVFPKLDTNQDGFLSKEEVLQLI....REFF.
SmajNb   WGNLLC.....IYNTSPI.............YAPLIFSRLTTDTDGNLSREAVLSHI....DEFF.
NnodNb   WGQLLA.....AFNESPV.............YAPLVFPPLDADQDGYLTKAEVLQHF....SSFC.
MproNb   YKQGFL.....SHFSDGN.............LTDEIFSKLDLNGDGHLSKEEVLQHV....HDFF.
PsojNb   LVQFMG.....SEEQA...............QEVINDVDANGDGVISFEEFVAMM....NRKG.
PhalNb   LVQFMG.....SEEQA...............QEIIDDVDANGDGVISFEEFVAMM....NRKG.
PinsNb   LVQFMG.....SEEQA...............DEVMRDVDTNGDGQISFDEFVAMM....NRQG.
AcanNb   LIQFMG.....SEEQA...............QHVIDDVDVDGDGQISFEEFVAMM....QQKN.
TclaNb   LVRFMG.....SEDHA...............QEVMNEIDSNGDGQISFEEFVAMM....QKRG.
EsilNb   LVSIFG.....SEDHA...............REIVGDIDLDGDGEISFDEFRQMM....EEDQ.
EguiNb   LQQACK.....EHNMTDV.............LIEDIIREVDQDNDGRIDYDEFVAMM....QKGN.
DdisNb   IEESIKKLNIPLYSEQEL.................IRLFHRIDKNRDNQIDFNEWRELLVLL.PNSN.
AsubNb   IEQSIDRIGLKLYSDQEL.................VRLFQRVDTNNDNKIDFNEWRELLVLL.PNSN.
AcasNb   VRGSLRRLGMK.YDDGAV.................TKLIKRIDVDGNGKIDFNEWQTFLLLV.PSAT.
LocuNb   IKQALEDLGMD.ISAEEA.................KKILQSIDADGTMTVDWNEWREHFLLN.PATN.
CcarNb   IQQSFKDLGLN.LTDRDA.................EKILHSIDADGTMTVDWNEWREHFLFN.PADD.
SansNb   IQQSFKDLGLN.LTDRDA.................EKILHSIDADGTMTVDWNEWREHFLFN.PADN.
GproNb   LHELLDTMHGLFTSPGD..............ETLRRMRDAATCRAKPNKRD..AEEEELSRA.........
RgloNb   LKAVMDSLLEMFEKSGKA............................AGVVEEEEVYMKA.........
AbacNb   .............................SFEHKQAATETVTEQMEKD.......................
TgonNb   .............................EYVSKRAETEASLEELQQD.......................
HhamNb   .............................EYVSKRAETEESLEELQQD.......................
BbesNb   .............................EYVSKRAESEPSSEELQPD.......................
CsuiNb   .............................EYISKRAPREEAAEELQPT.......................
CcayNb   .............................EHLSKRADTQTVAAELQPN.......................
TbruNb   KAALSGSDVSG.........SDEAPFEGVGVENADRLSKEVSLEKHSGTGQQVAATRT.KQALSALGKQ..
LbraNb   VAALTIHEDTAVEESRP.QEDGGALERGRMDDARRLRQQVQQQLHV...QHVTEEEM.ERAVRALEEQFN
LseyNb   VSALSYTEQQLVYDNRSPESDGGEVQCSVMLHATRLQQEVAREANV...TQVMQEQVKDEAMEDCKKRFG
PfalNb   LEILFNNNHVKLYNNKDDNGKNEQNEQGKNEENEKSKKHIISKVDMNNEENFTMEEWKMLM....KKNEQ
```

```
                330                                   340
HsapNb1    ..FGD.TGEGWE.........................TV.E..MHPAY..TEEE.L..............
PtroNb1    ..FGD.TGEGWE.........................TV.E..MHPAY..TEEE.L..............
PpanNb1    ..FGD.TGEGWE.........................TV.E..MHPAY..TEEE.L..............
PabeNb1    ..FGD.TGEGWE.........................TV.E..MHPAY..TEEE.L..............
GgorNb1    ..FGD.TGEGWE.........................TV.E..MHPAY..TEEE.L..............
CatyNb1    ..FGD.TGEGWE.........................TV.E..MHPAY..TEEE.L..............
MmulNb1    ..FGD.TGEGWE.........................TV.E..MHPAY..TEEE.L..............
RroxNb1    ..FGD.TGEGWE.........................TV.E..MHPAY..TEEE.L..............
CangNb1    ..FGD.TGEGWE.........................TV.E..MHPAY..TEEE.L..............
MfasNb1    ..FGD.TGEGWE.........................TV.E..MHPAY..TEEE.L..............
MnemNb1    ..FGD.TGEGWE.........................TV.E..MHPAY..TEEE.L..............
CcapNb1    ..FGD.TGEGWE.........................TV.E..MHPAY..TEEE.L..............
OgarNb1    ..FGD.TGEGWE.........................TV.E..MHPAY..TEEE.L..............
PcoqNb1    ..FGD.AGEGWE.........................TV.E..MHPAY..TEEE.L..............
GvarNb1    ..FGD.TGEGWE.........................TV.E..MHPAY..TEEE.L..............
TchiNb1    ..FGD.TGEGWE.........................TV.E..MHPAY..TEDE.L..............
BmutNb1    ..FGD.TGEGWEQ...GKGGVPLPTSPVLT.....LQTV.E..MHPAY..TEEE.L..............
MjavNb1    ..FGD.TGEGWE.........................TV.E..MHPAY..TQEE.L..............
VpacNb1    ..FGD.TGEGWE.........................TV.E..MHPAY..TEDE.L..............
CbacNb1    ..FGD.TGEGWE.........................TV.E..MHPAY..TEDE.L..............
VpacNb1    ..FGD.TGEGWE.........................TV.E..MHPAY..TEDE.L..............
CferNb1    ..FGD.TGEGWE.........................TV.E..MHPAY..TEDE.L..............
HarmNb1    ..FGD.TGEGWE.........................TV.E..MHPAY..TEDE.L..............
RsinNb1    ..FGD.TGEGWE.........................TV.E..MHPAY..TEDE.L..............
JjacNb1    ..FGD.TGEGWE.........................TV.E..MHPAY..TEEE.L..............
DordNb1    ..FGD.TGEGWE.........................TV.E..MHPAY..TEEE.L..............
NgalNb1    ..FGD.TGEGWE.........................TV.E..MHPAY..TEEE.L..............
RnorNb1    ..FGE.TAEGWK.........................TV.E..MYPAY..TEEE.L..............
MmusNb1    ..FGD.TGEGWK.........................TV.E..MSPAY..TEEE.L..............
MaurNb1    ..FGD.TGEGWK.........................TV.E..MYPAY..TEEE.L..............
CasiNb1    ..FGD.TGEGWERALCSEAGPWLGLTPVMASFRSPPQTV.E..MQPAY..TEEE.L..............
NparNb     ..FNN.P.DGWQ.........................TV.E..ETQIY..SEEE.L..............
XlaeNb1    ..FNE.A.DGWE.........................TV.D..ETQIY..TEEE.L..............
XtroNb1    ..FNE.A.DGWE.........................TV.D..ETQVY..TEEE.L..............
PmucNb1    ..FNE.A.DGWE.........................TV.E..ETQIY..SEAE.L..............
PbivNb1    ..FNE.A.DGWE.........................VC.E..RTL........................
GjapNb1    ..FNE.A.DGWK.........................TV.E..ETQVY..SEAE.L..............
DrerNb1    ..LNN.Q.KEWE.........................TL.D.DTKPVY..TEEE.L..............
TrubNb1    ..FNN.P.KEWE.........................TL.D..MKEVY..TEEE.L..............
TnigNb1    ..FNN.P.TEWE.........................TL.D..TKQVY..TEEE.L..............
IpunNb     ..FLE.P.DSWE.........................TL.E..QNQAY..TEEE.M..............
DrerNb2    ..FLE.P.DSWE.........................TL.E..QNQAY..TEEE.M..............
TrubNb2    ..FLE.P.DSWE.........................TL.E..QNQVY..TDEE.M..............
HsapNb2    ..FLE.P.DSWE.........................TL.D..QQQFF..TEEE.L..............
PpanNb2    ..FLE.P.DSWE.........................TL.D..QQQFF..TEEE.L..............
GgorNb2    ..FLE.P.DSWE.........................TL.D..QQQFF..TEEE.L..............
CatyNb2    ..FLE.P.DSWE.........................TL.D..QQQFF..TEEE.L..............
CangNb2    ..FLE.P.DSWE.........................TL.D..QQQFF..TEEE.L..............
RroxNb2    ..FLE.P.DSWE.........................TL.D..QQQFF..TEEE.L..............
PabeNb2    ..FLE.P.DSWE.........................TL.D..QQQFF..TEEE.L..............
MmulNb2    ..FLE.P.DSWE.........................TL.D..QQQFF..TEEE.L..............
MfasNb2    ..FLE.P.DSWE.........................TL.D..QQQFF..TEEE.L..............
CcapNb2    ..FLE.P.DSWE.........................TL.D..QQQFF..TEEE.L..............
PtroNb2    ...........E.........................TL.D..QQQFF..TEEE.L..............
OgarNb2    ..FLE.P.DSWE.........................TL.D..RQQFF..TEEE.L..............
PcoqNb2    ..FLE.P.DSWE.........................TL.D..QQQFF..TEEE.L..............
TchiNb2    ..FLE.P.DSWE.........................TL.D..QQQLF..TEEE.L..............
MjavNb2    ..FLE.P.DSWE.........................TL.D..QQQFF..TEEE.L..............
HarmNb2    ..FLE.P.DSWE.........................TL.D..QQQFF..TEEE.L..............
RsinNb2    ..FLE.P.DSWE.........................TL.D..QQQFF..TDEE.L..............
DordNb2    ..FLE.P.DSWE.........................TL.D..QQQLF..TEEE.L..............
BmutNb2    ..FLE.P.DSWE.........................TL.D..HQQFF..TEEE.L..............
VpacNb2    ..FLE.P.DSWE.........................TL.D..QQQFF..TEEE.L..............
CferNb2    ..FLE.P.DSWE.........................TL.D..QQQFF..TEEE.L..............
CbacNb2    ..FLE.P.DSWE.........................TL.D..QQQFF..TEEE.L..............
JjacNb2    ..FLE.P.DSWE.........................TL.D..QQQLF..TEEE.L..............
CasiNb2    ..FLE.P.DSWE.........................TL.D..QQQLF..TEEE.L..............
NgalNb2    ..FLE.P.DSWE.........................TL.D..QQQLF..TEEE.L..............
RnorNb2    ..FLE.P.DSWE.........................TL.D..QQQLF..TEEE.L..............
MmusNb2    ..FLE.P.DSWE.........................TL.D..QQQLF..TEDE.L..............
MaurNb2    ..FLE.P.DSWE.........................TL.S..QQQLF..TEEE.L..............
PbivNb2    ..FLE.P.ESWE.........................TL.D..QQQIF..TEED.L..............
PmucNb2    ..FLE.P.ESWE.........................TL.D..QQQVF..TEED.L..............
GjapNb2    ..FLE.P.DSWE.........................TL.D..QQQLF..TEEE.L..............
LchaNb     ..FLE.P.DSWE.........................TL.D..KQQLF..TEEE.L..............
XlaeNb2    ..FLE.P.DGWE.........................TV.A..NQQLY..TEEE.L..............
XtroNb2    ..FLE.P.DGWE.........................TV.A..DQPIY..TEEE.L..............
AcalNb     ..FKK.D.EGWK.........................TV.D..EEPVF..TDED.Y..............
BglaNb     ......................................................................
LgigNb     ..FEE.N.EDWK.........................V.......SCY..S...................
DmagNb     ..FER.D.AGWN.........................TI.D..EQPVY..SQQE.Y..............
DpulNb     ..FER.D.PGWN.........................TI.D..EQPVY..SQQE.Y..............
HrobNb     ..FKK.D.EEWD.........................TL.E..DERQF..TDEE.L..............
HaztNb     ..FER.D.EGWQ.........................TL.D..EQELY..SQSQ.Y..............
TurtNb     ..FER.D.EGWK.........................GI.D..EQQMF..NEDE.L..............
PtepNb     ..FEQ.D.EGWQ.........................GL.D..EQQVY..SESE.L..............
SmimNb     .....................................GI.D......Y......VM..............
GoccNb     ..FDR.D.EGWK.........................GL.D..DEQVY..TQEE.L..............
DsuzNb     ..FQK.D.PEWE.........................TI.D..RQPQY..SHEE.Y..............
DmelNb     ..FQK.D.PEWE.........................TI.D..RQQQY..THEE.Y..............
AaegNb     ..FQK.D.PEWD.........................TV.D..HQPQF..THEE.Y..............
CtelNb     ..FKE.N.EEWE.........................TI.D..EDEEF..TDEE.L..............
OvicNb     ..FEN.D.DGWK.........................DI.N..QEEQF..TEDE.L..............
SkowNb     ..FDN.E.ESWE.........................DL.G..DKDLF..TEEE.L..............
SpurNb     ..FEE.D.SEWQ.........................DL.N..QESQF..TEGD.L..............
NvecNb     ..FDK.D.DGWK.........................SV.E..DERPY..TDEE.L..............
EpalNb     ..FEK.D.EGWK.........................SV.EQPDQRPY..TDDE.L..............
OfavNb     ..............................................AY..T....L..............
HvulNb     ..FEK.D.EDWK.........................PL.T..EQDQF..TEEE.L..............
CbriNb     NPPKQ.E.EAWE.........................DL.G..QKKVY..TDEE.L..............
CbreNb     NPPKKDD.EGWQ.........................DL.G..EQKFY..TDEE.L..............
HconNb     ..PNK.D.PGWT.........................DL.G..DQQVY..TDEE.L..............
```

```
TcanNb   ..PNK.D.DGWK.........................DI.G..DEAVY..SDDE.L..............
SratNb   ..TPS.P.EGWK.........................AL.D..ETKIY..SEEE.L..............
ShaeNb   ..PSY.D.SDWM.........................TAQD..EEDMFETNEEE.L..............
SmanNb   ..PSY.D.RDWM.........................TAQN..EEDMFKTNEEE.L..............
SjapNb   ..VLG.D.HYWK.........................TAQD..VEDVFETDESE.L..............
AqueNb   ..FNI.K.EEWK.........................PVGEDDEHKEF..TDDE.F..............
SrosNb   ..FQT.N.RDWK.........................AV.....FPDF..NGTE.L..............
BdenNb   ......................................................................
BsalNb   ......................................................................
CcucNb   ......................................................................
ArepNb   ......................................................................
McirNb   ......................................................................
MbreNb   ..FAD.S.KNWT.........................AI.....VPEF..DDAS.L..............
DqueNb   N.FDSLGAEGHHY........................DM....ESEFF.......LHHE...........
FpinNb   N.FDDLGAEGHHY........................DM....ESEFF.......LHHE...........
FradNb   N.FDDLGAEGHHY........................DV....ESEFF.......LHHE...........
GfroNb   N.FDDLGAEGHHY........................DV....ESEFF.......LHHEGSPLRYALLLS
ShirNb   N.FDNMGAEGHHY........................DV....ESEFF.......LHHE...........
PostNb   N.FDNLGAEGHHY........................DV....ESEFF.......LHHE...........
RsolNb   D.FSALGAEGHHY........................DV....ESEFF.......LHHE...........
NirrNb   ..LPDPG.....................................ITY.N..FGHHF..DEET.........D......
ScerNb   ..FPDLG.....V........................GV.G..HHSDFE.LEYE.IHHW...........
VpolNb   ..LPDLG.....V........................GV.G..HHADFE.LEYE.LHHW...........
TphaNb   ..FPDLG.....V........................GV.G..HHADFE.LEYE.LHHW...........
CglaNb   ..LPDLG.....V........................GV.G..HHSDFE.REYE.IHHW...........
LmirNb   ..FPDLG.....V........................GV.G..HHSDFE.LEYE.IHHW...........
EgosNb   ..FPDLG.....V........................GV.G..HHLDFE.KEFN.VHHW...........
LngbNb   ..CGD.D.....P...........................TSGANSMF......................
McolNb   ..YSN.D.....P...........................EAPANSMF......................
CchtNb   ..YSD.N.....P...........................DVPANEMF......................
SmajNb   ..YSD.D.....P...........................GAPGNGMF......................
NnodNb   ..CSD.D.....A...........................DNPANGMF......................
MproNb   ..YSD.D.....P...........................EAPGNWIL......................
PsojNb   ..YGD.D.....S...........................EMKVQSP.......................
PhalNb   ..YGD.D.....S...........................EMDAYS........................
PinsNb   ..LND.S.....N...........................SAGAAPPL......................
AcanNb   ..YSD.D.....DI..........................SMPVCD........................
TclaNb   ..LGE.E.....L...........................TPPANT........................
EsilNb   ..TTT.T.....A...........................TAPVDSAA......................
EguiNb   ..MGI.G.....PI..........................GR............................
DdisNb   ..LQLII.SFWKDSQILDAGFDGGFIPPMVE..KP.ISLISA.FKACYKVSPESAVKFGTY.........
AsubNb   ..LSAAL.AYWKDAQILDGGDDGGFAPPPATLAAP.RSIAET.FRVVYAVSPESAVKFATF.........
AcasNb   ..VDAIF.RYWEDAMMAFDSEDGVILPPAHK..KP..SIDAI.SRQIYAVAPEKAIKFWTY.........
LocuNb   ..IQEII.RYWKHSSVLDIGD.SLTIPDEFT..KTKIGLVSG.FKQMLKIAPETAIKFMAY.........
CcarNb   ..LQEII.RYWKHSTVLDIGD.SLTIPDEFT..KSKISLVDG.FKQMIKIAPETAIKFMAY.........
SansNb   ..LQEII.RYWKHSTVLDIGD.SLTIPDEFT..KSKISLVDG.FKQMIKIAPETAIKFMAY.........
GproNb   ............V........................GV........F..LSTA.LKGSTGGSGTA....
RgloNb   ..VSS............................................F..LNSA.LK.............
AbacNb   ........GSWR...............................IAGYF.......I..............
TgonNb   ........AVWRCPVC..............RFRTP..SV.QSSLKKLF.......V..............
HhamNb   ........AVWRCPVC..............RFRIP..SV.QPSLKKLF.......V..............
BbesNb   ........AVWRCPVC..............RFKRP..GV...SLKKLF.......V..............
CsuiNb   ........ATWRCPVCVGGAKKSTASPCALRLRRK.ATA.ERSLKKIF.......V..............
CcayNb   ........AHWSCPLC..............KAKQPCATA.PGALKRLF.......V..............
TbruNb   .NLTD...EQWMQ.........RLYDAIMKSTS....HV.DRQKQE...GQERQ...RER.....ER...
LbraNb   A..TD...EEWVA.........NLRLAALAMVR....TL.QPAREPWL.SPEES........AGFAH...
LseyNb   T..TD...EEWLV.........ELHSVAAAMIK....SV.HQSRQP...APQHQ...HQHYLSPAAR...
PfalNb   TSFKDLNTDGWKWLI.......LSEEVQNKIFL....TVAELQDKKIYHKFEPE.LHHAYGKIGCILVEC
```

```
               350        360
                 .          .
HsapNb1  .....RRFEEELAA.REAE..............LNAKAQR..............................
PtroNb1  .....RRFEEELAA.REAE..............LNAKAQR..............................
PpanNb1  .....RRFEEELAA.REAE..............LNAKAQR..............................
PabeNb1  .....RRFEEELAA.REAE..............LNAKAQR..............................
GgorNb1  .....RRFEEELAA.REAE..............LNAKAQR..............................
CatyNb1  .....RRFEEELAA.REAE..............LNAKAQR..............................
MmulNb1  .....RRFEEELAA.REAE..............LNAKAQR..............................
RroxNb1  .....RRFEEELAA.REAE..............LNAKAQR..............................
CangNb1  .....RRFEEELAA.REAE..............LNAKAQR..............................
MfasNb1  .....RRFEEELAA.REAE..............LNAKAQR..............................
MnemNb1  .....RRFEEELAA.REAE..............LNAKAQR..............................
CcapNb1  .....RRFEEELAA.REAE..............LNAKAQR..............................
OgarNb1  .....RRFEEELAI.REAE..............LNAKAQR..............................
PcoqNb1  .....RRFEEELAA.REAE..............LNAKAQR..............................
GvarNb1  .....RHFEEELAA.REAE..............LNAKAQR..............................
TchiNb1  .....RRFEEELAA.REAE..............LNAKAQR..............................
BmutNb1  .....RRFEEELAA.REAE..............LNAKAQR..............................
MjavNb1  .....KRFEEELAA.REAE..............LNAKAQR..............................
VpacNb1  .....RRFEEELAA.REAE..............LNAKAQR..............................
CbacNb1  .....RRFEEELAA.REAE..............LNAKAQR..............................
VpacNb1  .....RRFEEELAA.REAE..............LNAKAQR..............................
CferNb1  .....RRFEEELAA.REAE..............LNAKAQR..............................
HarmNb1  .....RRFEEELAA.REAE..............LNAKAQR..............................
RsinNb1  .....RRFEEELAA.REAE..............LNAKAQR..............................
JjacNb1  .....RRFEEELAA.REAE..............LNAKAQR..............................
DordNb1  .....RRFEEELAA.REAE..............LNAKAQR..............................
NgalNb1  .....RRFEEELAV.REAE..............LNAKAQR..............................
RnorNb1  .....KRFEEELAA.REAE..............LNARAQR..............................
MmusNb1  .....KRFEEELAA.REAE..............LNARAQR..............................
MaurNb1  .....RRFEEELAA.REAE..............LNAKAQR..............................
CasiNb1  .....RRFEEELAV.REAE..............LNAKAQR..............................
NparNb   .....RRFEQELVA.QESA..............LNQRAEE..............................
XlaeNb1  .....RRFEQELSA.QESA..............LNQRAEE..............................
XtroNb1  .....RKFEQELSA.QETA..............LNQRAEE..............................
PmucNb1  .....QRFEAELKA.QEEE..............LNRRAEQ..............................
PbivNb1  ......................................................................
GjapNb1  .....QRFEVELVA.QEAE..............LGRRAEQ..............................
DrerNb1  .....QRFETELRD.KELE..............LGRRAEK..............................
TrubNb1  .....QRFEAELRD.KEEE..............LRRKKEK..............................
TnigNb1  .....QRFEAELQD.KEEE..............LRRKKEK..............................
IpunNb   .....REFEEHLTR.QEQD..............LNQKTAE..............................
DrerNb2  .....REFEEQLVR.QEED..............LNQKAAD..............................
TrubNb2  .....REFEEHLTQ.QEQD..............LNMRSAD..............................
HsapNb2  .....KEYENIIAL.QENE..............LKKKADE..............................
PpanNb2  .....KEYENIIAL.QENE..............LKKKADE..............................
GgorNb2  .....KEYENIIAL.QENE..............LKKKADE..............................
CatyNb2  .....KEYENIIAL.QENE..............LKKKADE..............................
CangNb2  .....KEYENIIAL.QENE..............LKKKADE..............................
RroxNb2  .....KEYENIIAL.QENE..............LKKKADE..............................
PabeNb2  .....KEYENIIAL.QENE..............LKKKADE..............................
MmulNb2  .....KEYENIIAL.QENE..............LKKKADE..............................
MfasNb2  .....KEYENIIAL.QENE..............LKKKADE..............................
CcapNb2  .....KEYENIIAL.QENE..............LKKKADE..............................
PtroNb2  .....KEYENIIAL.QENE..............LKKKADE..............................
OgarNb2  .....KEYENIIAL.QESE..............LKKRAHE..............................
PcoqNb2  .....KEYENIIAL.QESE..............LKKRADE..............................
TchiNb2  .....KEYENIIAL.QENE..............LKKKADE..............................
MjavNb2  .....KEYENLISL.QENE..............LKKKAEE..............................
HarmNb2  .....KEYENLISL.QENE..............LKKKADE..............................
RsinNb2  .....KEYENLISL.QENE..............LKKKADE..............................
DordNb2  .....KEYENIIAL.QENE..............LKKKADD..............................
BmutNb2  .....KEYENLISV.QESK..............LKKKADE..............................
VpacNb2  .....KEYENLISV.QEIE..............LKKKADE..............................
CferNb2  .....KEYENLISV.QEIE..............LKKKADE..............................
CbacNb2  .....KEYENLISV.QEIE..............LKKKADE..............................
JjacNb2  .....KEYESMISL.QEDE..............LKRKADE..............................
CasiNb2  .....KEYENHISL.QEDE..............LKKKAEE..............................
NgalNb2  .....KEYENIIAL.QEDE..............LKKKADE..............................
RnorNb2  .....KEYESIIAI.QENE..............LKKKADE..............................
MmusNb2  .....KEYESIIAI.QENE..............LKKRAEE..............................
MaurNb2  .....KEYESIIAM.QENE..............LKKKADE..............................
PbivNb2  .....KEFESHISQ.QENE..............LQKQALE..............................
PmucNb2  .....KDFESHIFQ.KEDE..............LQKQALE..............................
GjapNb2  .....KEFESHIFQ.QEDE..............LQKKAKE..............................
LchaNb   .....KEFEQHISN.QENE..............LQKKAED..............................
XlaeNb2  .....QEFEKQILQ.QEEE..............LKRKADE..............................
XtroNb2  .....QEFEKQILQ.QEEE..............LKRKADE..............................
AcalNb   .....MQYIQEHHA.QPGV..............VNPPPDL..............................
BglaNb   ......................................................................
LgigNb   ......................................................................
DmagNb   .....MEFERQ....RQME..............IQRLIDQ.....GMLP.PHPDMM.RNHPRMP....NE
DpulNb   .....MEFEHQ....RQME..............IQRLIDQ.....GMLP.PHPGMM.PNHPRMP....N.
HrobNb   .....AAFEKDLNE.RQHASD...VTPPTTTLSMQQQQHDDQ............................
HaztNb   .....EQYEQQ....RLRE..............LQAAGVYVP..PNAIPLPHPDQQYQVYQGYHP...NQ
TurtNb   .....RQYEAR....RQEL..............LAQQYGY......YMP.PYQPYGNQQGPGQGG..YHQ
PtepNb   .....QEYMKQRAM.QHHG..............DSRMYYDVGNAPQGMP.PHINF..QGHAGAPPPGYPQ
SmimNb   ......................................................................
GoccNb   .....QRFMQM....REHE..............MQMQMAQ.....GYYP.GAPPNL.QYHPGYA......
DsuzNb   .....LEYERR....RQEE..............VQRLIAQ.....GQLP.PHPNMP.QGYYAAP......
DmelNb   .....LEYERR....RQEE..............VQRLIAQ.....GQLP.PHPNMP.QGYYAAP......
AaegNb   .....LEYERR....NQEH..............IQRLIAE.....GKLP.PHPNMP.QGYYPGP....NN
CtelNb   .....EKFEEDYEA.ARRQ..............MQ...................................
OvicNb   .....QEYNRMVNE.RLER...................................................
SkowNb   .....ADFQRQLQE...................MEARRMRNN............................
SpurNb   .....EKYQEELRR.TLEE..............TKAKMAEAG..DNFVR...................VR
NvecNb   .....AEFEKSLEQ.DQHPS.............EQQQRHD..............................
EpalNb   .....AEFEKSLQE.EEKN..............REEHKDS..............................
OfavNb   ......................................................................
HvulNb   .....QEYEKMLSE.NPHGEN.................................................
CbriNb   .....QKFEKEYAA.QQEQ..............LHPQAAQ..............................
CbreNb   .....QKFEQEYAQ.KQEQ..............QHQQAAQ..............................
HconNb   .....QEFEREYAK.QHGQ...................................................
```

```
TcanNb  .....KKFEEEYAK.QQGWGEYAYSTPATTV..AHRDAQQ.............................R
SratNb  .....QVFEEQLAK.ENNWGPDAYKIESTIA...................P...................
ShaeNb  .....RRLASEAQL.KEQN..............HLTHEPD..............................
SmanNb  .....RRLVSEAQS.KEQN...................................................
SjapNb  .....RRLASEAQS.LEKD..............HLTDAPN..............................
AqueNb  .....KQYENEMEE.DDYDYDDEG.........NLIHKPK..............................
SrosNb  .....KEFKDRARK.AGDP..............RQEHVQAR.............................
BdenNb  .................................SYHHKLN..............................
BsalNb  .................................SYHHKLD..............................
CcucNb  .................................NYHHAEV..............................
ArepNb  .................................LYHGKGGI.............................
McirNb  .................................LYHE.................................
MbreNb  .....EKFRERVEA.LKEDRD.GGAVP.....QGVAETAD..............................
DqueNb  .....EEFHSTPET.QTDE..............SYTHPED..............................
FpinNb  .....EEFHSTPET.QTDD..............SYTHPED..............................
FradNb  .....EEFHSTPET.QTDE..............SYVHPED..............................
GfroNb  LTIFPEEFHSSPET.QTDE..............SYTHPED..............................
ShirNb  .....EEYHSTPET.QTDE..............AYNHPED..............................
PostNb  .....EMYHSTPET.QTDE..............SYNHPED..............................
RsolNb  .....EMYHNTPET.QTDE..............SYNHPED..............................
NirrNb  .....KPFYPRCEC..........................L.............................
ScerNb  .....NKFHKD....KDPD......V.......KVVHKED..............................
VpolNb  .....NKYHKD....SDPD......V.......KNVHKAD..............................
TphaNb  .....NKFHRD....TDPD......V.......KNVHKED..............................
CglaNb  .....NKYHKD....KDVN......V.......KNVHKED..............................
LmirNb  .....NKYHKD....KDPY......V.......KTVHKED..............................
EgosNb  .....NTHHQN....NE................KTTHRED..............................
LngbNb  .....GPY..............................................................
McolNb  .....GPY..............................................................
CchtNb  .....GPY..............................................................
SmajNb  .....GPY..............................................................
NnodNb  .....GPYS.............................................................
MproNb  .....GPY..............................................................
PsojNb  ........I.............................................................
PhalNb  ........L.............................................................
PinsNb  .....QPTI.............................................................
AcanNb  ......................................................................
TclaNb  ......................................................................
EsilNb  .....AAAV.............................................................
EguiNb  ......................................................................
DdisNb  .....EYVKKLFAE.NDCE..............LTSAQRFIS...GSVA.................G...
AsubNb  .....EAVKRMFAE.TDAE..............LTSAQRFIS...GASA.................G...
AcasNb  .....ETIKATFGK.KDAD..............ISPHERFIA...GAGA.................G...
LocuNb  .....EQYKKIISS.EGGK..............VRTHERFMA...GSLA.................G...
CcarNb  .....EQYKKLLTK.DGGK..............IQSHERFMA...GSLA.................G...
SansNb  .....EQYKKLLAK.DGGK..............IQSHERFMA...GSLA.................G...
GproNb  .GTGGGEIDGDAAGSRTGS......V.............................................
RgloNb  ......................................................................
AbacNb  ......................................................................
TgonNb  ......................................................................
HhamNb  ......................................................................
BbesNb  ......................................................................
CsuiNb  ......................................................................
CcayNb  ......................................................................
TbruNb  .....QRWQKQEEQWEREEHMHA............SEPENVQRGKRRTPRPTPPTGRREMWRLAGHNTEQ
LbraNb  .....QFFADDVVQ.HNDQRVVVE.........VPREAEDVDTGQRTLGH.........YWR....LTVP
LseyNb  .....DYFESEILQWDND.................GEEDDEGAGGRAVGGPAPPT....HWR....HDAG
PfalNb  LYTLNKMFEKSGKKWQDDNFSFGSYAYPYINIGLHLKNNDTKNPQNKCPFPLPPIKLTDYESSSDECNNN
```

```
                                370                                     380
                                  .                                       .
HsapNb1  .......................LSQ.................ETEA...........LGRSQGRL....
PtroNb1  .......................LSQ.................ETEA...........LGRSQGRL....
PpanNb1  .......................LSQ.................ETEA...........LGRSQGRL....
PabeNb1  .......................LSQ.................ETEA...........LGRSQGRL....
GgorNb1  .......................LSQ.................ETEA...........LGRSQGRL....
CatyNb1  .......................LSQ.................ETEA...........LGRSQGRL....
MmulNb1  .......................LSQ.................ETEA...........LGRSQGRL....
RroxNb1  .......................LSQ.................ETEA...........LGRSQGRL....
CangNb1  .......................LSQ.................ETEA...........LGRSQGRL....
MfasNb1  .......................LSQ.................ETEA...........LGRSQGRL....
MnemNb1  .......................LSQ.................ETEA...........LGRSQGRL....
CcapNb1  .......................LSQ.................ETEA...........LGRSQGRL....
OgarNb1  .......................LSQ.................ETEA...........LGRSQGRL....
PcoqNb1  .......................LSQ.................ETEA...........LGRSQGRL....
GvarNb1  .......................LSQ.................ETEA...........LGRSQGRL....
TchiNb1  .......................LSQ.................ETEA...........LGRSQDRL....
BmutNb1  .......................LSQ.................ETEA...........LGRSQGRL....
MjavNb1  .......................LSQ.................ETEA...........LGRSQGRL....
VpacNb1  .......................LSQ.................ETEA...........LGRSQGRL....
CbacNb1  .......................LSQ.................ETEA...........LGRSQGRL....
VpacNb1  .......................LSQ.................ETEA...........LGRSQGRL....
CferNb1  .......................LSQ.................ETEA...........LGRSQGRL....
HarmNb1  .......................LSQ.................ETEA...........LGRSQGRL....
RsinNb1  .......................LNQ.................ETEA...........LGRSQGRL....
JjacNb1  .......................LSQ.................ETEA...........LGRSQDRL....
DordNb1  .......................LSL.................ETEA...........LGRSQDRL....
NgalNb1  .......................LSQ.................ETEA...........LGRSQDRL....
RnorNb1  .......................LSQ.................ETEA...........LGRSQDRL....
MmusNb1  .......................LSQ.................ETEA...........LGRSQDRL....
MaurNb1  .......................LSQ.................ETEA...........LGRSQDRL....
CasiNb1  .......................LSQ.................ETEA...........LGRSQGHL....
NparNb   .......................LRK.................EHEV...........LQQRKIEL....
XlaeNb1  .......................LRK.................EHEQ...........LQQQQIEL....
XtroNb1  .......................LRK.................EHEQ...........LQQQQIEL....
PmucNb1  .......................LHQ.................EHQE...........LQQRQVEL....
PbivNb1  ......................................................................
GjapNb1  .......................LRR.................QHDE...........LIQRQVQL....
DrerNb1  .......................LRQ.................EQEL...........LKERSKAL....
TrubNb1  .......................LQE.................EQEL...........LRERGRAL....
TnigNb1  .......................LHQ.................EQEL...........LQERGRAL....
IpunNb   .......................LQK.................QRDE...........LERQQEQL....
DrerNb2  .......................LQK.................QRED...........LERQQEQL....
TrubNb2  .......................LQK.................QRDD...........LERQQAEL....
HsapNb2  .......................LQK.................QKEE...........LQRQHDQL....
PpanNb2  .......................LQK.................QKEE...........LQRQHDQL....
GgorNb2  .......................LQK.................QKEE...........LQRQHDQL....
CatyNb2  .......................LQK.................QKEE...........LQRQHDQL....
CangNb2  .......................LQK.................QKEE...........LQRQHDQL....
RroxNb2  .......................LQK.................QKEE...........LQRQHDQL....
PabeNb2  .......................LQK.................QKEE...........LQRQHDQL....
MmulNb2  .......................LQK.................QKEE...........LQRQHDQL....
MfasNb2  .......................LQK.................QKEE...........LQRQHDQL....
CcapNb2  .......................LQK.................QKEE...........LQRQHDQL....
PtroNb2  .......................LQK.................QKEE...........LQRQHDQL....
OgarNb2  .......................LQK.................QKEE...........LQRQHDQL....
PcoqNb2  .......................LQK.................QKEE...........LQRQHDQL....
TchiNb2  .......................LQK.................QKEE...........LQRQHDQL....
MjavNb2  .......................LQK.................QKEE...........LQRQHDQL....
HarmNb2  .......................LQK.................QKEE...........LQRQHDQL....
RsinNb2  .......................LQK.................QKEE...........LQRQHDQL....
DordNb2  .......................LQK.................QREE...........LQRQHDQL....
BmutNb2  .......................LQK.................QKED...........LQRQHDQL....
VpacNb2  .......................LQK.................QKEE...........LQRQHEQL....
CferNb2  .......................LQK.................QKEE...........LQRQHEQL....
CbacNb2  .......................LQK.................QKEE...........LQRQHEQL....
JjacNb2  .......................LQK.................QKEE...........LQRQHDQL....
CasiNb2  .......................LQK.................QKEE...........LQRQHDQL....
NgalNb2  .......................LQK.................QKEE...........LQLQHDHL....
RnorNb2  .......................LQK.................QKEE...........LQRQHDHL....
MmusNb2  .......................LQK.................QKED...........LQRQHDHL....
MaurNb2  .......................LQK.................QKEE...........LQRQHDHL....
PbivNb2  .......................LQR.................KKEE...........LQRQQDFL....
PmucNb2  .......................LQK.................KREE...........LQQQQDFL....
GjapNb2  .......................LQR.................QKEE...........LERQQDQL....
LchaNb   .......................LNK.................QREE...........LQRQQDEL....
XlaeNb2  .......................LYR.................QRED...........LQKQHDYI....
XtroNb2  .......................LYR.................QRED...........LQKQHEYI....
AcalNb   .......................RFQ.................QEQH...........LQQQQ.......
BglaNb   .............................................QH.......................
LgigNb   .............................................TN...........LV..........
DmagNb   VPYGVPP.....QQPYYPSPP..VAQQKMITPEEAMRM.....QQQY.......................
DpulNb   VPYGVAPGQYAGQQPYYPSP...VAQGKPITPEEAIRMQ....QQQY.......................
HrobNb   ............QQQH.......VDQ........QQHVDQ...QQQQH...........VDQQQQHV....
HaztNb   VPMA.HPG....QVPMAH.PGQVMGQVPMAHPGQ.....................VP.............
TurtNb   QP....P.....QQHFGNVPPYQVAQ.................QQGYQ.......KPNGYQAPPQF....
PtepNb   QPQG.YP.....QQGH....PQQFQQ...GYPQQGHPQQ...FQQGH.......................
SmimNb   ......................................................................
GoccNb   .PQGPHPGYQGGHQGAY..PPQGYPQ.QGGHPQQGYPQQGGHPQQGY.........P.QQGGHPQQ....
DsuzNb   .....PPGGVAYQQAP...PP..GGQLHYQQPDQVHAQQ....QQQY...........AQQQQQYA....
DmelNb   .....PPGGVAYQQA....PP..GAQLHYQHPDQVHAQQ....QQQY...........AQQQQQYA....
AaegNb   GPYQVHPNA..............IPQNQQYHPQQGHPNQ..VPQYQA.......................
CtelNb   .......................EEQ.................ERRE...........NERRRDMG....
OvicNb   ......................................................................
SkowNb   .......................VKK................QQQQQ...........LGGEDDKLKFQA
SpurNb   QPQG........QQDAVN.....MNQ................AQAQQIKDGLSAGLPALKEGNDKIRFEG
NvecNb   ..........................................QQQQH.......................
EpalNb   ..........................................QTQEH.......................
OfavNb   ......................................................................
HvulNb   .......................VKQ.................EASH.......................
CbriNb   QP........................................PQQQV.......................
CbreNb   QPAQ......................................AQQQV.......................
HconNb   ......................................................................
```

```
TcanNb  RPAV......................................GEQQA.......................
SratNb  .PTT...............TPLTIVH...............GSQQQQ.......................
ShaeNb  .......................RQQ................TTTQQ.........P.............
SmanNb  .......................RQQ................TTTEQ.........P.............
SjapNb  .......................LHQ.................TTEQ.........P.............
AqueNb  .......................MEN...........HDGVPPEHHE...........VPTEHHEV....
SrosNb  .......................VAE..............ASKRFKD...........KFAKTMKL....
BdenNb  .......................VNQT...........................................
BsalNb  .......................VEPK...........................................
CcucNb  .......................VLH............................................
ArepNb  ......................................................................
McirNb  ......................................................................
MbreNb  .......................LVQAR............LAKERSD...........GFKRLQKM....
DqueNb  .......................IEH...............FAQHEK...........IEREEAAR....
FpinNb  .......................LEH...............FAQHEK...........IEHEEAER....
FradNb  .......................IEH...............FAQHEA...........IERQEAER....
GfroNb  .......................IEH...............FAHHEQ...........IEHQEAER....
ShirNb  .......................IEH...............FASHEK...........IEIQEAER....
PostNb  .......................LEH...............FAQHES...........IERKEAEK....
RsolNb  .......................M..............................................
NirrNb  .......................ICY............................................
ScerNb  .......................IEH............................................
VpolNb  .......................IEH............................................
TphaNb  .......................IEH............................................
CglaNb  .......................IEH............................................
LmirNb  .......................IEH............................................
EgosNb  .......................VEH............................................
LngbNb  ......................................................................
McolNb  ......................................................................
CchtNb  ......................................................................
SmajNb  ......................................................................
NnodNb  ......................................................................
MproNb  ......................................................................
PsojNb  ......................................................................
PhalNb  ......................................................................
PinsNb  ......................................................................
AcanNb  ......................................................................
TclaNb  ......................................................................
EsilNb  ......................................................................
EguiNb  ......................................................................
DdisNb  ......................VVSH..........................TTLFP.LEVVRLRL....
AsubNb  ......................VVSH..........................ASLFP.LEVVRTRL....
AcasNb  ......................VFTH..........................TLSFP.LEVIKTRL....
LocuNb  ......................ATAQ..........................TIIYP.MEVMKTRL....
CcarNb  ......................ATAQ..........................TAIYP.MEVMKTRL....
SansNb  ......................ATAQ..........................TAIYP.MEVMKTRL....
GproNb  ......................................................................
RgloNb  .......................LGQIR............VVGQMDL...........VWRIQERL....
AbacNb  ......................................................................
TgonNb  ......................................................................
HhamNb  ......................................................................
BbesNb  ......................................................................
CsuiNb  ......................................................................
CcayNb  ......................................................................
TbruNb  NPRPLHAHMSKKYQPPVPATHGAVRSNSCLQPPEYSRDLSTVSDARRNV......REGVSQVKHHL....
LbraNb  SPRMAHHGASL..EPQMPASPADVDS.............PAARQAQERV......NAQVKQATAYI....
LseyNb  TPSQGPAAVVR..RPVVPSSPASVQS.............GARAARQRLV......TENIQKRTAFL....
PfalNb  IPISVMKKEYESLQSIQN.....LKKQNT..........KNFKQDKNINH........IKNETDALK...
```

```
HsapNb1  ...........................EAQK.......................................
PtroNb1  ...........................EAQK.......................................
PpanNb1  ...........................EAQK.......................................
PabeNb1  ...........................EAQK.......................................
GgorNb1  ...........................EAQK.......................................
CatyNb1  ...........................EAQK.......................................
MmulNb1  ...........................EAQK.......................................
RroxNb1  ...........................EAQK.......................................
CangNb1  ...........................EAQK.......................................
MfasNb1  ...........................EAQK.......................................
MnemNb1  ...........................EAQK.......................................
CcapNb1  ...........................EAQK.......................................
OgarNb1  ...........................EAQK.......................................
PcoqNb1  ...........................EAQK.......................................
GvarNb1  ...........................EVQK.......................................
TchiNb1  ...........................EAQK.......................................
BmutNb1  ...........................EAQK.......................................
MjavNb1  ...........................EAQK.......................................
VpacNb1  ...........................EAQK.......................................
CbacNb1  ...........................EAQK.......................................
VpacNb1  ...........................EAQK.......................................
CferNb1  ...........................EAQK.......................................
HarmNb1  ...........................EAQK.......................................
RsinNb1  ...........................EAQK.......................................
JjacNb1  ...........................EAQK.......................................
DordNb1  ...........................EAQR.......................................
NgalNb1  ...........................EAQK.......................................
RnorNb1  ...........................EAQK.......................................
MmusNb1  ...........................EAQK.......................................
MaurNb1  ...........................EAQK.......................................
CasiNb1  ...........................EAQK.......................................
NparNb   ...........................DAQK.......................................
XlaeNb1  ...........................NAQK.......................................
XtroNb1  ...........................DAQK.......................................
PmucNb1  ...........................DAQK.......................................
PbivNb1  ......................................................................
GjapNb1  ...........................DAQK.......................................
DrerNb1  ...........................EAQK.......................................
TrubNb1  ...........................EAQR.......................................
TnigNb1  ...........................EAQR.......................................
IpunNb   ...........................NAQK.......................................
DrerNb2  ...........................NAQK.......................................
TrubNb2  ...........................NAQK.......................................
HsapNb2  ...........................EAQK.......................................
PpanNb2  ...........................EAQK.......................................
GgorNb2  ...........................EAQK.......................................
CatyNb2  ...........................EAQK.......................................
CangNb2  ...........................EAQK.......................................
RroxNb2  ...........................EAQK.......................................
PabeNb2  ...........................EAQK.......................................
MmulNb2  ...........................EAQK.......................................
MfasNb2  ...........................EAQK.......................................
CcapNb2  ...........................EAQK.......................................
PtroNb2  ...........................EAQK.......................................
OgarNb2  ...........................EAQK.......................................
PcoqNb2  ...........................EAQK.......................................
TchiNb2  ...........................EAQK.......................................
MjavNb2  ...........................EAQK.......................................
HarmNb2  ...........................EAQK.......................................
RsinNb2  ...........................EAQK.......................................
DordNb2  ...........................EAQK.......................................
BmutNb2  ...........................EAQK.......................................
VpacNb2  ...........................EAQK.......................................
CferNb2  ...........................EAQK.......................................
CbacNb2  ...........................EAQK.......................................
JjacNb2  ...........................EAQK.......................................
CasiNb2  ...........................QAQK.......................................
NgalNb2  ...........................EAQK.......................................
RnorNb2  ...........................EAQK.......................................
MmusNb2  ...........................EAQK.......................................
MaurNb2  ...........................EAQK.......................................
PbivNb2  ...........................QAQK.......................................
PmucNb2  ...........................QAQK.......................................
GjapNb2  ...........................QAQR.......................................
LchaNb   ...........................QAQK.......................................
XlaeNb2  ...........................QAQK.......................................
XtroNb2  ...........................QAQK.......................................
AcalNb   ..............F............QQQQ.......................................
BglaNb   ......................................................................
LgigNb   ......................................................................
DmagNb   .........GQPPQFAPQQHY....GQQPQY.......APQ...QPYYGQPQ...FAQQQ....FVAQPQ
DpulNb   .........AQQPQFAPQQQY....YGQPQF.......AQQ...QQYAGHPQQQFAANPQ....YAAQPQ
HrobNb   ..........................DQQQQ......HVDQ...QQHDDQQQQHDDQQQQH.........
HaztNb   .....MAHPGQVPMAHPGQ.......VMGQV...PMAHPNQMVGHQPVGQVPIHAGQVPPLQRVAQVQPG
TurtNb   .....AVHPGLSQQAHPNQQYNQ...NQNQN.......LNQ...NQNQNQNMQHLPPPP.....PPPQPH
PtepNb   ........PQQFQQGHPQQ.FQQ...GQPQH.......FEQ.......GQPQQFQQGQPQH..FQQGQPQ
SmimNb   ......................................................................
GoccNb   .........GYPQGAHPQQGYPQ..GAHPQQGYPQGAHPQQG..YPQGANPPQGYPQQGYQ...AQGYQA
DsuzNb   .............QQYQQQQY.......GQQ..PVQLHPNQV..YQHAGQIPQ............QQQAV
DmelNb   .............QQYQQQQY.....GNGQQ..PVQLQPNQV..YQHAGQIPQ............QQQPV
AaegNb   ...............HPQQ.......GYPQQ...VNLHPNQV..YQH.HQDPNYQPQTGQQSNVYQHPPQ
CtelNb   ..........................EEVQM........NQG.P..........................
OvicNb   ......................................................................
SkowNb   EAV......AQVAAQQAAQQKSAQELAQQQK...DAQLRNA...QEAARQNQQAAQQQTEQK..EQMKQQ
SpurNb   DTQPVTRHDPPGNIAAANQQPPQPPAGQPQQ...PPAGQPQ...QPPAGQPQQPPAGQPQQP..PAGQPQ
NvecNb   .........................HDQQQQ.......................................
EpalNb   .........................HDQPQE.......................................
OfavNb   ......................................................................
HvulNb   ......................................................................
CbriNb   ...............HPA...........QP...................................IQPV
CbreNb   ...............HPA.........QQQP...................................IQPV
HconNb   ......................................................................
```

```
TcanNb   ............AAVAPV.........QHQM.......................................
SratNb   ......................................................................
ShaeNb   ...........................KVQN.......................................
SmanNb   ...........................NVQN.......................................
SjapNb   ...........................NVQK.......................................
AqueNb   .........................PPEHQE.......................................
SrosNb   ......GTNALKEAIKPFAKVGGNKKPAAQG.......................................
BdenNb   ......................................................................
BsalNb   ......................................................................
CcucNb   ......................................................................
ArepNb   ......................................................................
McirNb   ......................................................................
MbreNb   .........................QADHQE.......................................
DqueNb   .........................EARFQG.......................................
FpinNb   .........................EARFQG.......................................
FradNb   .........................EARFQG.......................................
GfroNb   .........................EAKFQG.......................................
ShirNb   .........................EAKFQG.......................................
PostNb   .........................EAKFQG.......................................
RsolNb   ......................................................................
NirrNb   ......................................................................
ScerNb   ......................................................................
VpolNb   ......................................................................
TphaNb   ......................................................................
CglaNb   ......................................................................
LmirNb   ......................................................................
EgosNb   ......................................................................
LngbNb   ......................................................................
McolNb   ......................................................................
CchtNb   ......................................................................
SmajNb   ......................................................................
NnodNb   ......................................................................
MproNb   ......................................................................
PsojNb   ......................................................................
PhalNb   ......................................................................
PinsNb   ......................................................................
AcanNb   ......................................................................
TclaNb   ......................................................................
EsilNb   ......................................................................
EguiNb   ......................................................................
DdisNb   .............SAEIAGTYNGIKIAISEKSFYRGLGASIT............................
AsubNb   .............SAEPVGTYSGIQTYRAE.GFYRGLTASIL............................
AcasNb   .............AAAPNGTYTGIKIVTKE.GFFRGLTPSLL............................
LocuNb   .............TLRKTGQYSGMKILKRE.GFYKGYVPNIL............................
CcarNb   .............TLRKTGQYSGMKILKKE.GFYKGYVPNIL............................
SansNb   .............TLRKTGQYSGMKILKKE.GFYKGYVPNIL............................
GproNb   ......................................................................
RgloNb   ......................................................................
AbacNb   ......................................................................
TgonNb   ......................................................................
HhamNb   ......................................................................
BbesNb   ......................................................................
CsuiNb   ......................................................................
CcayNb   ......................................................................
TbruNb   ........................QSALFNQG................................SEILPY
LbraNb   ........................CGSRFDDF................................EYVVPI
LseyNb   ........................YGSQFEDF................................RYIVPI
PfalNb   .....CSSYKKEIKSNTYESFTNNINFEDKKSEYNNKSTDQIHKYKNGSYLLELHSKDSENDSNDVHNDV
```

```
                       390       400                                    410
                         .         .                                      .
HsapNb1  .............RELQQAVLHMEQRK..................QQQQQ.QQ..........GHKAPA.
PtroNb1  .............RELQQAVLHMEQRK..................QQQQQ.QQ..........GHKAPA.
PpanNb1  .............RELQQAVLHMEQRK...............QQQQQQQQ.QQ..........GHKAPA.
PabeNb1  .............RELQQAVLHMEQRK...............QQQQQQQQ.QQ..........GHKAPA.
GgorNb1  .............RELQQAVLHMEQRK.................QQQQQQ.QQ..........GHKAPA.
CatyNb1  .............RELQQAVLHMEQRK..............QQQQQQQQQ.QQ..........GHKAPA.
MmulNb1  .............RELQQAVLHMEQRK.............QQQQQQQQQQ.QQ..........GHKDPA.
RroxNb1  .............RELQQAVLHMEQRK.............QQQQQQQQQQ.EQ..........GHKAPA.
CangNb1  .............RELQQAVLHMEQRK................QQQQQQQ.QQ..........GHKAPA.
MfasNb1  .............RELQQAVLHMEQRK.................QQQQQQ.QQ..........GHKAPA.
MnemNb1  .............RELQQAVLHMEQRKQQ..........QQQQQQQQQQQ.QQ..........GHKAPA.
CcapNb1  .............RELQQAVLHMEQRK................QQQQQQQ.QQ..........GHQAPA.
OgarNb1  .............RELQQAVLHMEQRK.....................QQ.QQ..........GHNAPA.
PcoqNb1  .............RELQQAVLHMEQRK.....................QQ.QQ..........GPNAPA.
GvarNb1  .............RELQQAVLHMEQRK...................QQQQ.QQ..........GHSAPA.
TchiNb1  .............RELQQAVLHMEQRK.....................QQ.QQ..........DHKAPA.
BmutNb1  .............RELQQAVLQMEQRK.....................QQ.QQ..........SHNNPA.
MjavNb1  .............RELQQAVLQMEQRK.....................QQ.QQ..........GHNGPA.
VpacNb1  .............RELQQAVLQMEQRK.....................QQ.QQ..........SHNDPA.
CbacNb1  .............RELQQAVLQMEQRK.....................QQ.QQ..........SHNDPA.
VpacNb1  .............RELQQAVLQMEQRK.....................QQ.QQ..........SHNDPA.
CferNb1  .............RELQQAVLQMEQRK.....................QQ.QQ..........SHNDPA.
HarmNb1  .............RELQQAVLQMEQRK.....................QQ.QQ..........DLRDTA.
RsinNb1  .............RELQQAVLQMEQRK.....................QQ.QQ..........SLNDPA.
JjacNb1  .............RELQQAVLHMEQRK.....................QQ.QL..........EGNVPP.
DordNb1  .............RELQQAVLHMEQRK.....................QQ.QQQQE.......GHNAPP.
NgalNb1  .............RELQQAVLQMEQRK.....................QQ.QQ...........NNAPP.
RnorNb1  .............RELQQAVLQMEQRK.....................QQ.QQ..........EQSAPP.
MmusNb1  .............RELQQAVLQMEQRK.....................QQ.LQ..........EQSAPP.
MaurNb1  .............RELQQAVLQMEQRK.....................QQ.QE...........QSAPP.
CasiNb1  .............RELQQAVLHMEQRK.....................QQ.QQQQQQQQQQQQALNAPA.
NparNb   .............KEYQQAVIQMEQHK.....................AQ.QT.........GAAAAPD.
XlaeNb1  .............KEYHQAVLQMEQKK.....................AQ.QT...........GAEAA.
XtroNb1  .............KEYQQAVMQMEQKK.....................AQ.QT...........GAEAA.
PmucNb1  .............KEYQQVVLQMEQRK.....................SQ.QL...........DQASQ.
PbivNb1  ......................................................................
GjapNb1  .............KEYQQAVLQMEQRK.....................SQPQA...........EQGHP.
DrerNb1  .............REYQQAVMEMSQKQ.....................KE.QQ.........ALNKQPP.
TrubNb1  .............REYQQAVLEMSRRQ........................QA..........GDGQPP.
TnigNb1  .............REYQQAVMEMSRRQ........................QA..........GDGQPP.
IpunNb   .............IELQQAVEHMERLK.....................NQ.KV.............EAP.
DrerNb2  .............IELQQAVEHMERIK.....................TQ.KV.............PPP.
TrubNb2  .............VELQQAVEHMERLK.....................SQ.KL.............EPP.
HsapNb2  .............LEYHQVIQQMEQKK.....................L..QG.............IPP.
PpanNb2  .............LEYHQVIQQMEQKK.....................LQ.QG.............IPP.
GgorNb2  .............LEYHQVIQQMEQKK.....................LQ.QG.............IPP.
CatyNb2  .............LEYHQVIQQMEQKK.....................LQ.QG.............IPP.
CangNb2  .............LEYHQVIQQMEQKK.....................LQ.QG.............IPP.
RroxNb2  .............LEYHQVIQQMEQKK.....................LQ.QG.............IPP.
PabeNb2  .............LEYHQVIQQMEQKK.....................LQ.QG.............IPP.
MmulNb2  .............LEYHQVIQQMEQKK.....................LQ.QG.............IPP.
MfasNb2  .............LEYHQVIQQMEQKK.....................LQ.QG.............IPP.
CcapNb2  .............LEYHQVIQQMEQKK.....................LQ.QG.............IPP.
PtroNb2  .............LEYHQVIQQMEQKK.....................LQ.QG.............IPP.
OgarNb2  .............LEYHQVIQQMEQKK.....................L..QA.............IPP.
PcoqNb2  .............LEYHQVIQQMEQKK.....................L..QG.............IPP.
TchiNb2  .............LEYHQVVQQMEQKK.....................LQ.QG.............IPP.
MjavNb2  .............LEYHQVVQQMEQKK.....................LQ.QG.............IPP.
HarmNb2  .............QEYHQVVQQMEQKK.....................LQ.QE.............ISP.
RsinNb2  .............LEYHQVIQQMEQKK.....................IQ.QE.............ISP.
DordNb2  .............QEYHQAVQQLEQKK.....................FQ.QG.............IAP.
BmutNb2  .............LEYHQVVQQMEQKK.....................LQ.Q..............ISP.
VpacNb2  .............LEYHQVVQQMEQKK.....................L..QG.............ISP.
CferNb2  .............LEYHQVVQQMEQKK.....................L..QG.............ISP.
CbacNb2  .............LEYHQVVQQMEQKK.....................L..QG.............ISP.
JjacNb2  .............QEYLQAVQQMEQKK.....................LQ.QG.............LAP.
CasiNb2  .............LEYHQAVQQMEQKK.....................LQ.QG.............ISP.
NgalNb2  .............QEYHQAVQQLEQKK.....................LQ.QG.............VAP.
RnorNb2  .............QEYQQAVQQLEQKK.....................FQ.QG.............IAP.
MmusNb2  .............QEYHQAVQHLEQKK.....................LQ.QG.............IAP.
MaurNb2  .............QEYHQAVQQLEQKK.....................FQ.QG.............IAP.
PbivNb2  .............QELEMAVKQMEKKK.....................LQ.QG.............HPP.
PmucNb2  .............QELEMAVKQMEQKK.....................LQ.QG.............HPP.
GjapNb2  .............QELQQAVQQMELKK.....................LQ.QG.............IPP.
LchaNb   .............EELQQVVQQMEQKK.....................IQ.QG.............LPP.
XlaeNb2  .............QELQQVVQQMEQKK.....................VE.QQ...........QAVPP.
XtroNb2  .............QELQQVVQQMEQKK.....................LE.QQ...........QPVPP.
AcalNb   .............QQFQQQQQQFQQQQ.....................HQ.QQ.........QFQQHPG.
BglaNb   ......................................................................
LgigNb   ......................................................................
DmagNb   HPQFVPQAQQAVQQSFQQPAQLPQQPI...............PQYQIPAQ.QQ........PAIQQQPP.
DpulNb   QPQFA....QPEQQNFQQAAQQQQQ..................NFQQASQ.QQ........QPIQQVP..
HrobNb   .............VDQQQHVDQQQQQQD.......................QRLH...............
HaztNb   YNAQPIS..GHEAMQQQQQAYQQQQQQQVYQQQLQQQQAYQQQQQQQAYQ.QQQQQ.....QAYQQQPNF
TurtNb   YPQVP........QQQQQNGRQYQQPQYQ..............SQSQGNQ.QNPHF.....NSQGNLPP.
PtepNb   QFQQG........QHQQFQQGQHQQFQQG..............QPQQFPQGQPQQFP...QGQPQQFPQ.
SmimNb   ......................................................................
GoccNb   YPNAPPP....PVNNQQPPQINPQIPA................NAQLAHH.QQQ.......QVHQQLPAP
DsuzNb   YQQQPV.......YQQQQPVYQQQQPVQQQQQP....VAQPPAQQQQPVQ.QQQQT.....AQLQQQPVQ
DmelNb   YQNQPV.......YQQQQPV..................................................
AaegNb   YQQQPPQ..QPPQYQQQQPQQPPAHPQQQYQQH....PAQPQPQYQQQQQ.QHAQN.....PAAQQNPQP
CtelNb   .............GGVHPAMAQRGQPM...................................KVDAQPP.
OvicNb   ......................................................................
SkowNb   FDEAHAA...IQQQHHQQQQQQQQHHQ........................QQQQFD.............
SpurNb   QPPASQP...PPPPASDQQQQQQQQQQ........................QQ...........QNQPP.
NvecNb   .............QHHDQQQQHHEQQK...........................................
EpalNb   ..............QHDQAQKDHDKQE........................EK.................
OfavNb   ......................................................................
HvulNb   ......................................................................
CbriNb   NAHAPPP...LPVHNAQPPVQQVEQTLIL...................KI.QQ........PPQQNLPPV
CbreNb   NANPPPP....PVQNAQPPVQQQPPQQPP...................QQ.HQ........QPPQNLPPV
HconNb   ......................................................................
```

```
TcanNb  .............QDIRVDVGSQQQQRPA...................AQ.AA........HSAQSQVPA
SratNb  ...................................................QH.................
ShaeNb  ......................................................................
SmanNb  ......................................................................
SjapNb  ......................................................................
AqueNb  ..........VPTEHHEVPAAHDTKPDTG...............................LPNDHGLPPN
SrosNb  ..........DEGAAANEQQQQQQQQE........................QK.................
BdenNb  ......................................................................
BsalNb  ......................................................................
CcucNb  ......................................................................
ArepNb  ......................................................................
McirNb  ......................................................................
MbreNb  ..........RIKTRRAQKMAHLARKK...........................................
DqueNb  .............ISVEEALKQHEPHPEDNAQ.........QQPVQAPANTDGEAAP..QDPAVDDPA.
FpinNb  .............ISVEDALKQHEPHPEENKA.........QQPVQVPSGTDGETAA..QDPALDDPA.
FradNb  .............ISVDEALAQHEPHD...NA.........QAPVKAPDQQQADTTS.............
GfroNb  .............ISVEEALAQHEPHDEPAAD.........NQP.................EPEVDTPP.
ShirNb  .............ISVEEALAQHEPPPEDPNA.........PAPESITNDSANVVAE..GGDVAQEPV.
PostNb  .............ISVDEVLKSHEEHAPDGAA.........PANPPVPGEGQHVFDEH.PPNDHVPPPA.
RsolNb  ......................................................................
NirrNb  ......................................................................
ScerNb  ...........ELLHHEHEIEHEEEIQRGASR..........ATVITDDELES.................
VpolNb  ...........ELLHHMHEIEHEEKVQRGSSR..........GTVITDDELES.................
TphaNb  ...........ELLHHFHEIEHEEKVQRGASK..........GTVITDDELES.................
CglaNb  ...........ELLHHEHDIEHQEKIQRGAAR..........GTVITDDQLEA.................
LmirNb  ...........ELLHHEHEIEHEEQIQKGASR..........ATVITNDELES.................
EgosNb  ...........DLLHYYHSLEQDE.LKRGATR..........GSYVTDDQLES.................
LngbNb  ......................................................................
McolNb  ......................................................................
CchtNb  ......................................................................
SmajNb  ......................................................................
NnodNb  ......................M...............................................
MproNb  ......................................................................
PsojNb  ......................KETEL...........................................
PhalNb  ......................KETEL...........................................
PinsNb  ......................KETEL...........................................
AcanNb  ......................RSTEL...........................................
TclaNb  ......................ASTEL...........................................
EsilNb  ......................PTTAAL.......................TG.................
EguiNb  ......................................................................
DdisNb  ........ATIPHSGVNMMVYEFLKHK.......VI...............KM.........TGNEFPTA
AsubNb  ........STIPHSGINMLVYESLKHQ.......VI...............NS.........KHGEDPSA
AcasNb  ........STAPHSGIDLTVYEVLKRE.......YT...............KR.........NEGKSPGV
LocuNb  ........GIIPYAGIDLAIYESLKNA.......WL...............SHY.......ATDSPNPGI
CcarNb  ........GIIPYAGIDLAVYETLKNA.......WL...............SKY.......AKDTANPGV
SansNb  ........GIIPYAGIDLAVYETLKNA.......WL...............SRY.......AKDTANPGV
GproNb  ......................................................................
RgloNb  ...........................ES.........................................
AbacNb  .......................Q..............................................
TgonNb  .......................EEVK......CLLSFLARQRERREAVQ.QRKAQR......AKSAPRG
HhamNb  .......................EEVK......CLLSFLARQRERREAVQ.QRKAER......AKVAPRG
BbesNb  .......................EVMK......CFLASLVRQRQRREEAQ.QRKEQR......PQVSTP.
CsuiNb  .......................EEVK......VFLTWLAYQRLRREKKL.NRSERKKAETEPSQCSAR.
CcayNb  .......................DKTR......GLLAFLQSKRK...................PRSEPP.
TbruNb  ATAARRLQGGALPTWMREAVVMLRREEKDRNQKM...AKLIFQKVATNIRTRSILK.R............
LbraNb  ISTATRVTGADVDEALRAAVRQLRAEASERCKRC...VCLLFRRVASTIRTRRLLRGR............
LseyNb  ISTATGVSGGEMSTAMRAAVQDLRKKERMCRKRG...VRLLFNRVANNIRTRTLLRRK............
PfalNb  YNDKYTSNVFYNYNEIENTILNKQHNAEYNKSYFVLLDFTYFIQKELFLKIQAMKHILFNDNNFQSLQDM
```

```
                        420                                 430
                           .                                   .
HsapNb1   ..AHP.....EGQLK..........FH...PD.............TDDVPV...................
PtroNb1   ..AHP.....EGQLK..........FH...PD.............TDDVPV...................
PpanNb1   ..AHP.....EGQLK..........FH...PD.............TDDVPV...................
PabeNb1   ..AHP.....EGQLK..........FH...PD.............TDDVPV...................
GgorNb1   ..AHP.....EGQLK..........FH...PD.............TDDVPV...................
CatyNb1   ..AHP.....EGQLK..........FH...PD.............TDDAPV...................
MmulNb1   ..AHP.....EGQLK..........FH...PD.............TDDAPV...................
RroxNb1   ..AHP.....EGQLK..........FH...PD.............TDDAPV...................
CangNb1   ..AHP.....EGQLK..........FH...PD.............TDDAPV...................
MfasNb1   ..AHP.....EGQLK..........FH...PD.............TDDAPV...................
MnemNb1   ..AHP.....EGQLK..........FH...PD.............TDDAPV...................
CcapNb1   ..ADP.....KGRLK..........FH...PD.............TDDAPV...................
OgarNb1   ..SNP.....EAQLK..........FH...PD.............TDDAPV...................
PcoqNb1   ..SNP.....EGQLK..........FH...PD.............TDDAPV...................
GvarNb1   ..SNS.....EGQLK..........FH...PD.............TDDAPV...................
TchiNb1   ..SNP.....QGQLK..........FH...PD.............TEDAPV...................
BmutNb1   ..PGP.....EGQLK..........FH...PD.............TDDVPV...................
MjavNb1   ..PGP.....EGQLK..........LH...PD.............TDDALV...................
VpacNb1   ..VGP.....EGQLK..........FH...PN.............TDDAPV...................
CbacNb1   ..VGP.....EGQLK..........FH...PN.............TDDAPV...................
VpacNb1   ..VGP.....EGQLK..........FH...PN.............TDDAPV...................
CferNb1   ..VGP.....EGQLK..........FH...PN.............TDDAPV...................
HarmNb1   ..PGP.....EGQLK..........FH...PD.............TDDAPV...................
RsinNb1   ..TGP.....EGQLK..........FH...PN.............TDDAPV...................
JjacNb1   ..SNA.....EGHLQ..........FR...AD.............TDDVPV...................
DordNb1   ..ANP.....EGQLK..........FH...SD.............TDDVPV...................
NgalNb1   ..SNP.....NGQLQ..........FR...PD.............TDDVPV...................
RnorNb1   ..SQP.....DGQLQ..........FR...AD.............TGDAPV...................
MmusNb1   ..SKP.....DGQLQ..........FR...AD.............TDDAPV...................
MaurNb1   ..SKP.....EGQLQ..........FR...AD.............TDDAPV...................
CasiNb1   ..PGP.....EGQLN..........FH...PE.............TDNMPV...................
NparNb    ..VGP.....NGELK..........FQ...AQ.............VPDAAAEKDKIGNNSFQCVKKGDGN
XlaeNb1   ..AGP.....NGELK..........FQ...PE.............APHAESENAK...............
XtroNb1   ..AGP.....NGELK..........FQ...PE.............APHAESEHAK...............
PmucNb1   ..VGP.....GGELK..........FQ...AQ.............PPHA.....................
PbivNb1   ......................................................................
GjapNb1   ..AGP.....GGELK..........FQ...AQ.............PPHA.....................
DrerNb1   ..SGP.....NGELK..........FQ...EGIQK.......LINEDSQAVK..................
TrubNb1   ..VGP.....NGELK..........FE...PP.............GQPMEAK..................
TnigNb1   ..LGP.....NGELK..........FE...PP......................................
IpunNb    ..PEV.....HGR.........................................................
DrerNb2   ..SEI......LEGNAVPESVG.QD.Q...PPVPLEHQP..LPPGHQDVP....................
TrubNb2   ..PEV.....VVEGNALPEIHA.VDNQ...PPMMQEEQ....LQGHHAAPQN..................
HsapNb2   ..SGP.....AGELK..........FE...PH.............I........................
PpanNb2   ..SGP.....AGELK..........FE...PH.............I........................
GgorNb2   ..SGP.....AGELK..........FE...PH.............I........................
CatyNb2   ..SGP.....AGELK..........FE...PH.............I........................
CangNb2   ..SGP.....AGELK..........FE...PH.............I........................
RroxNb2   ..SGP.....AGELK..........FE...PH.............I........................
PabeNb2   ..SGP.....AGELK..........FE...PH.............I........................
MmulNb2   ..SGP.....AGELK..........FE...PH.............I........................
MfasNb2   ..SGP.....AGELK..........FE...PH.............I........................
CcapNb2   ..SGP.....AGELK..........FE...PH.............I........................
PtroNb2   ..SGP.....AGELK..........FE...PH.............I........................
OgarNb2   ..SGP.....TGELK..........FE...PH.............I........................
PcoqNb2   ..SGP.....TGELK..........FQ...PH.............I........................
TchiNb2   ..SGP.....TGDLK..........FQ...PH.............I........................
MjavNb2   ..PGP.....DEESK..........FE...PY.............I........................
HarmNb2   ..PGH.....GGESK..........FE...PL.............I........................
RsinNb2   ..SVP.....GGESK..........FEF.EPL.............I........................
DordNb2   ..SGP.....NGELK..........FE...PH.............V........................
BmutNb2   ..SAP.....GGESK..........I............................................
VpacNb2   ..SVP.....GGESK..........IQ...PH.............I........................
CferNb2   ..SVP.....GGESK..........IQ...SH.............I........................
CbacNb2   ..SVP.....GGESK..........IQ...SR.............M........................
JjacNb2   ..SGP.....AGELK..........FQ...PH.............T........................
CasiNb2   ..SGP.....AGELK..........YQ...PH.............V........................
NgalNb2   ..SGP.....AG.LK..........FQ...PH.............T........................
RnorNb2   ..SGP.....AGELK..........FE...PH.............T........................
MmusNb2   ..SGP.....AGELK..........FE...PH.............T........................
MaurNb2   ..SGP.....AGELK..........FK...PH.............T........................
PbivNb2   ..SGP.....GGELK..........FQ...PS.............VSQPDGS..................
PmucNb2   ..SGP.....GGELK..........FQ...PS.............VSQLDGN..................
GjapNb2   ..SSP.....NGELK..........FQ...PH.............GAQPVGN..................
LchaNb    ..SGP.....NGELK..........FH...PP.............GEQSVQN..................
XlaeNb2   ..AV..........N..........FQ...QP.............GDQT.....................
XtroNb2   ..AVP.......DAN..........FQ...QG.............GDQA.....................
AcalNb    ..................Q.QQ.QFQQ...PPQHQ...................................
BglaNb    ......................................................................
LgigNb    ......................................................................
DmagNb    .......................NQQQ..PPIQQQ...................................
DpulNb    .........................QQ..QPIAQQ...................................
HrobNb    .........................FE...PQ......................................
HaztNb    ..QQP...QAPQQQQIPQQ......QQ..VPQQQQ...................................
TurtNb    ..QAS.MTMNQGAPPRINQ.QS.NFGQ...PSNSF.......GA.............NN...........
PtepNb    ..GQP.QQFPQGQPQQFPQGQPLQFQQG.QPIVQQ.......GQ..........................
SmimNb    ......................................................................
GoccNb    .K......VVAGQPAAPPQ.QQ.QYKQ...PPTQQ.......QQ..........................
DsuzNb    .QQQP..SVQQQQQQTAQ.......QQ...PI......................................
DmelNb    ......................................................................
AaegNb    .NRQP..VANQAQPAVPQQ......NQ...PPPPP...................................
CtelNb    ..PVPTAGHNQGQGR..........QQ...PI...................HH.................
OvicNb    ......................................................................
SkowNb    ..........EAHKAIKQQ.....HHQ...........................................
SpurNb    ..............................PP......................................
NvecNb    ..............................PQ......................................
EpalNb    ......................................................H...............
OfavNb    ......................................................................
HvulNb    ......................................................................
CbriNb    .H.......................HE...PI......................................
CbreNb    .H.......................HE...PI......................................
HconNb    ......................................................................
```

```
TcanNb   .G.......................GE...AA......................................
SratNb   .............................IPE......................................
ShaeNb   ......................................................................
SmanNb   ......................................................................
SjapNb   ......................................................................
AqueNb   DHGLP....................NDHGLPPYDHRLP....ND..........H...............
SrosNb   ......................................................................
BdenNb   ......................................................................
BsalNb   ......................................................................
CcucNb   ......................................................................
ArepNb   ......................................................................
McirNb   ......................................................................
MbreNb   ......................................................................
DqueNb   ..LQP.AKPGPVLPKFTRSKQ..E.DD...PAVRYQNAKA....NGQWGEGDSGY...............
FpinNb   ..LQP.AKPGPVVPKFTRSKQ..EDDD...PAVRYQNAKA....NGQWGEGDHGY...............
FradNb   ..........PSAPRVTRSKP..EIDD...PVARYSSIKA....EGQWGEGDAGY...............
GfroNb   ....P.AS..GGIPKIKRPVPP.EAQD...PAQRFADAKS....DGEWGAGTGGY...............
ShirNb   ....A.AA..SKKIQVQREPSP.DEVD...PAIRFREAKAESEKHGEWGSGDSGY...............
PostNb   ....P.AD..PPQPKYTRAAS....QD...PTTKYKEAERASNQKAEWGQGPDGY...............
RsolNb   ......................................................................
NirrNb   ......................................................................
ScerNb   ......................................................................
VpolNb   ......................................................................
TphaNb   ......................................................................
CglaNb   ......................................................................
LmirNb   ......................................................................
EgosNb   ......................................................................
LngbNb   ......................................................................
McolNb   ......................................................................
CchtNb   ......................................................................
SmajNb   ......................................................................
NnodNb   ......................................................................
MproNb   ......................................................................
PsojNb   ......................................................................
PhalNb   ......................................................................
PinsNb   ......................................................................
AcanNb   ......................................................................
TclaNb   ......................................................................
EsilNb   ..............................A.......................................
EguiNb   ..............................AH......................................
DdisNb   GVCAS.TSSVCGQLV.............GYPFHVVKSRLI..............................
AsubNb   LTCAS.ISSTMGQLV.............SYPIHVIKTRLV..............................
AcasNb   IGCAS.ASSVAGLLA.............CYPLHVAKTRMI..............................
LocuNb   LGCGT.ISSTCGQIA.............SYPLALIRTRMQ..............................
CcarNb   LGCGT.ISSTCGQLA.............SYPLALVRTRMQ..............................
SansNb   LGCGT.ISSTCGQLA.............SYPLALVRTRMQ..............................
GproNb   ......................................................................
RgloNb   ......................................................................
AbacNb   ......................................................................
TgonNb   AECSEEEEQTEGKVKRRKRGR.DSRFAFWRPDVSSRTARQPSRSFTSRSPTSNIVTTSNIATSNGVTSSP
HhamNb   AEGSEEEEHTEGEMKRRKRGR.DSRFAFWRPDVFSRTARQSSGSFTSRSPSSNIV.TSNTAPSNAVTSSP
BbesNb   ARAAPEEENAERPAQKRARAA.DSRFSLVEPDESTLLSYRRAVTRASPGPAAGAGMVSRSTPGASVASPP
CsuiNb   EKSGESKKGEEGSAGRKEQDKDNRRFAKMRPSPAALVTSQHAKLNVSKNP......KSAPIPSCPRAASP
CcayNb   .PSSDEGSGRTGEGGRRRRGC.TPWFAKLRP.....QLQQQQQRVVDPPR..................SA
TbruNb   .RGSAFGGWEKEALRHTQKE..GRVMDMVVN.......................................
LbraNb   .HGEAFHRLEQRQMRVEQHEMYVSLMKLHAP.......................................
LseyNb   .QGEAFTALERRQLQLAKRSNCFTLLQL..P.......................................
PfalNb   NLSQAEVSHNNDRKKAYEQNTDISYFHEKMEKMKLSLIKSNGNNNDDDNNNNNGHIKSN.........DE
```

```
                                                                  440
                                                                    .
HsapNb1  .............................P............................APAGDQKEV.DT
PtroNb1  .............................P............................APAGDQKEV.DT
PpanNb1  .............................P............................APAGDQKEV.DT
PabeNb1  .............................P............................APAGDQKEV.DT
GgorNb1  .............................P............................APAGDQKEV.DT
CatyNb1  .............................P............................APAGDQKEV.DA
MmulNb1  .............................P............................APAGDQKEV.DA
RroxNb1  .............................P............................APAGDQKEV.DA
CangNb1  .............................P............................APAGDQKEV.DA
MfasNb1  .............................P............................APAGDQKEV.DA
MnemNb1  .............................P............................APAGDQKEV.DA
CcapNb1  .............................P............................APAGDQKEV.DT
OgarNb1  .............................P............................APAGDQKDV.EA
PcoqNb1  .............................P............................APAGDQKDV.DA
GvarNb1  .............................P............................APAGDQKDV.DA
TchiNb1  .............................P............................APVGDQRDV.DT
BmutNb1  .............................P............................APAGDQKDV.DA
MjavNb1  .............................P............................APAGDQKDV.DP
VpacNb1  .............................P............................APAGDQKDV.DT
CbacNb1  .............................P............................APAGDQKDV.DT
VpacNb1  .............................P............................APAGDQKDV.DT
CferNb1  .............................P............................APAGDQKDV.DT
HarmNb1  .............................P............................APAGDQKDV.AA
RsinNb1  .............................P............................APAGDQKDV.AA
JjacNb1  .............................P............................APAGDQKDV.DA
DordNb1  .............................P............................APAGDQKGV.EA
NgalNb1  .............................P............................APAGDQKDV.DA
RnorNb1  .............................P............................APAGDQKDV.PA
MmusNb1  .............................P............................APAGDQKDV.PA
MaurNb1  .............................P............................APAGDQKDV.AP
CasiNb1  .............................P............................APADDQKDA.DT
NparNb   LDRGITETVKKLREEHHFQNRARSEFTTTPR........YRNSETCVRGKPGSIRRGDEPSGNSKDF..P
XlaeNb1  ...AVEEQ.....................P............................KPVEQQQKT..E
XtroNb1  ...PVEEQ.....................P............................KLAEQQQKS..D
PmucNb1  ..........................................................APADHASPP..E
PbivNb1  ......................................................................
GjapNb1  ..........................................................APADQTVPP..D
DrerNb1  ..........................................................EP..DQQTG..D
TrubNb1  ..........................................................VPA........E
TnigNb1  ......................................................................
IpunNb   ......................................................................
DrerNb2  .............................PPAAQQHHDELQN....QQALGQE..HQNTPHENQPLP..P
TrubNb2  .............................PHDTPQEHLKQQNQILAQQGFPQS..HQDLPPGNQEAA..Q
HsapNb2  ......................................................................
PpanNb2  ......................................................................
GgorNb2  ......................................................................
CatyNb2  ......................................................................
CangNb2  ......................................................................
RroxNb2  ......................................................................
PabeNb2  ......................................................................
MmulNb2  ......................................................................
MfasNb2  ......................................................................
CcapNb2  ......................................................................
PtroNb2  ......................................................................
OgarNb2  ......................................................................
PcoqNb2  ......................................................................
TchiNb2  ......................................................................
MjavNb2  ......................................................................
HarmNb2  ......................................................................
RsinNb2  ......................................................................
DordNb2  ......................................................................
BmutNb2  ......................................................................
VpacNb2  ......................................................................
CferNb2  ......................................................................
CbacNb2  ......................................................................
JjacNb2  ......................................................................
CasiNb2  ......................................................................
NgalNb2  ......................................................................
RnorNb2  ......................................................................
MmusNb2  ......................................................................
MaurNb2  ......................................................................
PbivNb2  .......................................................PQSHPVGGSQP...L
PmucNb2  .......................................................VQNHPVSSNQP...L
GjapNb2  .......................................................TQNNPEGSRQP...L
LchaNb   .......................................................VDPHPVGGNLP...L
XlaeNb2  ..........................................................IHLGSQQP...L
XtroNb2  ..........................................................VHLGSQQP...L
AcalNb   .......................................QQQF...........................
BglaNb   ......................................................................
LgigNb   ......................................................................
DmagNb   .............................PP........IQQQ...............PPIQQQ....PP
DpulNb   .............................PV........QQQQ...............IPAQQQ....PL
HrobNb   ......................................................................
HaztNb   ............................VPQ........QQQQ...............VPQQQQQV..PQ
TurtNb   ..........................RITPG........QQQVV.....QSSS..NQGIPQGNQSP...P
PtepNb   .............................PQ........QVQQGQPQQFQQTG..NNVQPSNPQA....A
SmimNb   ......................................................................
GoccNb   .............................PR........YQQA...QQPSQGQ..QVQQPAQAQAKAPVT
DsuzNb   .............................P.........QQQV........HN..QSPPPAQSQ......
DmelNb   ......................................................................
AaegNb   ...........................PAPQ........QQQQ...QKPISNN..PAPAPPKPQQ....P
CtelNb   .............................P............................GPPAHHRQQ..A
OvicNb   ......................................................................
SkowNb   ......................................NDANVVV.........MEGGQPLQAQAAQAAQ
SpurNb   ......................................................................
NvecNb   ......................................................................
EpalNb   ......................................................................
OfavNb   ......................................................................
HvulNb   ......................................................................
CbriNb   ......................................................................
CbreNb   ......................................................................
HconNb   ......................................................................
```

```
TcanNb   .....................................SERQQR...........................
SratNb   ......................................................................
ShaeNb   ...........................V..........................................
SmanNb   ...........................V..........................................
SjapNb   ...........................T..........................................
AqueNb   ...........................GLPS........................DHGSPKGEKGLPEHV
SrosNb   ......................................................................
BdenNb   ......................................................................
BsalNb   ......................................................................
CcucNb   ......................................................................
ArepNb   ......................................................................
McirNb   ......................................................................
MbreNb   .....................................................................R
DqueNb   ...........................APPK...........................APS.......DR
FpinNb   ...........................AAPK...........................LPS.......DR
FradNb   ...........................TPPK...........................LPS.......DR
GfroNb   ...........................KPPK...........................SPS.......DR
ShirNb   ...........................KFPS...........................SPS.......EK
PostNb   ...........................KPPV...........................NPA.......DK
RsolNb   ......................................................................
NirrNb   ...........................SLIS........................IMC............
ScerNb   ....................................................................RI
VpolNb   ....................................................................RI
TphaNb   ....................................................................RI
CglaNb   ....................................................................RI
LmirNb   ....................................................................RI
EgosNb   ....................................................................QI
LngbNb   ......................................................................
McolNb   ......................................................................
CchtNb   ......................................................................
SmajNb   ......................................................................
NnodNb   ......................................................................
MproNb   ......................................................................
PsojNb   ......................................................................
PhalNb   ......................................................................
PinsNb   ......................................................................
AcanNb   ......................................................................
TclaNb   ......................................................................
EsilNb   ......................................................................
EguiNb   ......................................................................
DdisNb   .......................................................TQGSSVNQEKYTVPS
AsubNb   .......................................................TGGTKANPEKYSVPN
AcasNb   .......................................................MQSMHGAPQIYSVPS
LocuNb   .......................................................AQASLEGSEQLSLPN
CcarNb   .......................................................AQASMAGSEQVSLPN
SansNb   .......................................................AQASMEGSEPVSLPN
GproNb   ......................................................................
RgloNb   ......................................................................
AbacNb   ......................................................................
TgonNb   TSNNLTSTTL........SSRDASGLPGVPE.....................S..VSLSPAGSRALAVAP
HhamNb   TSNNLTSSTL........SCRGASGLPGVPD.....................S..VSLSPAASRALAVSP
BbesNb   PGAGLP................AHGLQAHPG.....................A..SSGAPATAAAKK...
CsuiNb   ETDGLREDTLP......FAPRDSRHNSARPRDA......DCQV...KRTSSFQ..SPSAPAAVASTSTCP
CcayNb   ADALLQREGL........GSRATENVRQFPR.....................S..SVQSELAALAQQVQP
TbruNb   ........................GFNVVPQ........................QLRGPLETQESPYHF
LbraNb   ........................AAGAVAK........................T...PAAAGKSRYHF
LseyNb   ........................TSSSAAE........................TLTKPAPPSALPFHF
PfalNb   LD................INKI..NLNSKTKNNNYNNIDLQNKIKIHLSISKNIDEQIMSSNNNKWNLSS
```

```
                450                         460
HsapNb1   SEKKLLER...........LPEV...EVPQ....HL.................
PtroNb1   SEKKLLER...........LPEV...EVPQ....HL.................
PpanNb1   SEKKLLER...........LPEV...EVPQ....HL.................
PabeNb1   SEKKLLER...........LPEV...EVPQ....HL.................
GgorNb1   SEKKLLER...........LPEV...EVPQ....HL.................
CatyNb1   SEKKLVER...........LPEV...EVPQ....HL.................
MmulNb1   SEKKLVER...........LPEV...EAPQ....HL.................
RroxNb1   SEKKPAER...........LPEV...EVPQ....HL.................
CangNb1   PEKKLAER...........LPEV...EVPQ....HL.................
MfasNb1   SEKKLVER...........LPEV...EAPQ....HL.................
MnemNb1   SEKKLVER...........LPEV...EVPQ....HL.................
CcapNb1   SEKKLPEQ...........PPEA...EVPQPDSQHL.................
OgarNb1   SEKKILEQ..........QPPE.....VPQ....HL.................
PcoqNb1   SEKKVPEQ..........QPPV.....VPQLDSQQL.................
GvarNb1   SVRKVPEQ...........PPV.....VPQLDSQHL.................
TchiNb1   SEKKSPEN..........LPPE.....GAQPDSQHL.................
BmutNb1   SEKKVPEQ...........TPE.....PPQLDSQHL.................
MjavNb1   LGKKVPEQ...........PPR.....LPQLDSQHL.................
VpacNb1   SEKKVPEQ...........TPG.....LPQGDSQNL.................
CbacNb1   SEKKVPEQ...........TPG.....LPQGDSQNL.................
VpacNb1   SEKKVPEQ...........TPG.....LPQGDSQNL.................
CferNb1   SEKKVPEQ...........TPG.....LPQGDSQNL.................
HarmNb1   SEKKDPEQ...........PHK.....LPQPDSQHL.................
RsinNb1   SEKKDPEQ...........PPK.....LPQLDSQHL.................
JjacNb1   SEKKVPEE...........SPK.....VPQVDSQQL.................
DordNb1   AEQKVPEQ...........SPE.....LPQLDSQHP.................
NgalNb1   SEKKAPGQ...........PPE.....VPQLDSQHL.................
RnorNb1   SEKKVPEQ...........PPV.....LPQLDSQHL.................
MmusNb1   SEKKVPEQ...........PPE.....LPQLDSQHL.................
MaurNb1   SEKKAAEQ...........PPE.....LPQPDSQPL.................
CasiNb1   SEKKVLEQ....................PQLDSQRL.................
NparNb    QFTRSPEK...........EPK.....VSIMLKAAVREGPTSP..........
XlaeNb1   VHPGAPGEAD...HQVGQKEPE.....VPHSQGQDQQAHPGQS..........
XtroNb1   VHPAAPGEAD..GHPAGQKEPE.....VPHSQGQDQQAHPGQP..........
PmucNb1   SHPGAPGAPE...HDAQKDQAE.....PLQNQ.....AHPEVAVQ........
PbivNb1   ...........................PTQ.......................
GjapNb1   SHPGAPE......HAAPQNQAE.....PPQTQ.....QHPEVEIQ........
DrerNb1   VQNNLPAE...........PPQ....NLPQHTS....................
TrubNb1   TPNNLPAE...........PPQ....NLPVHT.....................
TnigNb1   .....................................................
IpunNb    .....................................................
DrerNb2   GHNNPPED...........PPL.....VPLDHNIVP.................
TrubNb2   GQHHLP...............................................
HsapNb2   .....................................................
PpanNb2   .....................................................
GgorNb2   .....................................................
CatyNb2   .....................................................
CangNb2   .....................................................
RroxNb2   .....................................................
PabeNb2   .....................................................
MmulNb2   .....................................................
MfasNb2   .....................................................
CcapNb2   .....................................................
PtroNb2   .....................................................
OgarNb2   .....................................................
PcoqNb2   .....................................................
TchiNb2   .....................................................
MjavNb2   .....................................................
HarmNb2   .....................................................
RsinNb2   .....................................................
DordNb2   .....................................................
BmutNb2   .....................................................
VpacNb2   .....................................................
CferNb2   .....................................................
CbacNb2   .....................................................
JjacNb2   .....................................................
CasiNb2   .....................................................
NgalNb2   .....................................................
RnorNb2   .....................................................
MmusNb2   .....................................................
MaurNb2   .....................................................
PbivNb2   PPGNIPVQEA.............AA.RSDHVQSHP..................
PmucNb2   APGNIPVQEA.............AA.RSDHAQSHP..................
GjapNb2   PPGTIPVQEA.............AA.GSDHVQTLT..................
LchaNb    TGGHMPVQEV.........QKEGAT.GAEQSNLHP..................
XlaeNb2   PPGHLPET..................GVPQSHPVP..................
XtroNb2   PPGHLPEA..................GVPQSHPVP..................
AcalNb    ..QQPPQQQQ...........................................
BglaNb    .....................................................
LgigNb    .....................................................
DmagNb    IQQQPPIQQQPPIQQQ....P.....PIQQ.......................
DpulNb    VQPQVPVQQQAPIQSQPVALP.....PAPQQIPAAKTV...............
HrobNb    ...QQPQHDQ..........PKGVI............................
HaztNb    QQQQVPQQQQ...QQQ..........QVPQQ......................
TurtNb    LMNQNPNSNM..........NQGSINSIPQQNSGSFNSQPNHKS.........
PtepNb    TETQKPSDQQ.......AHKTEG...NTAQPGAPSVAATKQQT..........
SmimNb    .....................................................
GoccNb    QGNNAPVVQQ................PSQQARQ....................
DsuzNb    ...QVPVQQQ..................QKQHQETANQANQPH..........
DmelNb    .....................................................
AaegNb    AQQNAPNSQP..........PAG...PIQQQQQPAPVQSQQN...........
CtelNb    TNEHDPTHHE.....Q..........SNPQEQSDSLKYAHGRNAEQGGAPSAN
OvicNb    .....................................................
SkowNb    VNENKPQQ.............................................
SpurNb    ...KQPAQQP...........................................
NvecNb    .....................................................
EpalNb    .....................................................
OfavNb    .....................................................
HvulNb    .....................................................
CbriNb    .............................QDHTKDPTYGI.............
CbreNb    .............................QDHTKDPTYGV.............
HconNb    .....................................................
```

```
TcanNb   ..........................LAPQGNTLDKTYGV.............
SratNb   .............................QNHKIDPVMGI.............
ShaeNb   .....................................................
SmanNb   .....................................................
SjapNb   .....................................................
AqueNb   PPQDVPHDVPV.GGANHNENPKAPIVETKQDDTAPPPHAASDDEKKL......
SrosNb   .....................................................
BdenNb   .....................................................
BsalNb   .....................................................
CcucNb   .....................................................
ArepNb   .....................................................
McirNb   .....................................................
MbreNb   AAQEPPAE.............................................
DqueNb   LKKNVPYK.............................YKFRRNWGDF......
FpinNb   LKKNVPYK.............................YKFRRTWGDF......
FradNb   LKKNLPYK.............................YKFRRTWGDF......
GfroNb   LRKNLPYKDRI.........................QYKFRRTWGDF......
ShirNb   MRKNLPYK.............................YKFRRSWGDF......
PostNb   LRYVVSPKERM.........................QYKFRRNWGDF......
RsolNb   .....................................................
NirrNb   .....................................................
ScerNb   ELKNIPEK.............................FKN....GIF......
VpolNb   DLKNIPNK.............................FKN....GFF......
TphaNb   DIKNIPKK.............................FKN....GFF......
CglaNb   ELKNIPSR.............................FKN....GYF......
LmirNb   DLSKVPSK.............................YRL....GLS......
EgosNb   MKQKIPSK.............................YLT....GRL......
LngbNb   .....................................................
McolNb   .....................................................
CchtNb   .....................................................
SmajNb   .....................................................
NnodNb   .....................................................
MproNb   .....................................................
PsojNb   .....................................................
PhalNb   .....................................................
PinsNb   .....................................................
AcanNb   .....................................................
TclaNb   .....................................................
EsilNb   .....................................................
EguiNb   .....................................................
DdisNb   FMKSIPSHSITFFKKAF.........DVNKEKKHH..................
AsubNb   FMKSIPSHGITFLKNAF.........NIKKHEKHH..................
AcasNb   ILKSVPSHCITFLKKQF.........GVEKHSKH...................
LocuNb   FMKVIPAVSISYMKTGL.........GISK.......................
CcarNb   FMKVIPAVSISYMRSGL.........GISK.......................
SansNb   FMKVIPAVSISYMRSGL.........GISK.......................
GproNb   .....................................................
RgloNb   .....................................................
AbacNb   .....................................................
TgonNb   RDKEHPGSLSTAGGRLSPAPP....VNVPSMQSPAPGEALQSVPDGNARPM..
HhamNb   RDKEQPGSLSTAGGGLSPAPP....VNVPSMQSPAPGEALQSVPDGNVRPM..
BbesNb   .....PAASSQG....SGAPPA...AKVPSRSAETPHQNVSSVLGAKAEPAAC
CsuiNb   ..AAHPSSTDQC....RGVAPASCQVKVPSRT.....................
CcayNb   ..RQQPAISSGA..LVQAAKERQEQ....SLSGTQPQI...............
TbruNb   RDARIPSKDK...........................................
LbraNb   DKVRFDDK.............................................
LseyNb   DEAKFNTES............................................
PfalNb   SEHNMNNKNDKF.......CPSCSNKHVQKKNK....................
```
